# *Anopheles* mosquitoes revealed new principles of 3D genome organization in insects

**DOI:** 10.1101/2020.05.26.114017

**Authors:** Varvara Lukyanchikova, Miroslav Nuriddinov, Polina Belokopytova, Jiangtao Liang, Maarten J.M.F. Reijnders, Livio Ruzzante, Robert M. Waterhouse, Zhijian Tu, Igor V. Sharakhov, Veniamin Fishman

**Author notes:** co-first authors (Varvara Lukyanchikova, Miroslav Nuriddinov). **correspondence to I.V.S. and V.F.**.

## Abstract

Chromosomes are hierarchically folded within cell nuclei into territories, domains and subdomains, but the functional importance and evolutionary dynamics of these hierarchies are poorly defined. Here, we comprehensively profiled genome organizations of five *Anopheles* mosquito species and showed how different levels of chromatin architecture influence contacts between genomic loci. Patterns observed on Hi-C maps are associated with known cytological structures, epigenetic profiles, and gene expression levels. At the level of individual loci, we identified specific, extremely long-ranged looping interactions, conserved for ~100 million years. We showed that the mechanisms underlying these looping contacts differ from previously described Polycomb-dependent interactions and clustering of active chromatin.

## Introduction

Three-dimensional genome organization has recently been recognized as a complex and dynamic mechanism of gene regulation. Understanding of these features has been advanced by the development of chromosome conformation capture (3C) methods, which enable genome-wide chromatin contacts to be studied at high resolution^1–3^. In addition, data obtained using 3C-methods help to generate high quality, chromosome-level genome assemblies, allowing comprehensive analysis of genome evolution^4^.

Comparative studies performed on multiple vertebrate species revealed that genome architecture is evolutionarily conserved and could be explained by dynamic interplay between processes of cohesin-mediated loop extrusion and chromatin compartmentalization^5–8^. In insects, comprehensive analyses^9–13^ and cross-species comparisons^12,14^ of genome architecture have to date focused only on *Drosophila* species. These studies suggested that, in contrast to mammals, the process of loop extrusion does not define the structure of chromatin contacts14. Instead, separation of active and repressed chromatin plays an essential role in formation and interaction of topologically associated domains (TADs)^15,16^, which are basic units of chromatin organization in *Drosophila*.

Recently, Ghavi-Helm et al.^17^ suggested that disrupting TADs does not influence coordinated gene expression, based on the analysis of *Drosophila* lines with highly rearranged genomes. In contrast, Renschler et al.^14^ used three distantly related *Drosophila* species to show that while chromosomal rearrangements might shuffle the positions of entire TADs, they are preferentially maintained as intact units. Thus, the roles that 3-dimensional chromatin interactions play in the function and evolution of insect genomes remain unclear.

To address these apparently conflicting observations, we characterized the chromosomal-level genome architectures of five *Anopheles* mosquito species using a Hi-C approach. Comparative analyses of multiple mosquito genomes^18^ previously revealed a high rate of chromosomal rearrangements, especially on the X chromosome, making them an attractive model for studying interconnections between structural variations, chromosome evolution and genome architecture. Malaria has a devastating global impact on public health and welfare and *Anopheles* mosquitoes are exclusive vectors of human malaria parasites.

The Hi-C data allowed us to improve existing genome assemblies of three mosquito species and generate new chromosome-length assemblies for others. Our analysis of TADs and genomic compartmentalization, supplemented by improved algorithms for compartment identification, demonstrated conservation of principles controlling chromatin organization in *Anopheles* species and other insects.

We found specific looping interactions, sometimes spanning several dozens of megabases (Mb) in all studied *Anopheles* species, and showed that these interactions are evolutionarily conserved over ~100 million years. We generated RNA-seq and ChIP-seq data to show that these loops cannot be explained using known molecular mechanisms and represent a specific type of chromatin interactions.

Aggregating long-range chromatin interactions, we found that there is a decrease of contact probabilities beyond a certain genomic distance. Performing broad evolutionary comparison between *Anopheles* species, other insects, and vertebrates, we showed that this limiting genomic distance is taxon-dependent and suggested a mechanistic explanation of this phenomenon.

## Results

### Hi-C-guided assembly of five Anopheles mosquito genomes

In the Hi-C experiment we used 15-18h embryos of mosquito species from three different subgenera of the *Anopheles* genus: Cellia (*An. coluzzii*, *An. merus*, *An. stephensi*), Nyssorhynchus (*An. albimanus*) and Anopheles (*An. atroparvus*) (Fig. 1, A-C). In addition, we sequenced Hi-C libraries from an adult *An. merus* mosquito. Based on our analysis, phylogenetic relationships of the selected species represent a broad range of evolutionary distances, from 0.5 million years (MY) between closely related *An. coluzzii* and *An. merus* species^19^ to 100 MY separating the most distant lineages such as *An. coluzzii* and *An. albimanus*^18^ (Fig. 1, A).

**Fig. 1.**
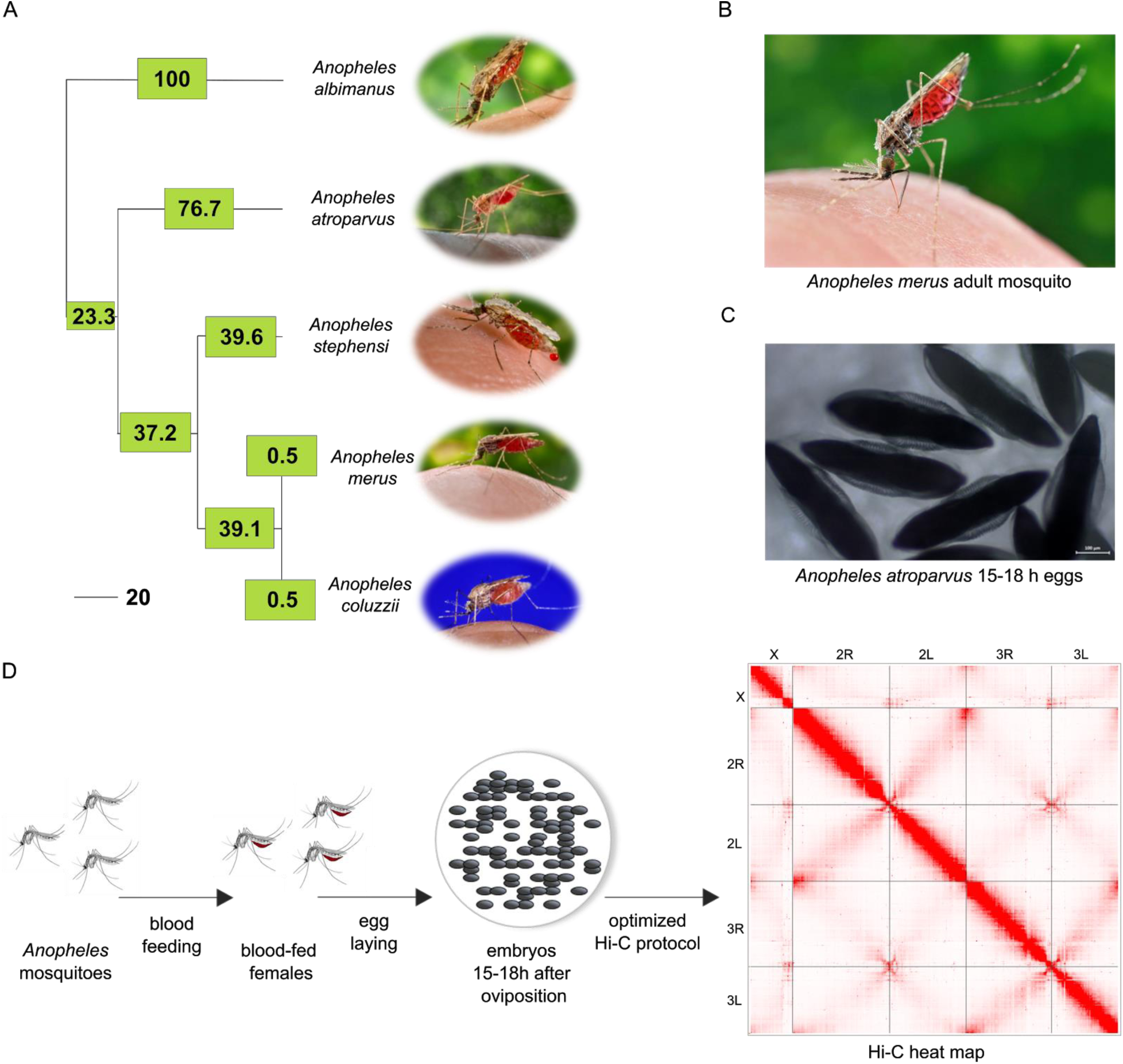
Selected species and Hi-C experimental setup. A) A time-calibrated phylogenetic tree shows estimated evolutionary distances among the selected Anopheles species; numbers in boxes show divergence times in millions of years (MY) for each branch; the scale bar represents 20 MY; B) Anopheles merus adult female mosquito; Adult mosquito illustrations were taken from VectorBase repository C) Anopheles atroparvus 15-18h embryo developmental stage; D) The experimental design of the embryo Hi-C experiments.

After sequencing the libraries and merging biological replicas, we obtained 60-194 million of unique alignable reads for each species (Supplementary Table 1). Library statistics show high quality of the obtained data (Supplementary Table 1).

Genomes of *Anopheles* species have been challenging to assemble to chromosomal levels, mainly due to the presence of highly repetitive DNA clusters that regular Illumina sequencing and assembly cannot successfully resolve^18^. Chromosome-length assemblies were already available for two species (*An. albimanus*, *An. atroparvus*), whereas for *An. coluzzii*, *An. merus* and *An. stephensi* there were only scaffold-level assemblies with N50s of 3.5-Mb^20^, 2.7-Mb^18^, and 1.6-Mb^21^ correspondingly. While evolutionary superscaffolding and chromosomal anchoring improved these assemblies, they did not reach a full chromosomal level^22^.

We employed a 3D-DNA pipeline^4^ to *de novo* assemble *An. coluzzii*, *An. merus*, and *An. stephensi* genomes using our Hi-C datasets. We reassembled chromosomes of *An. albimanus* and *An. atroparvus* using available chromosomal assemblies as drafts. For all species, five large scaffolds corresponding to the number of chromosomal arms (X, 2R, 2L, 3R, 3L) were identified (Table 1). Multiple misassemblies and several chromosome rearrangements were detected and fixed manually. Available physical maps of the genomes were used to verify these corrections^23–27^. For *An. coluzzii Mopti* we used recently published PacBio contigs from a single *An. coluzzii Ngousso* mosquito^20^, characterized by N50 of 3.5-Mb, because our Hi-C data revealed multiple errors in scaffolds of the existing *An. gambiae* PEST assembly. This indeed resulted in a more accurate assembly which was used in further analysis (Table 1). For *An. merus* we performed PacBio sequencing using whole genomic DNA extracted from 100 adult males, which produced reads with an average read length N50 of 2.7-Mb (Supplementary Table 2).

**Table 1.** Assembly statistics before (reference) and after (de novo) Hi-C assembly. *The column “break-points” displays numbers of misjoins corrected manually by redirecting or repositioning scaffolds.

| Species | Total scaffold length | Reference genome |  |  | De novo assembly |  |  |  |
| --- | --- | --- | --- | --- | --- | --- | --- | --- |
|  |  | length of chromosomes | length of gaps | Scaffold N50 | length of chromosomes | length of gaps | Scaffold N50 | break points* |
| *An. albimanus* | 170,336,140 | 167,376,415 | 3,003,101 | 9,735,467 | 169,448,125 | 10,000 | 5,915,397 | 23 |
| *An. atroparvus* | 224,290,125 | 200,912,972 | 1,005,100 | 9,206,694 | 216,747,066 | 116,200 | 688,000 | 465 |
| *An. coluzzii* | 251,414,185 | - | - | 3,468,756 | 231,617,039 | 13,700 | 3,835,000 | 77 |
| *An. merus* | 300,704,392 | - | - | 2,729,089 | 234,366,716 | 27,100 | 3,903,996 | 17 |
| *An. stephensi* | 221,324,304 | - | - | 1,631,802 | 196,394,606 | 62,100 | 980,283 | 144 |

Pairwise alignments of the resulting assemblies showed that the lengths of alignment blocks and percentages of alignable nucleotides correlate with evolutionary distances among species, ranging from 19% to 93% alignable nucleotides (Fig. 2, A). For all species, five large scaffolds corresponding to the number of chromosomal arms (X, 2R, 2L, 3R, 3L) were identified. Malaria mosquitoes’ chromosomal complement consists of a 5-arm. In *An. gambiae*, arms are denoted as chromosomal elements 1 (X), 2+3 (2R+2L), 4+5 (3R+3L). The correspondence of other species chromosomal arms is as follows: *An. merus* 1, 2+3, 4+5; *An. stephensi* 1, 2+5, 3+4; *An. atroparvus* 1, 4+3, 2+5; *An. albimanus* 1, 2+4, 5+3^18^. Therefore, *An. stephensi* and *An. atroparvus* have the same arm association, although different arm names for the same elements. We found that the vast majority of the rearrangements occur within the same chromosomal arm, and even for most evolutionary distant species, inter-chromosomal translocations are extremely rare (Fig. 2, B). Comparing individual chromosomes, we found that for all species alignment blocks on the X chromosome were smaller than on autosomes (Fig. 2, C), in agreement with previously shown elevated gene shuffling on the X chromosome^18^. Overall, the Hi-C data allowed us to improve existing genome assemblies for two *Anopheles* species and generate chromosome-level assemblies for three species *de novo*.

**Fig. 2.**
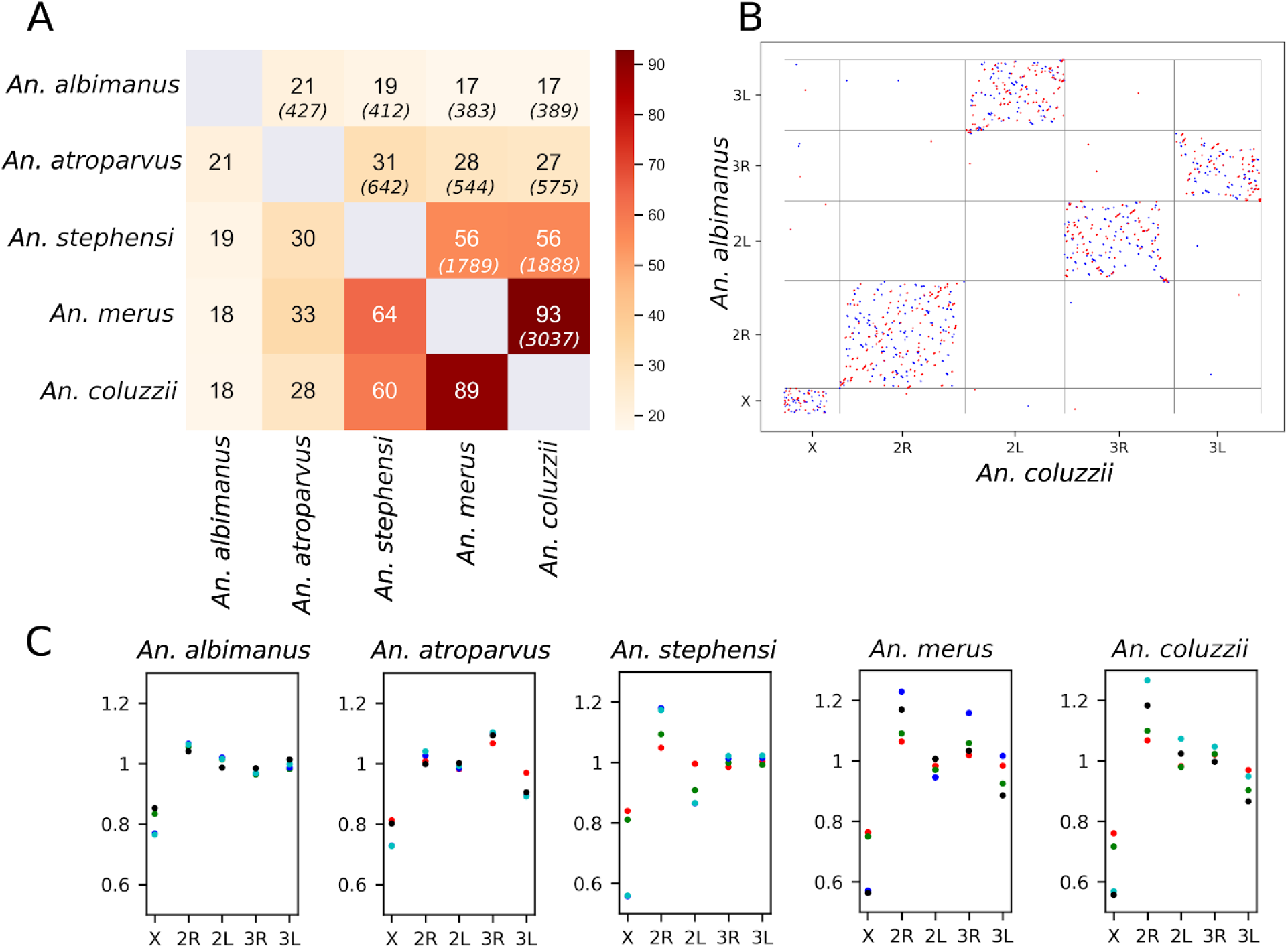
Comparison of alignability and rearrangements among five chromosome-level *Anopheles* genomes. A. Heatmap representing the percentage of alignable nucleotides and mean lengths of alignment blocks (numbers shown in italic) for each pair of genomes. B. Synteny dot-plot showing results of pairwise alignments between An. coluzzii and An. albimanus. C. Comparison of alignment block lengths for each chromosome arm. Y-axes represent the ratio of alignment block length on the specific chromosome (depicted on X-axes) to the genome-wide average. Titles of plots indicate alignment reference and colors of dots correspond to query species: red - An.albimanus, green - An.atroparvus, indigo - An.coluzzii, azure - An.merus, black - An.stephensi. Note that only alignment blocks longer than 1000 base pairs were used to filter out blocks representing individual exons.

### Hi-C data identifies polymorphic inversions

Our chromosome-length Hi-C maps of five *Anopheles* species revealed several regions characterized by “butterfly” contact patterns, associated previously^28^ with balanced inversions. The most prominent example of such contacts was found on chromosome 2R of *An. stephensi* (Fig. 3, A), where the Hi-C map suggests an inversion of a large (~16 Mb) chromosomal region.

**Fig 3.**
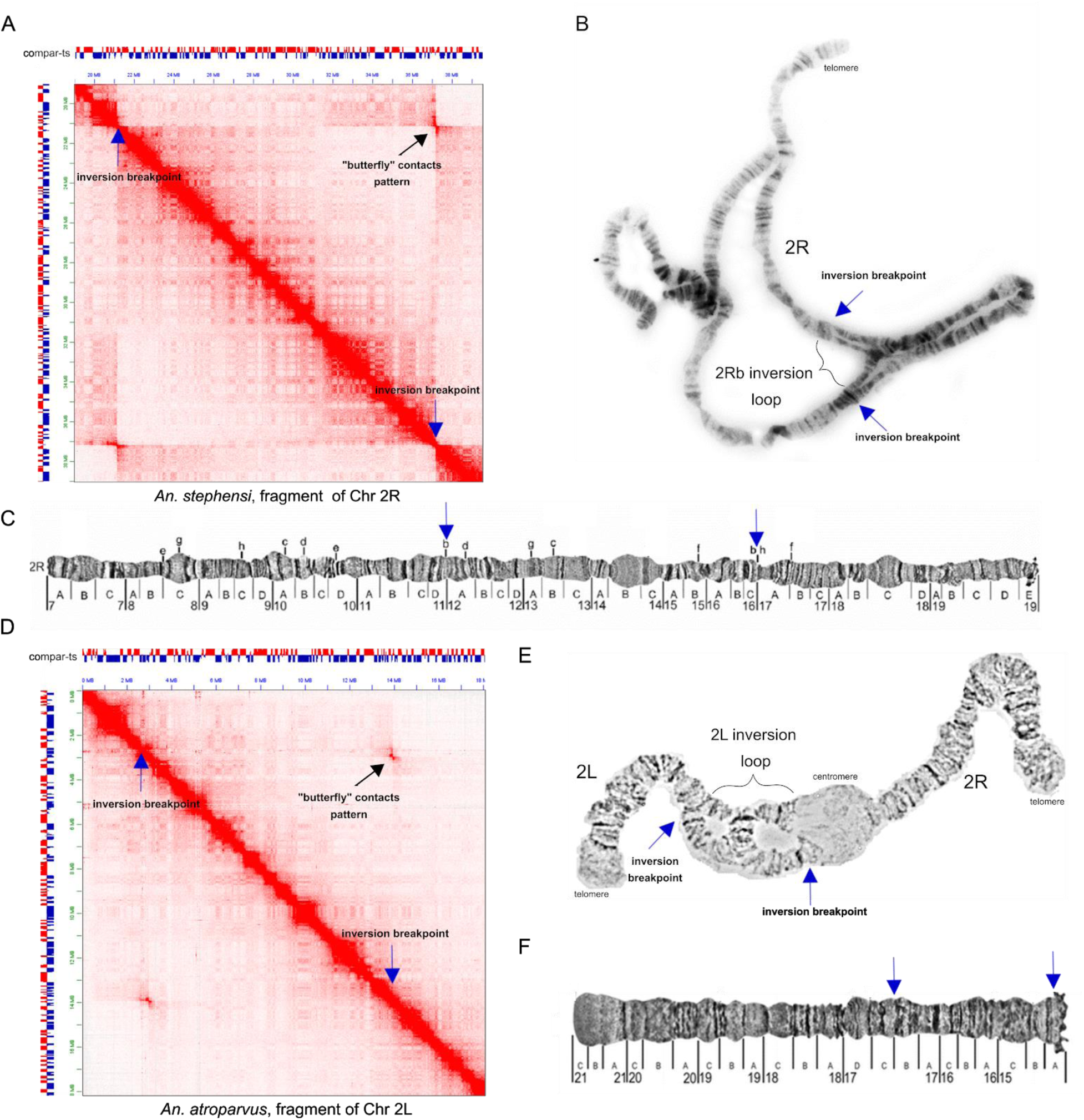
Polymorphic inversions identified using Hi-C data. A. Region of the An. stephensi Hi-C heat map with the 2Rb-polymorphic inversion manifested as a clear “butterfly” pattern; B. Light microscope image of An. stephensi polytene chromosomes showing the 2Rb-inversion loop; C. The physical map of An. stephensi chromosome 2R, published previously in Sharakhova et al.^32^; D. Region of the An. atroparvus Hi-C heat map with the 2L-polymorphic inversion; E. Light microscope image of An. atroparvus polytene chromosome 2 showing the 2L-inversion loop; F. Physical map of An. atroparvus chromosome 2L, published previously in Artemov et al.^27^. Inversion breakpoints are shown with blue arrows in all panels.

We observed both off-diagonal long-range interactions, as well as interactions near the diagonal, suggesting that both variants with inverted and standard arrangements were present in the population. A paracentric inversion was previously demonstrated for the Indian wild-type laboratory strain of *An. stephensi* and known as polymorphic inversion 2Rb^29–31^. The boundaries of this inversion on chromosome 2R were in agreement with previous cytological data^25^.

Further examination of *Anopheles* Hi-C contact maps allowed us to identify a total of four inversions, ranging from 2.8 Mb to 16 Mb (the 2Rb inversion) (Supplementary Table 3, Supplementary Figure 1).

### 3D-chromatin structure of Anopheles genome revealed by Hi-C

The Hi-C contact maps clearly delineated five chromosomal elements for all experimental genomes - X, 2R, 2L, 3R, 3L (Fig. 4, A shows an example of chromosomal elements of *An. albimanus*). Strong centromere-centromere clustering was detected as well as inter- and intra-chromosomal telomere-telomere interactions. These patterns suggest a Rabl-like configuration^33–35^ in *Anopheles* genomes where centromeres and telomeres form clusters within the nuclear space (Fig. 4, B). Additionally, we observed another manifestation of the Rabl-like configuration, represented by interactions between chromosomal arms as perpendicular to the main diagonal “wings” pattern on the Hi-C contact map (Fig. 4, A). Quantitative analysis of contact frequencies confirmed an increase of interactions between loci from different chromosome arms located equidistantly from the centromeres (Supplementary Fig. 2). This increase of interactions was more pronounced in embryonic tissues (1.49-2.45 times) than in adult mosquito data (1.19-1.25 times), suggesting that the Rabl-like configuration might be a feature of some but not all cell types. 3D-FISH experiments on *An. stephensi* ovarian tissue confirmed centromeric clustering in follicular cells and the lack of centromeric interactions in nurse cells within polytene chromosomes (Fig. 4, C).

**Fig. 4.**
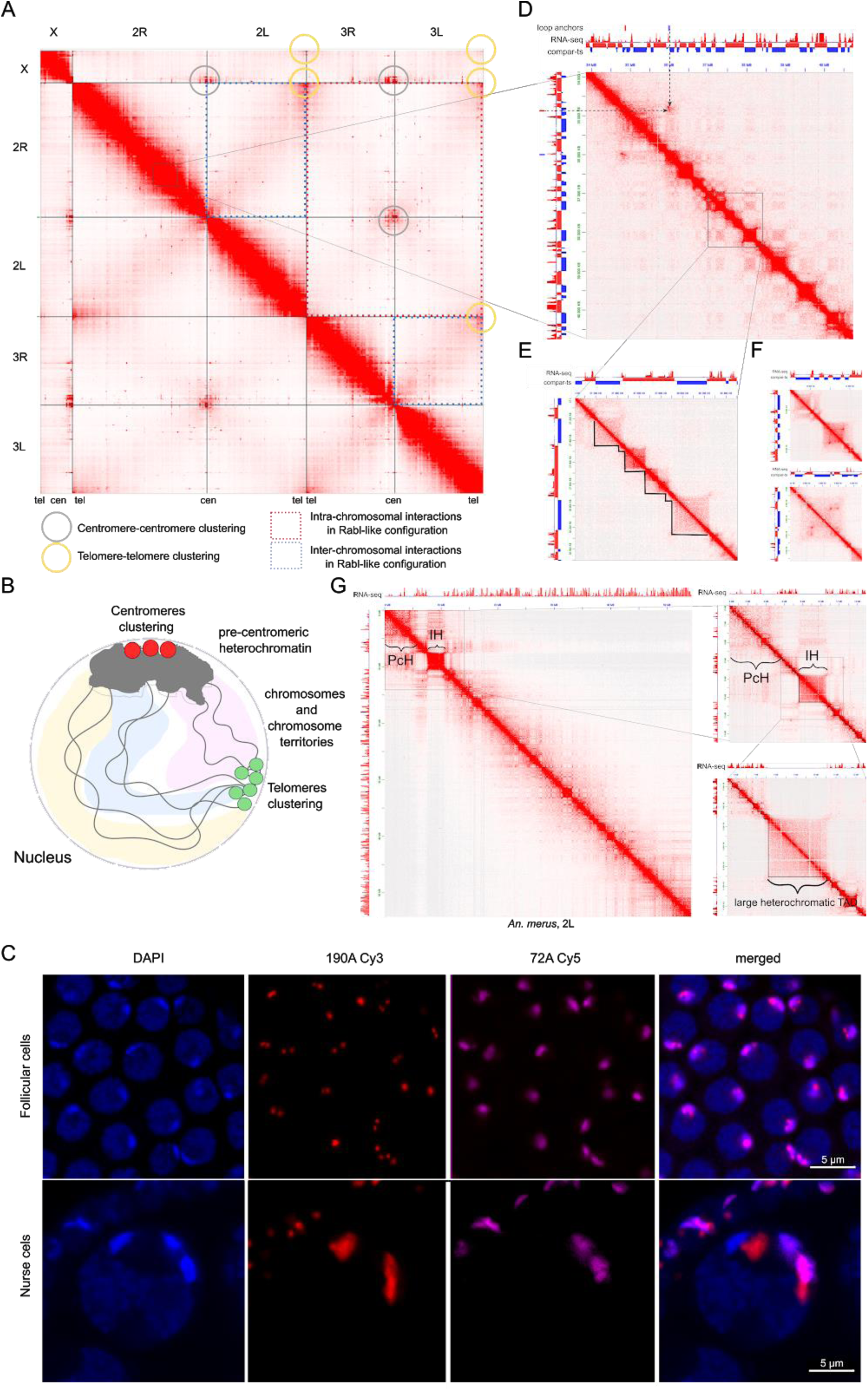
3D-chromatin structure of *Anopheles* mosquito genomes. Hi-C contact map for An. albimanus; grey and yellow circles correspond to contacts between telomeres and centromeres, respectively; B) Schematic representation of Rabl-like configuration; C) FISH showing position of centromeres: 190A Cy3 probe - autosomal centromere, 72A Cy5 probe - Chr X centromere; D-F) Zoomed-in Hi-C map regions showing long-distance loops between TADs (D, indicated by arrows), compartments (D), TADs (E), loops within TADs (F), and heterochromatic blocks either located near centromeres (PcH) or interspersed throughout the euchromatin (IH) (G).

Magnified views of the contact maps revealed large insulated blocks of chromatin located either near centromeres or interspersed throughout chromosomal arms (Fig. 4, G; Supplementary Fig. 3, A-F). The sizes and locations of these blocks were in agreement with the positions of pre-centromeric and intercalary heterochromatic blocks on standard cytological maps (Supplementary Fig. 4). Outside of the heterochromatic blocks, we observed a checkerboard-like pattern of long-range interactions, which was previously shown to represent spatial compartments of the chromatin (Fig. 4, D). Some long-range contacts were extremely pronounced, corresponding to looping interactions between loci located several Mb away from each other (Fig. 4, D; Supplementary Table 4). At the highest resolution (1-25 kb) we observed triangles formed above diagonal, corresponding to the TADs (Fig. 4, E), and chromatin loops located within some of these TADs (Fig. 4, F).

We next aimed to quantitatively characterize TADs, compartments, and loops identified in *Anopheles* genomes.

### Anopheles genomes are partitioned into compartments

Inspecting genomic interactions at various resolutions, we observed prominent signs of genome compartmentalization, manifested as plaid-patterns of Hi-C contacts (Fig. 4). However, when we employed principal component analysis, which is widely used to identify spatial chromatin compartments^1,36^, we found that for the majority of chromosomes the first principal component (PC1) of the Hi-C matrix does not reflect observed plaid-patterns (Fig. 5, A-B; Supplementary Fig. 5. A-M). In most cases, chromosomal arms were split into two or three large, contiguous regions characterized by similar PC1 values (see examples in Fig. 5 and Supplementary Fig. 5 A-I). Thus, PC1 values reflect the position of the locus along the centromere-telomere axis rather than the plaid-pattern of contacts.

**Fig. 5.**
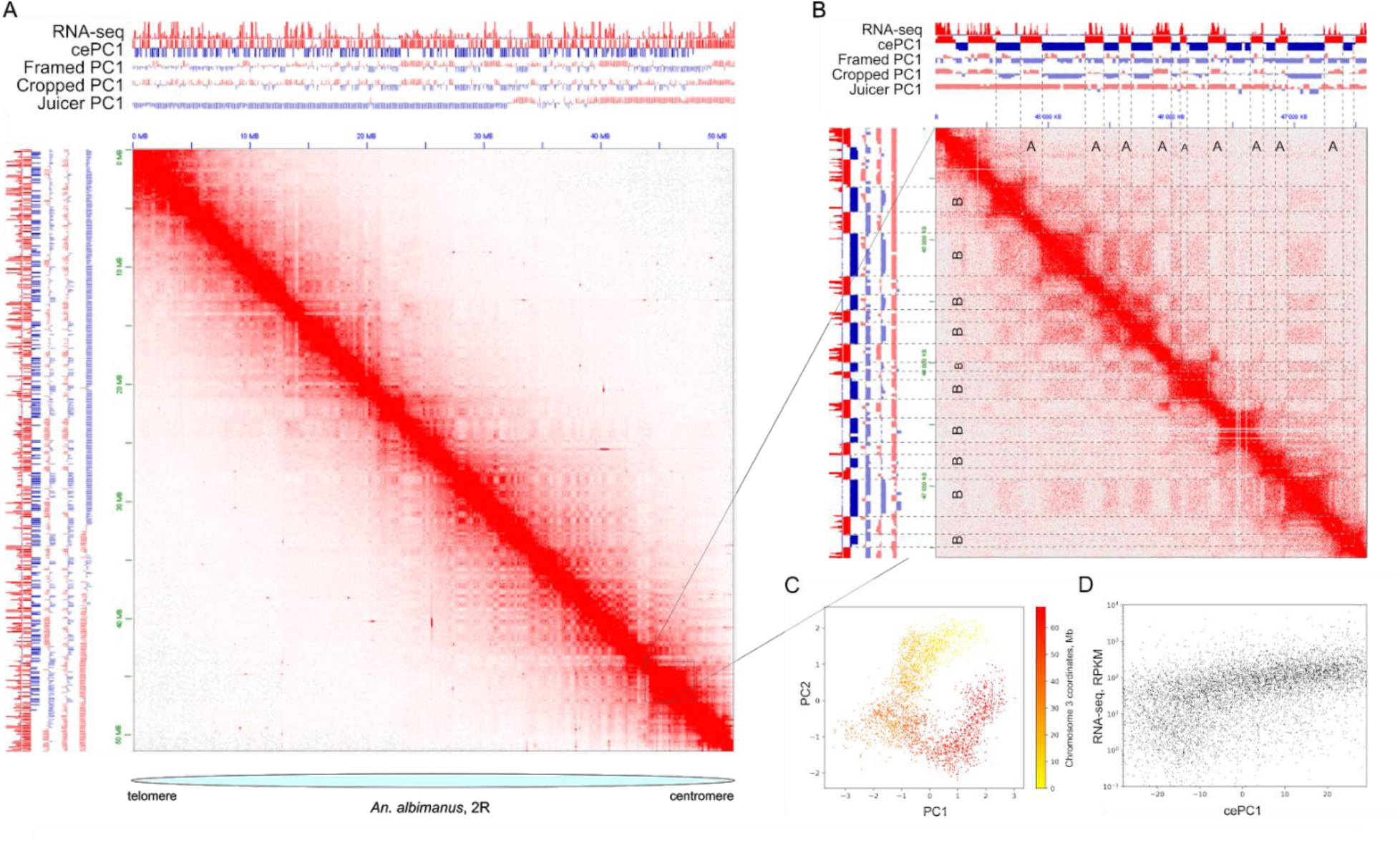
Anopheles genomes are partitioned into compartments. A. Compartmentalization demonstrated for Chr 2R, An. albimanus; B. Region zoomed-in from Fig. 5, A: correspondence between RNA-seq data and PC1 values, obtained using different algorithms; C. Scatter plot of PC1 vs. PC2 values with color representing the coordinates of loci on Chr 3 of An. albimanus illustrates the dependence of PC values on genomic location; D. cePC1 correlation with RNA-seq data.

This discordance between compartmental interactions and PC1-values could be explained by Rabl-like configuration of chromosomes, i.e. clustering of large blocks of centromeric and telomeric heterochromatin and/or elongated shapes of chromosome territories. First, clustering of centromeric and/or telomeric chromatin may dominate the clustering of compartments. This explains those cases where centromeric and telomeric regions display similar PC1 values, whereas the rest of the chromosomal arms display opposite PC1 values (Supplementary Fig. 5, A-I). Second, it was shown in *Drosophila* cells that a Rabl-like configuration results in more elongated shapes of the chromosome territories^37^, preventing clustering of actively expressed genes located far from each other on the centromere-telomere axis.

Supporting these hypotheses, distributions of PC1 values observed in *Anopheles* mosquitoes data were very similar to the distributions obtained from the data from plants^38^ (Fig. 5, C), which also display a Rabl-like configuration of chromosomes. Moreover, for adult *An. merus* samples, where signs of a Rabl-like configuration of chromosomes were much less pronounced (Supplementary Fig. 2), the standard PC1 algorithm was able to define compartments that agree with plaid-patterns on all chromosomes (Supplementary Fig. 5, K-M). Thus, we suggest that the Rabl-like configuration of chromosomes attenuates long-range interactions between compartments.

We developed an approach for robust identification of compartments on chromosomes with Rabl-like configurations. Briefly, our approach relies on local smoothing which reduces noise in long-range interactions followed by the normalization of Pearson’s correlation values within small blocks of the data matrix before PC1 calculation (see Supplementary Note I for details and comparisons of different approaches). We called this new algorithm “contrast enhancement” because the result of Pearson’s correlation scaling within small blocks resembles enhancement of image contrast (Supplementary Fig. 6). We refer to the obtained PC1 values as cePC1 (contrast-enhanced PC1 values). As shown in Fig. 5 B, using contrast enhancement we were able to identify compartments that correspond well to the plaid-pattern observed on the Hi-C maps.

To understand epigenetic mechanisms underlying *Anopheles* chromatin compartments, we performed RNA-seq on the same embryonic stages as used for Hi-C library preparation. We found that compartments show moderate correlation with expression levels (Fig. 5, D), slightly lower correlation with the gene density, and only weakly correlate with GC-content (Supplementary Table 5). Correlations were significantly higher for cePC1, obtained using contrast enhancement, whereas for original PC1 values we found almost no correlation with the aforementioned epigenetic features (Supplementary Table 5). Based on cePC1 values, we split the genome into two non-overlapping A- and B-compartments so that loci belonging to A-compartment show higher gene density, gene expression, and GC-content.

Overall, we have shown that *Anopheles* genomes are partitioned into distinct chromatin compartments, although compartmental interactions are attenuated in embryonic cells, presumably due to the Rabl-like configuration of chromosomes. We developed a computational approach to detect compartments and showed that spatial compartmentalization distinguishes active (gene dense, actively expressed, GC-rich) and inactive (gene-poor, silent, GC-pour) chromatin.

### Transitions between compartments correspond to TAD boundaries in *Anopheles* genomes

In addition to the plaid pattern, Hi-C maps display triangles above the main diagonal (TADs) (Fig. 4; Fig. 6, A-B). For each species, we delineated TADs at 5-kb resolution. The resulting TADs range in size from 15 to 650-kb, with a median size of ~135-kb (Supplementary Table 6). Distribution of TAD sizes was similar in all chromosomes (Fig. 6, C; Supplementary Fig. 7, A); however, within each chromosome, smaller TADs tend to colocalize with A-compartment (euchromatic regions), whereas longer TADs belong to B-compartment (heterochromatin) (Fig. 6, D; Supplementary Fig. 7, B). As evident from the analysis of cePC1-values, TADs longer than 400-kb are almost exclusively located in gene-poor, heterochromatic regions with a low level of gene expression. These TADs do not form long-range interactions, typical for smaller TADs. Comparing positions of large TADs with cytological maps of *Anopheles* polytene chromosomes^39^, we found that these structures often correspond to intercalary heterochromatin blocks (Supplementary Figure 4, Fig. 6, E) and demonstrate a reduction in RNA-seq signal.

**Fig. 6.**
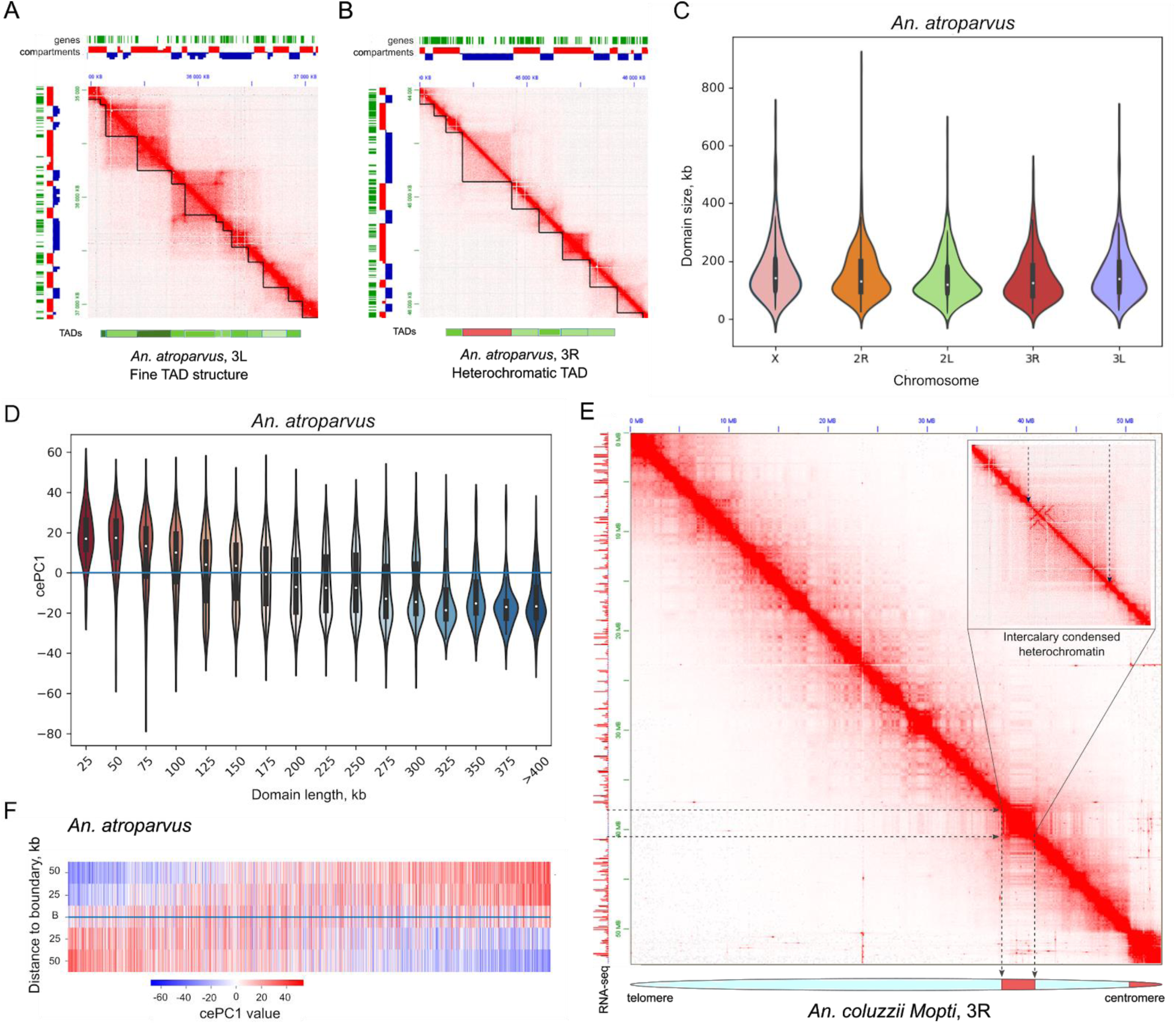
TADs in *Anopheles* genomes. A. Typical TAD structure; B. Example of heterochromatic TAD (marked in red below the contact map region); C. TAD size distributions within the chromosomes; D. Violin-plot illustrating the distributions of cePC1-values for different TAD lengths; E. Intercalary heterochromatic block visualized on the Hi-C map of Chr 3R Anopheles coluzzii Mopti; F. Heatmap of cePC1 values around TAD boundaries. Each line represents one TAD boundary. Panels A-D and F show data for An. atroparvus. Data for other species are shown in Supplementary Fig. 7

We next analyzed TAD boundaries and found that in the majority of cases the boundaries coincide with the transitions between compartments (Fig. 6, F, Supplementary Fig. 7, C), and the magnitude of cePC1 changes was higher around TAD boundaries than within TADs (p-value < 10E-30 for all species, Supplementary Fig. 7, D). We additionally inspected all cases where a strong TAD boundary (top quartile of insulatory scores distribution) separates regions with similar cePC1 values (cePC1 difference below bottom quartile of the distribution), and confirmed that all these cases were due to the errors in TAD or compartment calling.

Thus, we concluded that both TADs and compartments represent similar chromatin features; local insulation from the neighboring genomic regions is captured by TAD-callers, whereas preferences between long-range interactions of these locally insulated genomic regions are captured by cePC1. In-chromosome puffs, representing intercalary heterochromatic regions, correspond to the large TADs depleted for long-range interactions.

### Long-range interactions identified on Hi-C maps and loop validation by FISH

Examination of Hi-C data revealed looping interactions between specific loci (Fig. 4, D, F; Supplementary Fig. 8, A-H). Many of these interactions occurred within the same TAD, and genomic regions involved in looping (hereinafter referred as loop anchors) often formed networks of interactions (Supplementary Fig. 8, A-B). As chromatin loops were previously associated with Polycomb11, we performed H3K27me3 ChIP-seq on *An. atroparvus* and found that indeed anchors of some loops are located within H3K27me3-enriched regions (Supplementary Fig. 8, A-H).

Whereas the vast majority of loops spanned genomic distances of less than one Mb and occurred within TADs, we found notable examples of extremely long-distance loops, connecting loci separated by up to 31-Mb in the linear genome (Fig. 7, A-E). We found 2-6 such giant loops in each species (Supplementary Table 4) and focused on two of them (Table 2): one loop located on chromosome X (X-loop) (Fig. 7, A-E), and another on an autosome (A-loop) (Supplementary Figure 9).

**Table 2.** Coordinates of long-distance loops conserved in different Anopheles species. * Highest percentile score of all distance-normalized interactions between loop anchors computed at 25-kb resolution. Intra PS indicates interactions between anchors located on the same chromosome, inter PS - interactions of anchors of loops on different chromosomes.** Note that the arm 3R of An. atroparvus corresponds to arm 2R of other studied species.

| Species with loop | Chr | 1st anchor | 2nd anchor | PS inter* | Chr | 1st anchor | 2nd anchor | PS inter* | PS intra* |
| --- | --- | --- | --- | --- | --- | --- | --- | --- | --- |
| *An.coluzzii* | X | 7,370,000-7,610,000 | 14,950,000-15,400,000 | 98 | 2R | 41,330,000-41,445,000 | 43,045,000-43,225,000 | 99 | 48 |
| *An.merus* | X | 7,150,000-7,385,000 | 14,610,000-14,900,000 | 99 | 2R | 41,670,000-41,840,000 | 43,635,000-43,945,000 | 99 | 71 |
| *An.stephensi* | X | 9,440,000-9,730,000 | 14,670,000-15,080,000 | 98 | 2R | 37,355,000-37,525,000 | 42,920,000-43,025,000 | 99 | 56 |
| *An.atroparvus* | X | 625,000-780,000 | 10,015,000-10,490,000 | 99 | 3R** | 7,670,000-8,100,000 | 38,625,000-38,770,000 | 98 | 45 |
| *An.albimanus* | X | 4,600,000-4,665,000 | 6,615,000-6,670,000 | 99 | 2R | 34,775,000-34,875,000 | 35,965,000-36,035,000 | 99 | 94 |

**Figure 7.**
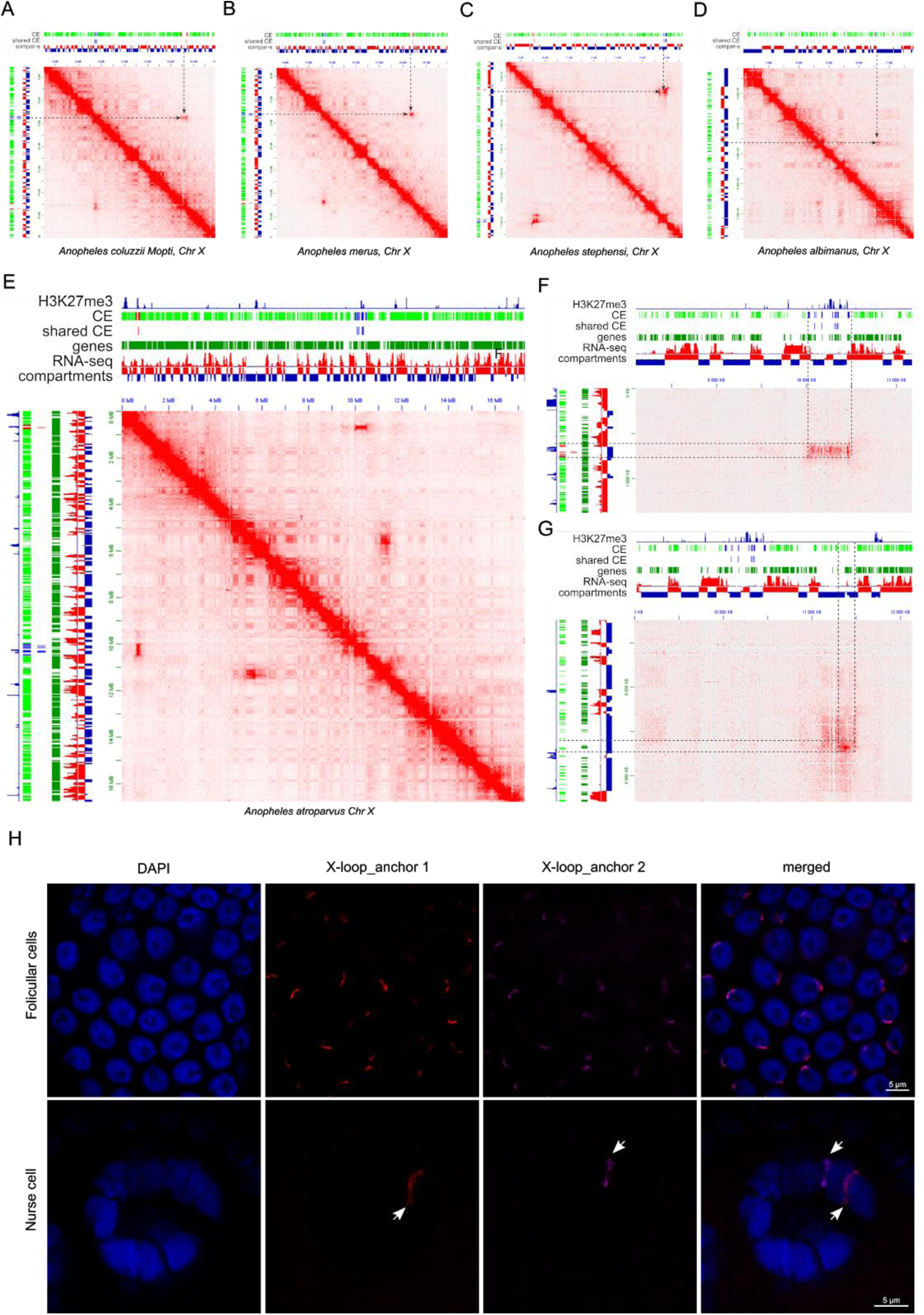
Long-range interactions identified on Hi-C maps and loop validation by FISH. A-D. X-loop in An. coluzzii (A), An. merus (B), An. stephensi (C), An. albimanus (D); E. Region of An. atroparvus chromosome X Hi-C map, showing the X-loop and another giant loop (chrX: 5,320,000-11,510,000) identified on the X chromosome; F and G. Zoomed-in regions of two loops identified on the X chromosome in An. atroparvus H. Colocalization of X-loop anchors identified by 3D FISH in follicular cells and nurse cells of An. atroparvus.

Anchors of these loops were represented by large (~200-300-kb) loci, interacting significantly more than expected at their genomic distance (top 1-2% of all interactions) (Supplementary Fig. 10, A, C, E, G, I; Table 2). Within long anchors of loops, we often observed two pairs of smaller loci (~25-kb) showing interaction frequency peaks (Fig. 7, Supplementary Fig. 9). We noticed that X-loop anchors do not interact with A-loop anchors above the expected level, with only one exception found in *An. albimanus*, where interaction between right anchor of the X-loop and left anchor of the A-loop was among 6% of strongest interchromosomal interactions; however, other pairs of anchors of these loops (left/left, left/right, right/right) did not show strong enrichment (0th, 54th and 64th percentiles), suggesting that X- and A-loop anchors do not colocalize in the nuclei space (Supplementary Fig. 9; Table 2).

To understand whether loops identified in different *Anopheles* species are homologous to each other, we performed whole-genome alignments to identify conserved elements (CEs). We used CEs located within *An. atroparvus* loop anchors to determine anchor synteny between species. The results showed that loops are formed between homologous (syntenic) regions (Fig. 7, Supplementary Fig. 9), except for the left anchor of the X-loop in *An. albimanus*, which does not share a CE with X-loop anchors in other species. Several of the CEs were shared between all species; however, at least in some species these shared CEs were located outside of the anchor regions displaying the highest interaction frequencies (Fig, 7, A; Supplementary Fig. 10), suggesting that none of the identified CEs could explain loop formation.

We next analyzed gene expression and H3K27me3 profiles within loop anchors. Several genes were located within anchors of all examined species (Supplementary Table 7), and some of them were expressed, although not all anchors contain actively expressed genes (Fig. 7, F-G, Supplementary Fig. 9) and the expression level was moderate. In accordance with this observation, loop anchors were enriched in B-compartment regions. Thus, long-range X- and A-loops could not be explained by clustering of active chromatin.

Another type of long-range interaction observed in *Drosophila* genomes is Polycomb-mediated loops^11^. We analyzed the H3K27me3 signal in X- and A-loop anchors and compared it with background genomic signal and with a manually curated set of intra-TAD loops located within H3K27me3-rich loci of *An. atroparvus* (Supplementary Table 8, Supplementary Fig. 8 A-E, Supplementary Fig. 11). This analysis showed only slight H3K27me3 enrichment within X-loop anchors (Fig. 7, F, Supplementary Fig. 11), and no enrichment for A-loop anchors, in contrast to high enrichment of H3K27me3 signal within intra-TAD loop anchors. Thus, X- and A-loops are not formed as a result of Polycomb-mediated interactions.

We next compared Hi-C maps obtained from adult and embryonic *An. merus* tissues, and found that A- and X-loops are present in both datasets (Supplementary Fig. 12, A-B), indicating that these loops are not developmental-stage specific.

Finally, we performed 2D and 3D FISH to validate the interactions between the putative loop anchors in nuclei of follicle cells and ovarian nurse cells of *An. coluzzii*, *An. stephensi*, and *An. atroparvus* (Fig. 7, H). In all tested cases, we found no colocalization of genes at putative anchors in highly polytenized chromosomes of nurse cells by both 2D and 3D FISH. For the *An. coluzzii* 7.5 Mb X-loop, we found 100% colocalization of genes at anchors in follicle cells by 2D FISH and variable colocalization by 3D FISH. For the *An. stephensi* 5.5-Mb A-loop no colocalization in follicle cells was identified by 3D FISH, but the anchors colocalized with high frequency (61% of nuclei, n=157) in ovarian nurse cells with low-level polytene chromosomes.

We tested three loops of different sizes in *An. atroparvus*: 31-Mb A-loop on arm 3R, 6-Mb X-loop on chromosome X, and 12-Mb loop on arm 2R: 11,210,000-11,700,000 – 23,055,000-23,700,000. The anchors are always colocalized in follicle cells in 2D FISH experiments. However, colocalization in 3D FISH experiments was variable: from partial colocalization for the large 31-Mb A-loop on arm 3R (12%, n=109) and medium 12-Mb loop on arm 2R (15%, n=55) to almost constitutive colocalization for the smaller 6-Mb X-loop (97%, n=60). Colocalization for the 6-Mb X chromosome loop was also confirmed by 3D FISH in low-level polytenized chromosomes of nurse cells.

Overall, we validated most of the long-range interactions detected by Hi-C using FISH, although colocalization of loop anchors was found only in a subset of examined cell types. Obtained data showed that *Anopheles* chromatin forms several extremely long-range, locus-specific contacts, which are evolutionarily conserved for ~100 million of years and based on currently unknown molecular mechanisms independent of active transcription or Polycomb-group proteins.

## Spatial contacts of the chromatin in insects and vertebrates are constrained in a genome-size-dependent manner

To study general principles of genome organization in *Anopheles* we analyzed how chromatin contact probability (P) scales with genomic separation s, P(s). As reported in previous studies, contact frequencies decay rapidly as genomic distance increases (Fig. 8, A). Since it was shown previously that P(s) follows a power law, we characterized the exponent by computing slope of the decay curve in log-log coordinates (Fig. 8, B). We found that decay speed is not uniform, and could be described by two different decay phases.

**Fig. 8.**
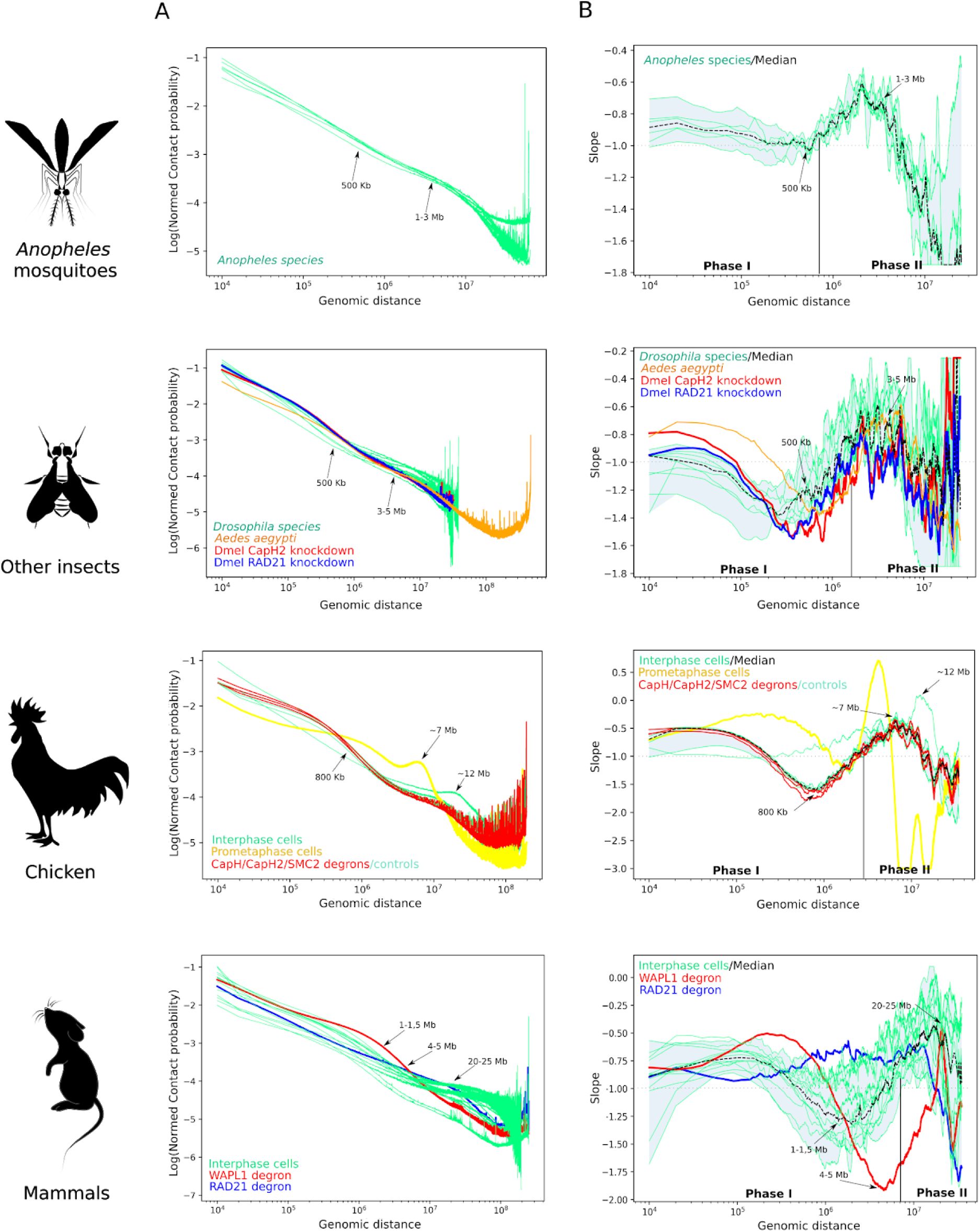
Contact frequency decays non-uniformly with genomic distance. A - dependence of contact probability on genomic distance for Anopheles, other insects, chicken and mammals shown in log-log coordinates. B -slope of the curve depicted in A as a function of genomic distance. Black dashed line shows the median, and the gray area represents minimal and maximal values for several species. Slope-plots for individual species are shown as thin green lines.

The first phase characterized by a U-shaped slope curve occurs between 10-kb and 1-Mb. For distances between 10 kb and ~200-500 kb slope decreases modestly starting from ~-0.8 and reaching the lowest point slightly below −1, which is the characteristic value for the fractal globule, and then starts increasing. At ~500 kb phase II starts, where the slope rapidly increases reaching values around −0.6 for genomic distances ~1-3 Mb and then falls sharply to the values below −1.

These two phases were observed in all *Anopheles* species (Supplementary Fig. 13), and analysis of *Drosophila* and *Aedes* Hi-C datasets demonstrated very similar patterns (Fig. 8, B, Supplementary Fig. 13, A; see Supplementary Table 9 for list of all datasets used to produce Fig. 8). We next analyzed vertebrate Hi-C data, and again found the characteristic shape of the slope curve: the phase I, characterized by prominent decrease of the slope, was much more prolonged than in insects, and reaches a minimum at genomic distances ~800 kb for chicken and ~1-1.5 Mb for mammals; phase II displayed a maximum at ~7 Mb for chicken and ~20-25 Mb for mammals, and after this point we observed a sharp drop of the slope, similar to the insect curves.

We used available Hi-C datasets obtained from cells lacking cohesin subunit RAD21, cohesin release factor WAPBL or cohesin loader NIPBL to confirm that the U-shape of the slope curve observed in phase I reflects the formation of TADs, and that genomic distance characterized by the minimal slope value corresponds to the characteristic TAD length (see Supplementary note II and associated Supplementary figures). This allowed us to provide estimations of TAD lengths without biases introduced by TAD-calling algorithms. The characteristic TAD length in *Anopheles* genomes is around 200-400 kb, which is very close to *Drosophila* species, and slightly smaller than *Aedes* TADs (500-800 kb).

We next aimed to explain changes of the slope during phase II, namely the abrupt drop of contact probabilities observed at genomic distance ~3 Mb for insects, ~7 Mb for chicken and ~20-25 Mb for mammals. We confirmed that this slope decrease was seen in multiple experiments on unsynchronized cells, G2-phase synchronized chicken DT40 cells, G1-phase quiescent chicken erythroblasts and S-phase synchronized *Drosophila* S2 cells or embryonic cells (Supplementary Fig. 13) cells, indicating that there is no cell type or cell cycle specificity and that observed results could not be explained by a fraction of mitotic cells. The only cell type where we did not observe the drop in slope was mammalian post-mitotic neurons (Supplementary Fig. 13, the last row in panel D), suggesting that progression through the M-phase (mitotic phase) of a cell cycle might be essential. Moreover, cohesin- or condensin-mediated loop-extrusion processes are not required to maintain chromosome conformation reflected by the drop of the slope, as we observed the drop under RAD21-degron conditions, NIPBL-degron conditions, both CapH-, CapH2-and SMC2-degron conditions, and in chicken erythrocytes which do not have extrusion-mediated TADs (Fig. 8 and Supplementary Fig. 13, A-D).

We next speculated about possible constraints underlying the observed drop of the slope. We hypothesized that the decrease of contact frequencies observed at large distances might be explained by the specific shape of chromosome territory. Indeed, it is known that chromosome territories are not spherical in mammalian^40,41^ and mosquito^42^ cells. If DNA occupies an ellipsoidal or cylindrical shape, then the radius of the cylinder will determine the characteristic spherical volume inside which the drop of contacts should be in agreement with the fractal globule model. Upon exiting this territory, loci are located in different slices of the cylinder, which makes the likelihood of contacts between them smaller than expected for the fractal globule model.

Following this hypothesis, our data suggest that the interphase chromosomes of all animals fill elongated volumes, characterized by the specific minor radius, different for insects, chicken, and mammals. We estimated that the size of a spherical globule of DNA that fits in this volume is ~1-3 Mb for insects, ~7 Mb for chicken, and ~20-25 Mb for mammals.

## Discussion

Here we used Hi-C data to comprehensively characterize three-dimensional genome organization in five *Anopheles* species. We found that several properties of chromatin organization previously identified in *Drosophila* are shared by *Anopheles* species. In particular, *Anopheles* data suggest that the genome is divided into active and inactive compartments, which correlate with gene expression, and that loci belonging to different compartments are insulated from each other by TAD boundaries.

At the same time, we found some features of genome organization which were not previously described in *Drosophila* or other insect data.

### Evolutionarily conserved long-range chromatin interactions in Anopheles genomes

We discovered that certain genomic regions form giant loops spanning dozens of megabases. Loops described previously in *Drosophila* Hi-C data are typically smaller in size and mainly attributed to interactions between active genes or Polycomb proteins. Our data suggest that loops found in *Anopheles* are formed by other, yet unknown, mechanisms.

FISH analysis showed cell-by-cell variability of contacts between loop anchors, suggesting that 100% formation of the loops is not essential for the function of follicle cells. If such variation exists in embryos then the Hi-C method is sensitive enough to identify the interaction in the assembly of cells. In the case of ovarian nurse cells, we found that colocalization correlates with the low-levels of chromosome polyteny. We conclude that the high polyteny likely creates specific mechanical properties of chromosomes that hinder specific interactions between the loop anchors. An alternative explanation is that highly-polytenized chromosomes do not require such interactions. Interestingly, no specific long-range interactions were found in polytene chromosomes of *D. melanogaster* by microscopic analysis^43^ and Hi-C^11^. However, specific distant chromosomal contacts have been detected in Hi-C heatmaps of *Drosophila* embryos^44^.

### Chromosome territory shape might constrain long-range contacts

We have shown that long-range contacts are constrained in genomes of all examined animal species, and suggested that chromosome territory shape might underline these constraints. It is not known what determines the non-spherical shape of chromosome territories. We confirmed that the drop of the slope is present after CapH2-knockdown in *Drosophila* cells, as well as in chicken cells with CapH, CapH2 or Smc2 proteins degraded, thus condensin proteins probably do not play an essential role in the maintenance cylindrical chromosome territory shapes during interphase. However, it is pertinent to note that chicken cells lacking condensin could not pass through mitosis, and degradation of condensin subunits was induced after cells pass through mitosis with condensin intact, in the next G2-phase. Thus, our analysis shows that condensins are not required to maintain constraints of chromatin contacts reflected by slope-plots, but we can not exclude that they are required to form these constraints during previous mitotic division.

As mentioned above, RAD21 degradation also does not show any effect on the slope changes at large genomic distances, arguing against the role of cohesin in this process. Interestingly, the only cell type where we did not observe a drop of the slope were quiescent mammalian neurons (Supplementary Fig. 13). This allows us to speculate that chromosomes acquire their elongated form during mitosis, and this elongated form might be preserved during the cell cycle. When cells exit the cycle and stay quiescent for a long enough period, diffusion processes intermix of chromatin allowing contacts between more far loci. Similar hypotheses were recently proposed to explain the difference between slope changes in mammalian oocytes^45^, which are arrested at the prophase stage for months and mammalian sperm cells, which are not subjected to such prolonged mitotic arrest^45^.

### Rabl-like chromosome configuration dominates compartmental interactions

Elongated chromosome territory shape reduces the frequency of interactions between loci more than several Mb away from each other. At the same time, we observed the clustering of centromeric and telomeric heterochromatin. According to principal component analysis, these factors dominate genomic compartmentalization, although compartmental interactions are present and well-pronounced at mid-range distances. We proposed a new approach for the identification of compartments in genomes with a Rabl-like configuration and successfully employed it to identify compartments in *Anopheles* genomes.

## Availability of data and materials

All data generated or analyzed during this study are included in this published article and its Supplementary files. The raw sequencing data for five *Anopheles* species have been deposited in the NCBI SRA database with accession numbers PRJNA615788 (RNA-seq raw reads), PRJNA623252 (ChIP-seq raw reads), PRJNA615337 (Hi-C raw reads for embryos), and PRJNA630123 (Hi-C raw reads for adults). Processed data, including genome assemblies, Hi-C heatmaps, RNA-seq and ChIP-seq tracks, TADs and compartments are available at https://genedev.bionet.nsc.ru/Anopheles.html

Source code can be found at github: https://github.com/labdevgen/Anopheles_Ps (analysis of contacts scaling); https://github.com/NuriddinovMA/ABCE (cePC1 computation). Miscellaneous scripts are available upon request.

## Materials and methods

### Mosquito colony maintenance

Laboratory colonies of the following strains were used for the experiments: MOPTI strain for *Anopheles gambiae* (MRA-763); MAF strain for *Anopheles merus* (MRA-1156); STE2 strain for *Anopheles stephensi* (MRA-128); EBRO strain of *Anopheles atroparvus* (MRA-493); STECLA for *Anopheles albimanus* (MRA-126). All mosquito strains were initially obtained through Malaria Research and Reference Reagent Resource Center (MR4) stocks and BEI Resources, NIAID, NIH and maintained in the insectary of the Fralin Life Science Institute at Virginia Polytechnic Institute and State University. Mosquito specimens were hatched from eggs in unsalted water and incubated for 10-15 days undergoing four larvae and pupae developmental stages at 27°C. Adult mosquitoes were maintained in the incubator at 27°C, 75% humidity, with a 12h cycle of light and darkness. 5-7d adult mosquitoes were blood-fed on defibrinated sheep blood using artificial bloodfeeders. Approximately 48-72h post-blood feeding, the egg dishes (Supplementary Fig. 14) were placed and after 15-18h embryos were collected for further experiments.

### In situ Hi-C on mosquito embryos and adults

The step-by-step mosquito Hi-C protocol can be found in Supplementary Protocol I. In brief, procedure for mosquito embryos was modified based on the previously published high-resolution 3C protocol^46^, Drosophila Hi-C protocol^44^, and *in situ* Hi-C protocol for mammalian cells^1^. Mosquito eggs were collected (optimized egg dish can be found in Supplementary Fig. 14) and Hi-C libraries were prepared using nuclei isolated from ~1000-3000 embryos of mixed sexes. Embryos were fixed with 3% formaldehyde at the developmental stage of 15-18h after oviposition. MboI restriction enzyme (NEB, #R0147) with average restriction fragment size ~250-300 bp was used in the experiment. Two biological replicates of Hi-C libraries were generated, prepared with NEBNext® Ultra™ II DNA Library Prep Kit for Illumina (NEB, #E7103), and sequenced using 150-bp pair-ended sequencing on Illumina platform. For Hi-C library preparation from *An. merus* adult mosquitoes, 15 males in two biological replicates were homogenized with liquid nitrogen in a precooled mortar. Separate fixations were done for each Hi-C library of *An. merus* adult males and the downstream procedures were the same as for Hi-C libraries from embryos.

### PacBio sequencing

About 100 one- to two-day old adult males of *An. merus* generated from a single pair of grandparents were used for extraction of the whole genomic DNA. Adult mosquitoes were ground to a fine powder with liquid nitrogen in a precooled mortar. High-molecular weight DNA was extracted using Blood & Cell Culture DNA Midi Kit (Qiagen, Hilden, Germany) with 100/G tips. Extracted DNA was purified with Genomic DNA Clean & Concentrator −10 kit (Zymo Research, Irvine, CA, USA). A total of 17.9 μg of high-molecular weight DNA was used for PacBio sequencing. The DNA concentration of 179 ng/μl in 100 μl was measured by the Qubit Fluorometer (Invitrogen, Carlsbad, CA, USA). PacBio sequencing was performed at the Duke Center for Genomic and Computational Biology (Duke University, Durham, NC, USA). Detailed sequencing reads statistics can be found in Supplementary Table 2.

### ChIP-seq

The anti-trimethyl-histone H3 (Lys27) (Millipore, #07-449) antibody was used for ChIP seq experiment. We followed the protocol previously published in Akulenko et al.^47^ with some modifications. Briefly, ~1000-2000 eggs were bleached, homogenized, and fixed according to optimized ChIP-seq protocol (Supplementary Protocol 2). Cells were lysed and chromatin was sonicated using Bioruptor Diagenode machine, 8-10 cycles of 10/10sec ON/OFF in the lysis buffer (140 mM NaCl, 15 mM HEPES, 1 mM EDTA, 0.5 mM EGTA, 1% triton, 0.5 mM DTT, 0.1% sodium deoxycholate, protease inhibitors) with SDS and N-lauroylsarcosine. After determining the total concentration of chromatin by Qubit, 5-10 μg were used per one ChIP reaction. Additionally, we used *Drosophila melanogaster* chromatin as a spike-in (5-10% of total chromatin amount). Before incubation with antibody, chromatin was pre-cleared by incubation with agarose beads (Pierce™ Protein A/G Agarose, ThermoFisher #20421) for 2h at 4°C with slow rotation. During this time, another aliquot of agarose beads was washed, blocked with BSA, combined with target antibody, and incubated overnight at 4°C with slow rotation. After washing step, pre-cleared chromatin was immunoprecipitated with antibody-agarose beads complexes and incubated overnight at 4°C with slow rotation. Next day, the beads were thoroughly washed in a series of buffers (LB: 150 mM NaCl, 20 mM Tris-HCl, 2 mM EDTA, 1% triton, 0.1% SDS; HB: 500 mM NaCl, 20 mM Tris-HCl, 2 mM EDTA, 1% triton, 0.1% SDS; LiB: 0.25 M LiCl, 10 mM Tris-HCl, 1 mM EDTA, 1% NP-40; TE buffer). DNA was eluted with 250 μL elution buffer (EB: 1% SDS, 0.1 mM NaHCO_3_) by incubation at 65°C for 10 minutes. To revert cross-linking, the DNA was incubated at 65°C overnight in presence of 0.25M NaCl, 10 mM EDTA and 40 mM TrisHCl. DNA was extracted using phenol-chloroform mix followed by ethanol precipitation. ChIP-seq libraries were prepared for sequencing using NEBNext® Ultra™ II DNA Library Prep Kit for Illumina (NEB, #E7103). The detailed ChIP-seq protocol can be found in Supplementary materials.

### RNA-seq

1500-2000 embryos of 15-18h *Anopheles* mosquitoes were bleached for 5 minutes. Total RNA was extracted following the Monarch Total RNA Miniprep Kit (NEB #T2010S) protocol with minor modifications. Mosquito embryos were homogenized in 800 μL of lysis buffer with 2 mL Dounce homogenizer. Samples were incubated at RT for 10 minutes, then proteinase K was added and samples were incubated for an additional 5 minutes at 55°C. After that, the tubes were centrifuged at max speed for 2 minutes. Supernatant was transferred to fresh RNAse/DNAse-free tubes and proceeded with gDNA removal columns. The incubation time with DNAse was increased to 20 minutes in total. Total RNA was eluted with 50 μL H2O. Sample concentration was verified with Nanodrop and 1 μg of total RNA was used for the next procedures. Samples were prepared for Illumina sequencing with NEBNext® Ultra™ II RNA Library Prep Kit for Illumina (NEB #E7775) accompanied by NEBNext Poly(A) mRNA Magnetic Isolation Module (NEB #E7490) with RNA insert size of 200 bp.

### Ovary preservation and polytene chromosome preparation

To prepare high-quality polytene chromosome slides we followed the protocol described in Sharakhova et al.^32^ with minor exceptions. Approximately 24-30h after the second or third blood feeding (the timeline for Christopher’s III developmental stage varies for different species and should be estimated by visual inspection), ovaries were fixed in Carnoy’s solution (3:1, ethanol: glacial acetic acid by volume), kept at RT for 24h, then stored at −20°C for a long term.

At least one week after fixation, ovaries were dissected in Carnoy’ solution. Cleaned and separated follicles were then placed on a slide in a drop of 50% propionic acid for ~5 min (5-10 follicles per slide), where they were macerated and squashed in a fresh portion of 50% propionic acid. The quality and banding pattern were briefly examined using a phase-contrast microscope (1,000×) and high-quality preparations were proceeded with snap-freezing in liquid nitrogen. After freezing the slides were immediately placed in pre-chilled 50% ethanol and kept at −20°C overnight. Next day, after removing coverslips, preparations were dehydrated in ethanol series (50, 70, and 96%), air-dried, and the quality of polytene chromosomes was checked. High-quality slides were placed in a cardboard holder and stored at RT up to 3 months.

### 2D-FISH

Probes were prepared by Random Primer Labeling method described in Protocols for Cytogenetic Mapping of Arthropod Genomes^48^. gDNA was freshly extracted using Monach Genomic DNA purification kit, PCR product amplification was performed by regular PCR with DreamTaq/Q5 polymerase. PCR Product was purified with Qagen purification columns and ~200-250 ng were used for 25 μL labeling reaction. After overnight incubation in the thermocycler at 37C°, labeled probes were precipitated with 96% ethanol, dried, and dissolved in 30 μL hybridization buffer. 10-15μL of one probe were applied to slide. Prepared slides were incubated for 25 minutes at 70°C in a humid chamber following the overnight incubation at 39°C. Washing steps included 2 times wash in 1xSSC at 39°C for 20 minutes, 1 time wash in 1xSSC at RT for 20 minutes, 1 time wash in 1xPBS at RT for 10 minutes. One drop of ProLong™ Gold Antifade Mountant with DAPI (ThermoFisher #P36931) was added to the slide for protection. Fluorescent signals were detected and recorded with a Zeiss LSM 710 laser scanning microscope (Carl Zeiss Microimaging GmbH, Oberkochen, Germany). Set of primers for PCR product amplification can be found in Supplementary Table 10. All PCR primers were designed against unique exon regions.

### 3D-FISH

Fluorescent probes were prepared by the same method as for 2D-FISH. Probe pellets were dried and resuspended in 20-30 hybridization buffer. 20μL of one probe were used per one experiment where the total volume of the hybridization probe solution contained at least 80 μL. Probes were denatured in Thermomixer at 90°C for 5 minutes, then transferred to 39°C and pre-annealed for at least 30 minutes.

24-30h after blood feeding ovaries were dissected in 1xPBS and placed in 1mL 1xPBST. Fixation was performed in 4% paraformaldehyde solution for 20 minutes with rotation. Then, ovaries were washed in PBS and treated with 0.2 μg/μL RNAse solution in PBS at 37°C for 20 minutes. After rinsing in PBS, ovaries were placed in 1% Triton-X-100/0.1M HCl solution and incubated at RT for 20 minutes with rotation. After brief washing, DNA denaturation was performed in 50% formamide/2xSSC solution at 75°C for exactly 30 minutes. Then, ovaries were rinsed in 100μL of hybridization buffer which was replaced with hybridization mix. Tubes were incubated at 39°C overnight with slow mixing. Next day, the ovaries were washed in a series of washing solutions at 39°C with rotation: 3 washes in 50% formamide/2xSSC; 3 washes in 2xSSC. Drop of ProLong™ Gold Antifade Mountant with DAPI was added and ovaries were placed on a 3D-FISH slide. Fluorescent signals were detected and recorded using a Zeiss LSM 880 confocal laser scanning microscope (Carl Zeiss AG, Oberkochen, Germany).

## Computational methods

### Hi-C data processing

Raw reads were processed using Juicer protocol^49^. Contacts were normalized using KR-normalization. Expected contact counts were obtained by dumping expected vectors using juicer tools *dump* tool.

### Genome assembly

3D-DNA pipeline4 was employed to assemble the genomes de novo using the generated Hi-C data set. Misassemblies were identified and fixed manually using assembling mode in Juicebox software^49,50^. The physical genome maps were used to assess the assemblies^23–27^.

### Whole-genome alignments and CE calling

For pairwise whole-genome alignments we used LastZ tool^51^ with parameters high-scoring segment pairs (HSPs) threshold (--hspthresh) = 3000, interpolation threshold (-- inner) = 2000, step size (--step) = 20, alignment processed with gap-free extension of seeds, gaps extension of HSPs and excluded chaining if HSPs (--gfextend --nochain -- gapped).

To find alignment blocks we used Mugsy^52^ with default parameters. To call CE, we used PhastCons. Parameter tunings were guided by a software recommendation, but 65% exon coverage by CE was an unreachable goal. To solve this problem, we analysed a relation between a summarized length of CEs and the parameters, when the phylogenetic information threshold was near 10 bits. We observed a “plateau” in increasing summarized CEs length when target coverage and expected length of CE varied between 0.40 to 0.60 and from 30 to 40 nucleotides, respectively. Thus, for each alignment we fixed parameters at the point when summarized CEs length reaches plateau.

### Phylogenomic analysis and calculation of divergence times

We analysed genome assemblies of the six mosquito species available from VectorBase^53^ release VB-2019-08 with the Diptera dataset of the Benchmarking Universal Single-Copy Orthologue (BUSCO v3.0.2) assessment tool^54^. From the results, we identified 1’258 BUSCO genes present as single-copy orthologues in all six species. We aligned the protein sequences for each BUSCO with MAFFT v7.450^55^ and then filtered/trimmed them with TrimAl v1.2^56^ using automated parameters to produce a concatenated superalignment. AliStat v1.12^57^ assessment of the superalignment: 6 sequences; 854’431 sites; completeness score: 0.96383. We then estimated the phylogeny using RAxML v8.0.0^58^ with the PROTGAMMAJTT model with 100 bootstrap samples. Rooted with *Aedes aegypti*, we converted the molecular phylogeny to an ultrametric time-calibrated phylogeny using the chronos function in R^59^ with the discrete model and fixing the *An. gambiae* complex age at 0.5 million years according to Thawornwattana et al.^19^ and the *Anopheles* genus age at 100 million years in line with Neafsey et al.^18^ and the geological split of western Gondwana.

### ChIP-seq data processing

All ChIP-seq data were processed using the Aquas pipeline (https://github.com/kundajelab/chipseq_pipeline) stopped at the signal stage.

### RNA-seq data processing

All data were processed using standard protocols with HISAT2, bedtools Genome Coverage, StringTie tools^60^. The sequencing data were uploaded to the Galaxy web platform, and we used the public server at usegalaxy.org to analyze the data^60^. We used averaged TPM values obtained from three biological replicas for all downstream analysis.

### Compartment calling

The default approach for PC1 value computation relied on using juicer tools *eigenvector* tool^49^ at 25-, 50 or 100-kb resolutions. For other approaches, explained in details in Supplementary Note I, we used custom R-scripts. To compute PC1 values for intrachromosomal submatrices we used window (frame) size equal to 10-Mb.

### TAD calling

To call TADs, we first used Armatus, Lavarbust, Dixon callers and hicExplorer *findTADs* utilities^61–64^ with default parameters at 25- and 5-kb resolutions. Visual inspection of obtained TADs revealed that results were similar for the Armatus, Lavarbust and hicExplorer algorithms, whereas the Dixon caller resulted in large, Mb-scaled TADs, which did not correspond well with triangles visible on heatmaps. Although this difference is most probably due to default parameters of the Dixon TAD caller, originally developed on mouse and human data, and results could be improved by tweaking parameters, we decided to focus on hicExplorer-based TADs at 5-kb resolution because visual assessment suggested that this caller provided the best results. By tweaking parameters as suggested in the manual, we found that a delta value of 0.05 provided TADs most closely matching triangles on heatmaps; changing other parameters did not lead to substantial improvements of TADs. Finally, resulting TADs were visually inspected to correct boundary positions in problematic regions (long heterochromatin blocks, gaps etc.).

### Slope plot analysis

To produce slope plots, we computed the best linear fit of the log-log scaled dependence of expected contact frequencies from genomic distance. For each distance, we used N points to obtain the local fit. As contacts become noisier with distance, we increased N at larger distances. In particular, to compute slope at distance X we used all expected values in the interval [X, 50 000 + 1.25*X]. We computed slope for each chromosome with a length of more than 20 Mb individually, and then used the median of all obtained values. We cropped the resulting plot at 50 Mb because slope values become too noisy beyond this point.

## Supporting information

Supplementary Materials_all

## Funding

This work was supported by the NSF grant MCB-1715207, NIH NIAID grant R21AI135298, and the USDA National Institute of Food and Agriculture Hatch project 223822 to IVS. The reported study of *An. atroparvus* was partly funded by RFBR according to the research project №19-34-50051 to IVS and VL. PacBio sequencing of *An. merus* was funded by a grant from the University of Lausanne Department of Ecology and Evolution to RMW and NIH grants AI133571 and AI121284 to ZT. MJMFR, LR, and RMW were supported by Novartis Foundation for medical-biological research grant #18B116 and Swiss National Science Foundation grant PP00P3_170664. VL was partly supported by the Fulbright Foreign Student Program, Grantee ID: 15161026. Hi-C data analysis was supported by the Ministry of Education and Science of Russian Federation, grant #2019-0546 (FSUS-2020-0040).

## Acknowledgments

The following reagents were obtained through BEI Resources, NIAID, NIH: *An. coluzzii*, Strain MOPTI, Eggs, MRA-763, contributed by Gregory C. Lanzaro; *An. merus*, Strain MAF, MRA-1156, contributed by Maureen Coetzee; *An. atroparvus*, Strain EBRO, Eggs, MRA-493, contributed by Carlos Aranda and Mark Q. Benedict; *An. albimanus*, Strain STECLA, Eggs, MRA-126, contributed by Mark Q. Benedict. All computations were performed using nodes of the high-throughput cluster of the Novosibirsk State University, and bioinformatics resource center of the Institute of Cytology and Genetics (Budget Project 0324-2019-0041-C-01).

## Supplementary materials

all Supplementary Notes, Figures, Tables, and Protocols referred to in this manuscript have been compiled into Supplementary Materials_all file.

## Notes

### Competing Interest Statement

The authors have declared no competing interest.

