## Supplementary Materials_all for "*Anopheles* mosquitoes revealed new principles of 3D genome organization in insects"

### Supplementary materials
### Supplementary notes

#### **Supplementary Note I.** *Detection of compartmental interactions on chromosomes in Rabl-like configuration*

We argued that standard algorithms designed for identification of compartments failed to detect compartments on most *Anopheles* chromosomes because long-range interactions between centromeric and/or telomeric regions dominate mid-range interactions between compartments.

To overcome this issue, we first attempted to crop centromeric and telomeric heterochromatin and calculate PC1 values for the remaining regions using a standard approach. However, PC1 values obtained on cropped data failed to describe plaid-pattern (see *cropped* tracks on Fig. 4 and associated supplementary figures).

Next, we decided to use large (1-15 Mb) regions (frames) when computing Pearson's correlation on cropped data. The size of these regions was motivated by the analysis of the speed of contacts decay, i.e. P(s) curves, which show that speed of contact frequency decay increases rapidly above this distance (see results). We speculate that PC1 obtained within each of these regions would reflect local compartments structure rather than capturing chromosome-wide patterns. Such local PC1-scores were recently proposed to define compartments on meiotic chromosomes in mammals, where long-range compartmental interactions were also attenuated due to specific configuration of the chromosomes[1]. Splitting chromosomes onto consecutive Mb-length regions indeed allows capturing kilobase-lengths compartments consistent with plaid-pattern contacts for several regions (see *framed* tracks in Fig. 4 and associated supplementary figures). However, in many cases, especially when a region contains a strongly insulated boundary between two blocks of chromatin (i.e. intercalary heterochromatin), the fine structure of compartments was not resolved by framed PC1 values because strong local insulation dominates the variation of relatively few mid-range interactions within the rest of the regions.

To overcome this issue we decided to use all intrachromosomal contacts but scale correlation values so that they are not dominated by effects of Rabl-like configuration or heterochromatin boundaries. This procedure, which we called contrast enhancement, implies 1.) cropping centromeric/telomeric heterochromatin; 2.) averaging (smoothing) of observed/expected values for the whole matrix except cropped regions; 3.) mean subtraction and calculation of Pearson's correlation for cropped matrix, as in the standard juicer[2] algorithm; 4.) normalization of all correlation coefficients to ensure they are in the diapason -1..+1 (contrast enhancement) 5.) splitting the remaining portion of the genome into Mb-sized frames; 6.) calculation of PC1 values within each frame, resulting in contrast-enhanced PC1 (cePC1) values; 7.) assigning PC1 signs within each frame according to the correlation with RNA-seq data. We illustrated how we transform data during these steps in Supplementary Fig. 6.

Contrast enhancement allowed us to call compartments on all chromosomes of all *Anopheles* species, excluding the chromosome 2R of *An. stephensi*, which harbors large polymorphic inversion 2Rb. For this chromosome, we were able to call compartments by splitting the chromosome into three regions (telomere-proximal region up to inversion breakpoint; inverted region itself; and centromere-proximal region downstream of inversion breakpoint) and applying the algorithm to each of these regions independently.

**Supplementary note II.** *The U-shape of the slope-plot reflects the formation of TADs.*

We speculate that the U-shape of the slope (phase I) reflects the formation of TADs. Loop extrusion and/or chromatin aggregation which occurs within TADs decreases speed of decay of intra-TAD contacts, making it lower than speed of decay of inter-TAD contacts. The fraction of intra-TAD contacts decreases with distance, thus average contact frequency decays more rapidly than expected for inter-TAD contacts to catch up the difference between inter- and intra-TAD contacts. This trend reverses at distances corresponding to a character intra-TAD length, which is reflected by the lowest point of the phase I.

To test this hypothesis we analyzed Hi-C data on HCT116 cells which lack loop extrusion process due to active degradation of the key cohesin component, RAD21[3]. These data show only a weak decrease of slope in the phase I, in strong contrast with other mammalian cell types and more similar to the phase I observed in the *Anopheles* species. The same slope curve shape was obtained by re-analyzing our data on chicken erythroblasts, which are unique somatic cells lacking TADs[4]. Meanwhile, other chicken cell types display phase I similarly to mammals.

The lowest point of the phase I in chicken and mammalian cell types fits with character TAD length in these species (800-kb and 1-Mb, respectively). Moreover, in HAP1 cells lacking WAPBL, the condensin release factor, the slope minimum was shifted to longer genomic distances (4-5-Mb) in accord to the longer cohesin loops reported under WAPL-knockdown condition[5]. Finally, the U-shape was observed under Rad21-knockdown condition in *Drosophila*, which fits with the previous observation that TAD-forming mechanisms are independent of RAD21-mediated extrusion in *Drosophila*[6]. These facts support our conclusion that the U-shape of the slope during phase I reflects TADs, and the lowest point corresponds to the character TAD size. This view is additionally supported by recent polymer models described in Fudenberg et al.[7]

#### Supplementary tables

**Supplementary Table 1.** Hi-C libraries statistics (2 replicas combined):

| **Parameters** | ***An. albimanus*** AalbS2_V4 | ***An. atroparvus*** AatrE3_V4 | ***An. coluzzii Mopti*** AcolNg_V4 | ***An. merus_ embryos*** AmerR4_V4 | ***An. merus_ adults*** AmerR4_V4 | ***An. stephensi*** AsteI2_V4 |
|---|---|---|---|---|---|---|
| **Sequenced Read Pairs** | 67,591,619 | 217,895,615 | 231,961,292 | 188,700,705 | 129,643,770 | 253,760,093 |
| **Normal Paired** | 30,787,793 (45.55%) | 103,420,169 (47.46%) | 108,826,067 (46.92%) | 78,420,401 (41.56%) | 53,633,589 (41.37%) | 129,373,218 (50.98%) |
| **Chimeric Paired** | 30,295,709 (44.82%) | 72,384,597 (33.22%) | 81,994,702 (35.35%) | 53,989,004 (28.61%) | 35,096,848 (27.07%) | 64,754,256 (25.52%) |
| **Chimeric Ambiguous** | 2,614,742 (3.87%) | 15,313,087 (7.03%) | 33,974,675 (14.65%) | 16,609,540 (8.80%) | 11,671,013 (9.00%) | 21,361,880 (8.42%) |
| **Unmapped** | 3,893,375 (5.76%) | 26,777,762 (12.29%) | 7,165,848 (3.09%) | 39,681,760 (21.03%) | 29,242,320 (22.56%) | 38,270,739 (15.08%) |
| **Ligation Motif Present** | 46,422,310 (68.68%) | 142,902,260 (65.58%) | 168,629,612 (72.70%) | 125,132,781 (66.31%) | 82,596,867 (63.71%) | 75,068,793 (29.58%) |
| **Alignable (Normal+Chimeric Paired)** | 61,083,502 (90.37%) | 175,804,766 (80.68%) | 190,820,769 (82.26%) | 132,409,405 (70.17%) | 88,730,437 (68.44%) | 194,127,474 (76.50%) |
| **Unique Reads** | 56,129,336 (83.04%) | 150,572,935 (69.10%) | 165,637,626 (71.41%) | 114,668,816 (60.77%) | 77,371,708 (59.68%) | 135,257,540 (53.30%) |
| **PCR Duplicates** | 3,502,506 (5.18%) | 22,928,735 (10.52%) | 21,846,321 (9.42%) | 15,999,379 (8.48%) | 11,326,157 (8.74%) | 55,601,774 (21.91%) |

| | | | | | | |
|---|---|---|---|---|---|---|
| **Optical Duplicates** | 1,451,660 (2.15%) | 2,303,096 (1.06%) | 3,336,822 (1.44%) | 1,741,210 (0.92%) | 32,572 (0.03%) | 3,268,160 (1.29%) |
| **Library Complexity Estimate** | 487,552,124 | 597,237,052 | 740,698,381 | 489,080,571 | 317,065,109 | 260,265,053 |
| **Intra-fragment Reads** | 2,461,570 (3.64% / 4.39%) | 8,205,249 (3.77% / 5.45%) | 11,381,671 (4.91% / 6.87%) | 5,708,656 (3.03% / 4.98%) | 3,065,395 (2.36% / 3.96%) | 16,939,144 (6.68% / 12.52%) |
| **Below MAPQ Threshold** | 682,547 (1.01% / 1.22%) | 5,901,863 (2.71% / 3.92%) | 29,040,047 (12.52% / 17.53%) | 22,026,541 (11.67% / 19.21%) | 13,482,444 (10.40% / 17.43%) | 5,777,870 (2.28% / 4.27%) |
| **Hi-C Contacts** | 52,985,219 (78.39% / 94.40%) | 136,465,823 (62.63% / 90.63%) | 125,215,908 (53.98% / 75.60%) | 86,933,619 (46.07% / 75.81%) | 60,823,869 (46.92% / 78.61%) | 112,540,526 (44.35% / 83.20%) |
| **Ligation Motif Present** | 23,987,831 (35.49% / 42.74%) | 60,551,709 ( 27.79% / 40.21%) | 61,809,999 ( 26.65% / 37.32%) | 40,425,437 (21.42% / 35.25%) | 26,855,598 (20.71% / 34.71%) | 27,906,559 (11.00% / 20.63%) |
| **3' Bias (Long Range)** | 50% - 50% | 50% - 50% | 50% - 50% | 50% - 50% | 50% - 50% | 50% - 50% |
| **Pair Type %(L-I-O-R)** | 25% - 25% - 25% - 25% | 25% - 25% - 25% - 25% | 25% - 25% - 25% - 25% | 25% - 25% - 25% - 25% | 25% - 25% - 25% - 25% | 25% - 25% - 25% - 25% |
| **Inter-chromosomal** | 6,862,999 ( 10.15% / 12.23%) | 24,255,276 ( 11.13% / 16.11%) | 21,566,658 ( 9.30% / 13.02%) | 29,961,382 (15.88% / 26.13%) | 6,270,919 ( 4.84% / 8.10%) | 18,456,987 (7.27% / 13.65%) |
| **Intra-chromosomal** | 46,122,220 (68.24% / 82.17%) | 112,210,547 (51.50% / 74.52%) | 103,649,250 (44.68% / 62.58%) | 56,972,237 (30.19% / 49.68%) | 54,552,950 (42.08% / 70.51%) | 94,083,539 (37.08% / 69.56%) |

| | | | | | | |
|---|---|---|---|---|---|---|
| **Short Range (<20Kb)** | 13,468,625 (19.93% / 24.00%) | 25,525,319 ( 11.71% / 16.95%) | 29,262,843 ( 12.62% / 17.67%) | 13,002,707 (6.89% / 11.34%) | 16,850,740 (13.00% / 21.78%) | 37,468,994 (14.77% / 27.70%) |
| **Long Range (>20Kb)** | 32,653,284 (48.31% / 58.18%) | 86,684,768 ( 39.78% / 57.57%) | 74,385,329 ( 32.07% / 44.91%) | 43,969,011 (23.30% / 38.34%) | 37,702,135 (29.08% / 48.73%) | 56,613,127 (22.31% / 41.86%) |

**Supplementary Table 2.** PacBio sequencing reads statistics.

| | Root folder | Sub folder | Mean read length | Median read length | Number of reads | Read length N50 | Total bases |
|---|---|---|---|---|---|---|---|
| **1** | r54089_20180220_133714 | 1_A01 | 5 124,70 | 3 072 | 931 662 | 9 931 | 4 774 508 323 |
| **2** | r54089_20180220_133714 | 2_B01 | 5 290,30 | 3 186 | 901 281 | 10 262 | 4 768 090 239 |
| **3** | r54089_20180731_171253 | 1_A01 | 7 402,30 | 4 911 | 964 615 | 13 282 | 7 140 360 112 |
| **4** | r54089_20180801_111617 | 1_A01 | 5 591,00 | 3 412 | 1 110 790 | 10 534 | 6 210 374 362 |
| **5** | r54089_20180801_111617 | 2_B01 | 5 270,70 | 3 134 | 963 799 | 10 036 | 5 079 882 453 |
| **6** | r54089_20180802_142428 | 1_A01 | 6 051,50 | 4 039 | 1 009 269 | 10 924 | 6 107 554 776 |
| **7** | r54089_20180802_142428 | 2_B01 | 5 824,80 | 3 871 | 980 478 | 10 504 | 5 711 084 909 |
| **8** | r54089_20180802_142428 | 3_C01 | 7 934,30 | 5 459 | 674 172 | 14 364 | 5 349 079 444 |
| **9** | r54089_20180802_142428 | 4_D01 | 6 337,90 | 4 251 | 1 027 733 | 11 495 | 6 513 622 660 |
| **10** | r54089_20180802_142428 | 5_E01 | 5 800,20 | 3 818 | 1 009 509 | 10 581 | 5 855 362 794 |
| **11** | r54089_20180802_142428 | 6_F01 | 5 850,70 | 3 664 | 968 902 | 11 061 | 5 668 761 247 |

**Supplementary Table 3.** Polymorphic inversions identified using Hi-C data.

| Species | Chromosome | Genomic coordinates | Size |
|---|---|---|---|
| ***An. stephensi*** | 2R | 21,140,000 - 37,180,000 | 16 Mb |
| ***An. atroparvus*** | 2L | 2,980,000 - 13,940,000 | 11 Mb |
| ***An. coluzzii* Mopti** | 2R | 30,940,000 - 35,340,000 | 4.4 Mb |
| ***An. merus*** | 2R | 60,175,000-62,930,000 | 2.8 Mb |

**Supplementary Table 4.** All loops detected using Hi-C heat map.

| Species | Chr | 1st anchor | 2nd anchor | Total amount of loops per genome |
|---|---|---|---|---|
| ***An. coluzzii*** | X: | 3,410,000-3,535,000 | 6,020,000-6,120,000 | 4 |
| | X: | 7,370,000-7,610,000 | 14,950,000-15,400,000 | |
| | 2R: | 41,330,000-41,445,000 | 43,045,000-43,225,000 | |
| | 2R: | 41,125,000-41,300,000 | 46,435,000-46,785,000 | |
| ***An. merus**** | X: | 7,150,000-7,385,000 | 14,610,000-14,900,000 | 9 |
| | X: | 3,335,000-3,395,000 | 5,740,000-5,925,000 | |
| | 2R (A): | 26,783,000-26,815,000 | 39,725,000-39,825,000 | |
| | 2R: | 41,670,000-41,840,000 | 43,635,000-43,945,000 | |
| | 3R (A): | 19,780,000-19,935,000 | 35,890,000-35,990,000 | |
| | 3R (A): | 19,780,000-19,935,000 | 37,515,000-37,740,000 | |
| | 3R (A): | 29,975,000-30,090,000 | 44,340,000-44,470,000 | |
| | 3R (A): | 13,840,000-14,040,000 | 46,090,000-46,170,000 | |
| | 3R (A): | 17,220,000-17,330,000 | 45,910,000-45,940,000 | |
| ***An. stephensi*** | X: | 7,215,000-7,355,000 | 9,670,000-9,725,000 | 6 |
| | X: | 9,435,000-9,815,000 | 14,665,000-15,130,000 | |
| | 2R: | 37,345,000-37,525,000 | 42,900,000-43,125,000 | |
| | 3R: | 10,780,000-10,910,000 | 38,430,000-38,580,000 | |

| | | | | |
|---|---|---|---|---|
| | 3R: | 16,075,000-16,295,000 | 24,265,000-24,585,000 | |
| | 3R: | 16,075,000-16,295,000 | 27,435,000-27,610,000 | |
| ***An. atroparvus*** | X: | 5,320,000-5,755,000 | 11,100,000-11,510,000 | 5 |
| | X: | 620,000-790,000 | 10,010,000-10,505,000 | |
| | 2R: | 11,210,000-11,700,000 | 23,055,000-23,700,000 | |
| | 3R: | 7,620,000-8,160,000 | 38,550,000-38,950,000 | |
| ***An. albimanus*** | X: | 4,595,000-4,715,000 | 6,590,000-6,675,000 | 3 |
| | 2R: | 34,730,000-34,905,000 | 35,925,000-36,070,000 | |
| | 2L: | 25,600,000-25,685,000 | 29,255,000-29,380,000 | |

* For *An. merus* some loops were identified only in adult mosquito’s data. These loops and marked with letter *(A)*

**Supplementary Table 5.** Correlation of compartments with RNAse, gene density and RNA-seq data.

| species | chrm | RNAseq | | | | %GC | | | | Number of genes | | | |
|---|---|---|---|---|---|---|---|---|---|---|---|---|---|
| | | juicer | cropped | framed | ce | juicer | cropped | framed | ce | juicer | cropped | framed | ce |
| *An. albimanus* | X | 0,60 | 0,71 | 0,69 | 0,67 | 0,23 | 0,21 | 0,19 | 0,18 | 0,52 | 0,59 | 0,58 | 0,59 |
| | 2R | 0,01 | 0,11 | 0,23 | 0,64 | 0,22 | 0,23 | 0,05 | 0,04 | 0,01 | 0,10 | 0,15 | 0,50 |
| | 2L | 0,61 | 0,58 | 0,34 | 0,50 | 0,04 | 0,01 | 0,02 | 0,02 | 0,45 | 0,40 | 0,24 | 0,37 |
| | 3R | 0,24 | 0,28 | 0,32 | 0,53 | 0,05 | 0,02 | 0,02 | 0,02 | 0,22 | 0,18 | 0,22 | 0,34 |
| | 3L | 0,65 | 0,34 | 0,41 | 0,62 | 0,13 | 0,54 | 0,28 | 0,06 | 0,55 | 0,34 | 0,26 | 0,44 |
| *An. atroparvus* | X | 0,01 | 0,54 | 0,65 | 0,68 | 0,23 | 0,01 | 0,11 | 0,14 | 0,01 | 0,32 | 0,41 | 0,47 |
| | 2R | 0,13 | 0,34 | 0,23 | 0,49 | 0,34 | 0,18 | 0,10 | 0,10 | 0,08 | 0,16 | 0,09 | 0,33 |
| | 2L | 0,61 | 0,34 | 0,37 | 0,60 | 0,30 | 0,05 | 0,11 | 0,19 | 0,49 | 0,25 | 0,29 | 0,39 |
| | 3R | 0,02 | 0,12 | 0,30 | 0,54 | 0,21 | 0,12 | 0,27 | 0,20 | 0,12 | 0,04 | 0,25 | 0,42 |
| | 3L | 0,13 | 0,48 | 0,43 | 0,48 | 0,13 | 0,23 | 0,20 | 0,22 | 0,14 | 0,34 | 0,31 | 0,37 |
| *An. coluzzii* | X | 0,36 | 0,61 | 0,58 | 0,60 | *0,06* | *0,04* | *0,03* | *0,01* | *0,29* | *0,35* | *0,34* | *0,34* |
| | 2R | 0,06 | 0,04 | 0,25 | 0,59 | *0,28* | *0,23* | *0,02* | *0,05* | *0,05* | *0,04* | *0,13* | *0,35* |
| | 2L | 0,04 | 0,21 | 0,35 | 0,56 | *0,29* | *0,31* | *0,07* | *0,05* | *0,12* | *0,05* | *0,17* | *0,31* |
| | 3R | 0,18 | 0,16 | 0,24 | 0,54 | *0,27* | *0,37* | *0,12* | *0,06* | *0,14* | *0,13* | *0,11* | *0,22* |
| | 3L | 0,29 | 0,16 | 0,27 | 0,57 | *0,32* | *0,23* | *0,02* | *0,02* | *0,23* | *0,13* | *0,12* | *0,31* |
| *An. merus* (embryo) | X | 0,18 | 0,63 | 0,60 | 0,68 | *0,27* | *0,05* | *0,01* | *0,02* | *0,27* | *0,34* | *0,32* | *0,38* |

| | | | | | | | | | | | | | |
|---|---|---|---|---|---|---|---|---|---|---|---|---|---|
| | 2R | 0,43 | 0,55 | 0,55 | 0,54 | *0,21* | *0,23* | *0,05* | *0,06* | *0,39* | *0,29* | *0,32* | *0,34* |
| | 2L | 0,30 | 0,49 | 0,48 | 0,50 | *0,56* | *0,04* | *0,04* | *0,05* | *0,27* | *0,31* | *0,31* | *0,31* |
| | 3R | 0,23 | 0,50 | 0,47 | 0,48 | *0,36* | *0,28* | *0,12* | *0,06* | *0,19* | *0,22* | *0,17* | *0,22* |
| | 3L | 0,27 | 0,48 | 0,36 | 0,49 | *0,46* | *0,06* | *0,01* | *0,07* | *0,19* | *0,27* | *0,18* | *0,30* |
| *An. merus* (adult) | X | 0,71 | 0,71 | 0,69 | 0,66 | 0,10 | 0,09 | 0,02 | 0,02 | 0,57 | 0,56 | 0,53 | 0,49 |
| | 2R | 0,62 | 0,61 | 0,60 | 0,58 | 0,01 | 0,04 | 0,03 | 0,02 | 0,61 | 0,59 | 0,57 | 0,53 |
| | 2L | 0,67 | 0,01 | 0,15 | 0,50 | 0,18 | 0,12 | 0,02 | 0,06 | 0,61 | 0,03 | 0,08 | 0,47 |
| | 3R | 0,66 | 0,66 | 0,58 | 0,58 | 0,13 | 0,11 | 0,07 | 0,05 | 0,54 | 0,53 | 0,45 | 0,46 |
| | 3L | 0,60 | 0,04 | 0,08 | 0,53 | 0,09 | 0,01 | 0,03 | 0,04 | 0,58 | 0,06 | 0,02 | 0,50 |
| *An. stephensi* | X | 0,03 | 0,73 | 0,70 | 0,69 | 0,04 | 0,46 | 0,45 | 0,42 | 0,07 | 0,52 | 0,52 | 0,53 |
| | 2R | 0,05 | 0,12 | 0,10 | 0,66 | 0,38 | 0,10 | 0,02 | 0,40 | 0,02 | 0,05 | 0,13 | 0,49 |
| | 2L | 0,10 | 0,01 | 0,03 | 0,53 | 0,09 | 0,11 | 0,01 | 0,41 | 0,13 | 0,06 | 0,07 | 0,38 |
| | 3R | 0,61 | 0,63 | 0,55 | 0,62 | 0,49 | 0,46 | 0,33 | 0,36 | 0,39 | 0,37 | 0,31 | 0,39 |
| | 3L | 0,48 | 0,63 | 0,54 | 0,60 | 0,60 | 0,54 | 0,36 | 0,39 | 0,37 | 0,45 | 0,40 | 0,43 |

**Supplementary Table 6.** Properties of TADs.

| | *An. coluzzii* | *An. atroparvus* | *An. merus* | *An. stephensi* | *An. albimanus* |
|---|---|---|---|---|---|
| **Min length** | 10 000 | 20 000 | 10 000 | 15 000 | 25 000 |
| **Max length** | 1 055 000 | 870 000 | 1 640 000 | 650 000 | 660 000 |
| **Median length** | 145 000 | 130 000 | 185 000 | 135 000 | 135 000 |
| **Mean length** | 173 393 | 151 749 | 236 575 | 155 902 | 155 133 |
| **Number of TADs** | 1 326 | 1 442 | 984 | 1 252 | 1 084 |

**Supplementary Table 7.** Gene orthologous located within the X- and A-loop anchors.

X-loop:

| **Molecular function** | ***An. coluzzii*** | ***An. stephensi*** | ***An. albimanus*** | ***An. atroparvus*** | ***An. merus*** |
|---|---|---|---|---|---|
| Cytochrome P450 CYP9K1 | **AGAP000818** | **ASTEI01072** | AALB003283* | **AATE016607** | **AMEM003066** |
| nuclear receptor subfamily 2 group E member (Tailless) | **AGAP000819** | **ASTEI01071** | AALB003284* | **AATE006814** | **AMEM005434** |
| ANK_REP_REGION domain-containing protein | **AGAP000413** | **ASTEI06259** | AALB007797* | AATE005968 | **AMEM017290** |
| prostaglandin reductase 1 | **AGAP000414** | **ASTEI06258** | AALB006758 | **AATE002874** | **AMEM010926** |
| hydroxyisourate hydrolase activity | **AGAP000415** | **ASTEI06257** | AALB006759 | **AATE016248** | **AMEM011098** |
| Non-specific serine/threonine protein kinase | **AGAP029239** | **ASTEI01297** | AALB006314 | **AATE016354** | **AMEM010384** |
| integrin beta subunit | **AGAP000815** | ASTEI07790 | AALB006392 | **AATE000443** | **AMEM009509** |
| metal ion binding | **AGAP029244** | **ASTEI06256** | AALB006313 | **AATE018169** | **AMEM014919** |
| integral component of membrane | **AGAP000420** | **ASTEI06255** | AALB006411 | **AATE017335** | AMEM017711 |
| (heparan sulfate)-glucosamine 3-sulfotransferase 3 | **AGAP000422** | **ASTEI06254** | AALB006570 | **AATE004104** | **AMEM010094** |
| Frataxin homolog, mitochondrial | **AGAP000813** | ASTEI07792 | AALB009856 | **AATE002095** | AMEM001133 |
| phosphatidylinositol binding | **AGAP000814** | ASTEI07791 | AALB006417 | **AATE012817** | - |

A-loop:

| **gene function** | ***An. coluzzii Ngousso*** | ***An. stephensi*** | ***An. albimanus*** | ***An. atroparvus*** | ***An. merus*** |
|---|---|---|---|---|---|
| ubiquitin thioesterase ZRANB1 | **<u>AGAP003853</u>** | **<u>ASTEI00780</u>** | **<u>AALB003080</u>** | **<u>AATE000898</u>** | **<u>AMEM006320</u>** |
| snRNA-activating protein complex subunit 3 | **<u>AGAP003852</u>** | **<u>ASTEI00779</u>** | **<u>AALB003081</u>** | **<u>AATE008621</u>** | **<u>AMEM002040</u>** |
| 26 proteasome complex subunit | **<u>AGAP003851</u>** | **<u>ASTEI00778</u>** | **<u>AALB003082</u>** | **<u>AATE007015</u>** | - |
| signal peptide | **<u>AGAP013134</u>** | **<u>ASTEI00777</u>** | **<u>AALB003083</u>** | **<u>AATE018674</u>** | **<u>AMEM017317</u>** |
| DNA-binding transcription factor activity | **<u>AGAP003726</u>** | **<u>ASTEI00671</u>** | AALB007121 | **<u>AATE016879</u>** | **<u>AMEM010172</u>** |
| integral component of membrane | **<u>AGAP003849</u>** | ASTEI00776 | AALB003084 | **<u>AATE004695</u>** | **<u>AMEM017419</u>** |

Orthologous are marked in underline/bold if they are located in the loop anchors. Genes, which are located outside of the loop anchors, but closer than 1 Mb from each other, are marked with an asterisk (*).

**Supplementary Table 8.** Genomic coordinates of long-range X- and A-loop and H3K27me3-associated (Polycomb) loop in *An. atroparvus*.

| Loop type | chr | start_1 | end_1 | start_2 | end_2 |
|---|---|---|---|---|---|
| Long-distance_loop | X | 5,320,000 | 5,755,000 | 11,100,000 | 11,510,000 |
| Long-distance_loop | X | 630,000 | 780,000 | 10,000,000 | 10,500,000 |
| Long-distance_loop | 3R | 7,300,000 | 8,200,000 | 38,550,000 | 39,050,000 |
| Long-distance_loop | 2R | 11,260,000 | 11,690,000 | 23,130,000 | 23,400,000 |
| Polycomb_loop | X | 5,740,000 | 5,770,000 | 6,130,000 | 6,150,000 |
| Polycomb_loop | X | 5,790,000 | 5,815,000 | 6,130,000 | 6,150,000 |
| Polycomb_loop | X | 9,480,000 | 9,500,000 | 9,620,000 | 9,635,000 |
| Polycomb_loop | X | 13,470,000 | 13,490,000 | 13,575,000 | 13,595,000 |
| Polycomb_loop | X | 13,500,000 | 13,515,000 | 13,575,000 | 13,595,000 |
| Polycomb_loop | X | 16,405,000 | 16,425,000 | 16,495,000 | 16,510,000 |
| Polycomb_loop | 2R | 16,600,000 | 16,630,000 | 16,710,000 | 16,730,000 |
| Polycomb_loop | 2R | 16,600,000 | 16,630,000 | 16,800,000 | 16,825,000 |
| Polycomb_loop | 2R | 19,675,000 | 19,680,000 | 19,765,000 | 19,790,000 |
| Polycomb_loop | 2R | 22,090,000 | 22,115,000 | 22,180,000 | 22,200,000 |
| Polycomb_loop | 2R | 22,535,000 | 22,560,000 | 22,640,000 | 22,655,000 |
| Polycomb_loop | 2R | 24,945,000 | 24,960,000 | 25,025,000 | 25,050,000 |
| Polycomb_loop | 2R | 35,980,000 | 36,005,000 | 36,130,000 | 36,160,000 |

**Supplementary Table 9.** Datasets used for evolutionary comparison of slope-plots.

| **Dataset** | **Source** |
|---|---|
| ***Anopheles* species** | |
| Five *Anopheles* species | This work |
| **Other insects** | |
| *Aedes aegypti* | [8] |
| *Drosophila melanogaster* (embryo) | [9] |
| *Drosophila busckii* (embryo) | [9] |
| *Drosophila viridis* (embryo) | [9] |
| *Drosophila melanogaster* (nc1-4, nc12, nc13, 3-4h embryo) | [10] |
| *Drosophila melanogaster* Kc167 cells | [11],[12] |
| *Drosophila melanogaster* S2 cells | [13],[14] |
| *Drosophila melanogaster* BG3 | [15] |
| *Drosophila melanogaster* salivary glands (Polytene) | [16] |
| *Drosophila melanogaster* Rad21/CapH2 knockdowns | [6] |
| **Chicken** | |
| Erythrocytes and fibroblasts | [4] |
| DT40 at various cell cycle stages, including cells with condensin subunits depleted | [17] |
| **Mammals** | |
| Mouse neural progenitors, embryonic stem cells, mouse cortical neurons | [18] |
| Mouse rod photoreceptors, thymus WT and LBR-/- cells | [19] |
| Human monocytes | [20] |
| Human HCT116 with/without RAD21 degron | [3] |
| Human HAP1 with/without WAPL | [5] |
| Human embryonic stem cells, human fibroblasts | [21] |

**Supplementary Table 10.** Set of primers used for PCR product amplification in FISH experiments.

| Primer name | Primer sequence |
|---|---|
| atr16879_ex2_F | ACGTGAAGGAAGAACCGCTACAG |
| atr16879_ex2_R | AATGGGTAGAACTCGTCCATCATTT |
| atr16879_ex7_1_F | TCAGCTTGTTTGGGGTATGAACTTA |
| atr16879_ex7_1_R | CTCTACCCTACGCTCTCTTCCTTTG |
| atr16879_ex7_2_F | CCCGTACACATATGAACCAAAACAT |
| atr16879_ex7_2_R | AAGCACAGCTATTACAGTGGTACGC |
| atr15805_ex2_F | ACAGGCTAAATATCACTGCATCGAG |
| atr15805_ex2_R | ATCCGTACACGTCAACGTTTCTAAC |
| atr8621_ex1_F | TACAAGCCAAAATCGAATAAGGTCA |
| atr8621_ex1_R | TCGTAATACGGTCCTTCATCACAGT |
| atr0898_ex2_F | GAAATCCTCTCCACCCAGAGTAAAA |
| atr0898_ex2_R | TGATGTTGCTGGCTAGGTTGATATT |
| aate009242_ex9_pr F | CGAGTAGTGGCCGGATACAAG |
| aate009242_ex9_pr R | GCTTGATTTCCAGATTCAGCAG |
| aate008403_ex6_pr F | CTGGATCGCATTCTACATCTGC |

| | |
|---|---|
| aate008403_ex6_pr R | GCCATTCCATGTTCACGTCTTA |
| aate013841_ex11_pr F | ACGTTGTCTTCACCTGGCACTA |
| aate013841_ex11_pr R | CCCAAACGGGATACTCCAATAG |
| aate019780_ex5_pr F | ACAAACTGCGGGACTCTCTGAT |
| aate019780_ex5_pr R | GTGATTCGTTCCTTGTCCCACT |
| aate009704_ex2_pr F | ATAGTGGTCACCAGGCCAATTT |
| aate009704_ex2_pr R | TTTGGCTTCTTCCAAACCTAGC |
| aate015765_F ex3 | GGTCAACCAGTTCTGGATAATGT |
| aate015765_R ex3 | AGCCGAGGGTGAATATTTGATA |
| aate015765_F ex4 | CTTCGTCCCGACTATGACTAGC |
| aate015765_R ex4 | AGAAAGGGAACGTACAGCTTGA |
| aate015765_F ex7 | ACCGTACTGCGTCTATCCAACT |
| aate015765_R ex7 | TACTCGTTTGAATGTGGCATCT |
| aate012799_F ex18 | CGACTTCCGCAGTAATACATGA |
| aate012799_R ex18 | TCGACATCCGAAACAGTAGTTG |
| aate000335_F ex5 | GGCTGGTAAAGGATCTTAGCAA |
| aate000335_R ex5 | TACACTGAGCTGTGGTACATGG |

| | |
|---|---|
| aate000335_F ex5_2 | CCATGTACCACAGCTCAGTGTAT |
| aate000335_R ex5_2 | GGCTCATTTGAGGAGGACACT |
| ste0671_ex1_F | GCCTTAACTACACCGGTATCGACTG |
| ste0671_ex1_R | ACGAGGGTGAAATTAAATTGTTCCA |
| ste0671_ex2_1_F | ATCGATGGTTTGTTTCACGATGAT |
| ste0671_ex2_1_R | AGTAAATTCTTCGCTCCGGTAGAGC |
| ste0671_ex2_2_F | CTCTACCGGAGCGAAGAATTTACTC |
| ste0671_ex2_2_R | CACCGCTCGAATAATCTTCATAGTG |
| ste0780_ex2_1_F | GATCGAACCCAAATCTAGCAAGC |
| ste0780_ex2_2_R | TACGTACAAAGCGAGCAGTACCATT |
| ste0780_ex2_1_F | GGCCAGTAACATCATGGTAAACAAC |
| ste0780_ex2_2_R | AGTAATTTCCAGCGACCAGTTCAAT |
| ste0779_ex1_F | CATTTACCTACCAAGAACGGAGCAG |
| ste0779_ex1_R | ATTTTGTCCTTCAGCTCGGTTAAGT |
| acol000412_F ex3 | CTGCCAGATACTGATGAAGTGC |
| acol000412_R ex3 | TACGTGTCGTGTATGAGCGACT |
| acol000413_F ex5 | GATCTGAACCTCACCACCTACC |

| | |
|---|---|
| acol000413_R ex5 | GTGCTAACCATCACGATGTCCT |
| acol000415_F ex1 | CACCAAAGCCGTCAACATCTC |
| acol000415_R ex1 | GATGAACGGGTACAGGCTTTC |
| acol000815_F ex5 | ACAACGTGATCGACGTGAAGTA |
| acol000815_R ex5 | AGAAGTTGTCGCACTCGCAGTA |
| acol000815_2F ex5 | GTGCAACGAGTTCAAGCACTG |
| acol000815_2R ex5 | GTGTCCCACTTTGCCATCAT |
| acol000815_F ex1_2 | TGTCCAACTATCAGCTCACGAC |
| acol000815_R ex1_2 | GATCGTCTTATCGTCCTCCATC |

### Supplementary figures

**Supplementary Figure 1.** Polymorphic inversions in *Anopheles* species visualized by Hi-C.

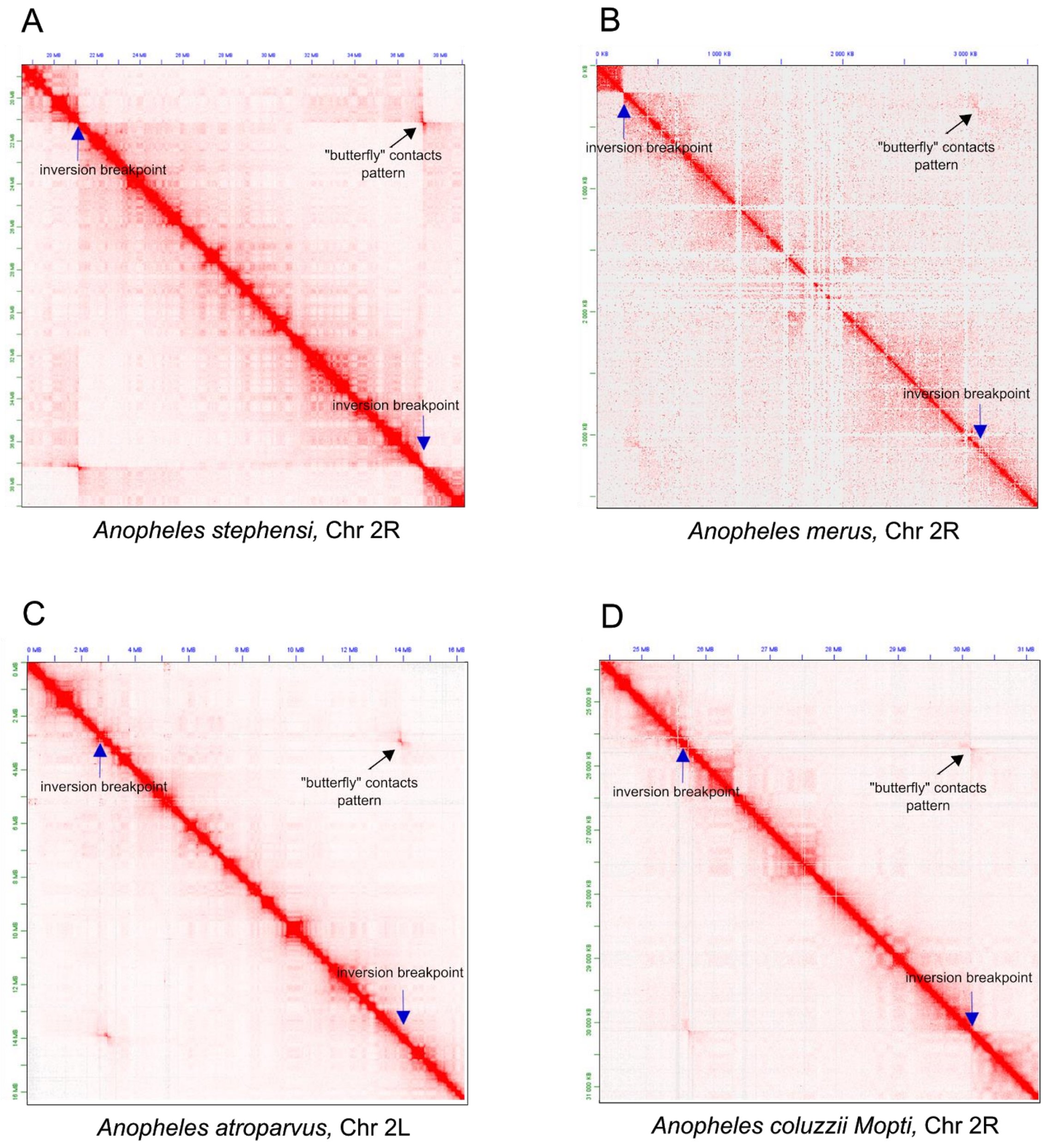


A) *Anopheles stephensi*, inversion coordinates: Chr 2R, 21,140,000 - 37,180,000; B) *Anopheles merus*, inversion coordinates: Chr 2R, 60,175,000-62,930,000; C) *Anopheles atroparvus*, inversion coordinates: Chr 2L, 2,980,000 - 13,940,000; D) *Anopheles coluzzii Mopti*, inversion coordinates: Chr 2R, 30,940,000 - 35,340,000. Blue arrow points the typical “butterfly” contacts pattern.

**Supplementary Figure 2.** Analysis of Hi-C contacts demonstrates signatures of Rabl-like chromosome configuration.

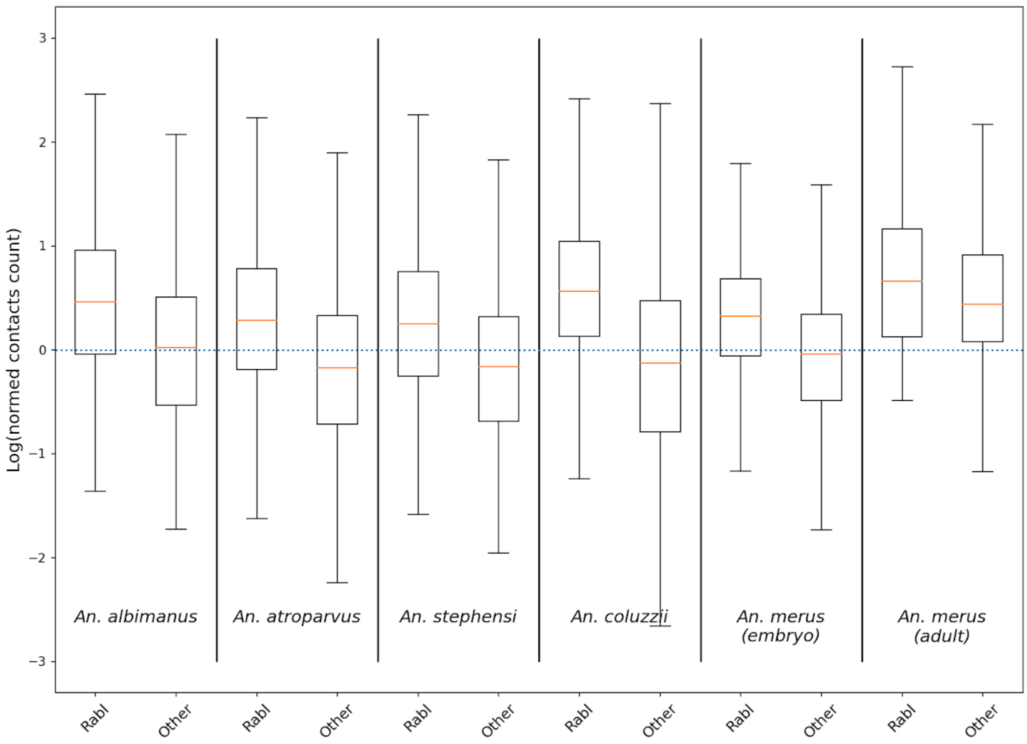


Box-plots represent a distribution of logarithm of distance-normalized contact frequencies. We define “Rabl contacts” (label “Rabl”) as contacts of loci located on two different arms of same chromosome at similar distance from centromere (difference in distance <100 kb); and other contacts (label “other”), defined as contacts of loci located on two different arms of same chromosome at different distance from centromere (difference in distance >300 kb). For distance normalization, we used expected values calculated by juicer.

**Supplementary Figure 3.** Large heterochromatic blocks located on Chr X were observed in all *Anopheles* species in the experiment.

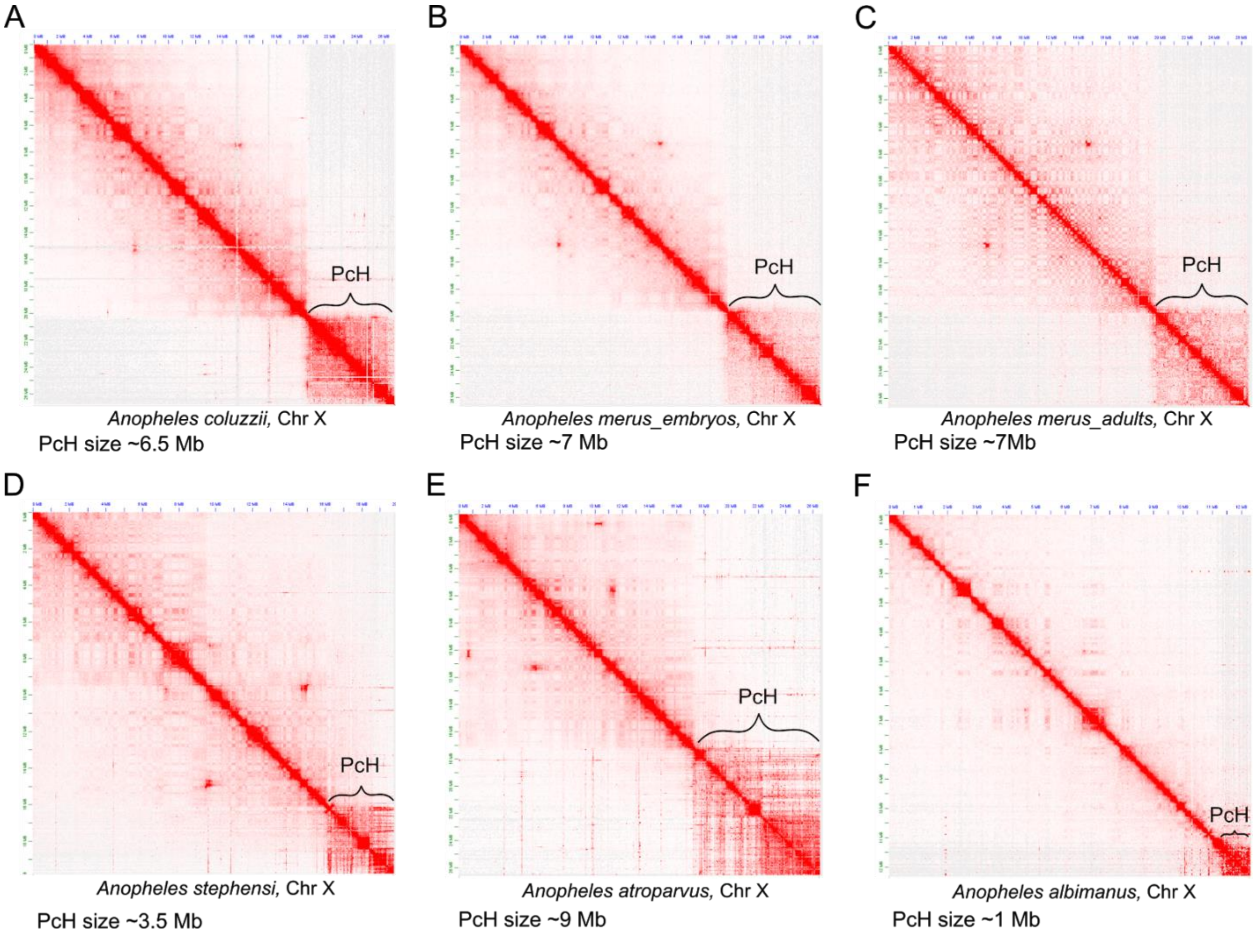


A. Hi-C heatmap for Chr X, *Anopheles coluzzii*. B. Hi-C heatmap for Chr X, *Anopheles merus* embryos. C. Hi-C heatmap for Chr X, *Anopheles merus* adults. D. Hi-C heatmap for Chr X, *Anopheles stephensi*. E. Hi-C heatmap for Chr X, *Anopheles atroparvus*. F. Hi-C heatmap for Chr X, *Anopheles albimanus*.

**Supplementary Figure 4.** Correspondence between pre-centromeric and intercalary heterochromatic blocks on Hi-C and cytological maps.

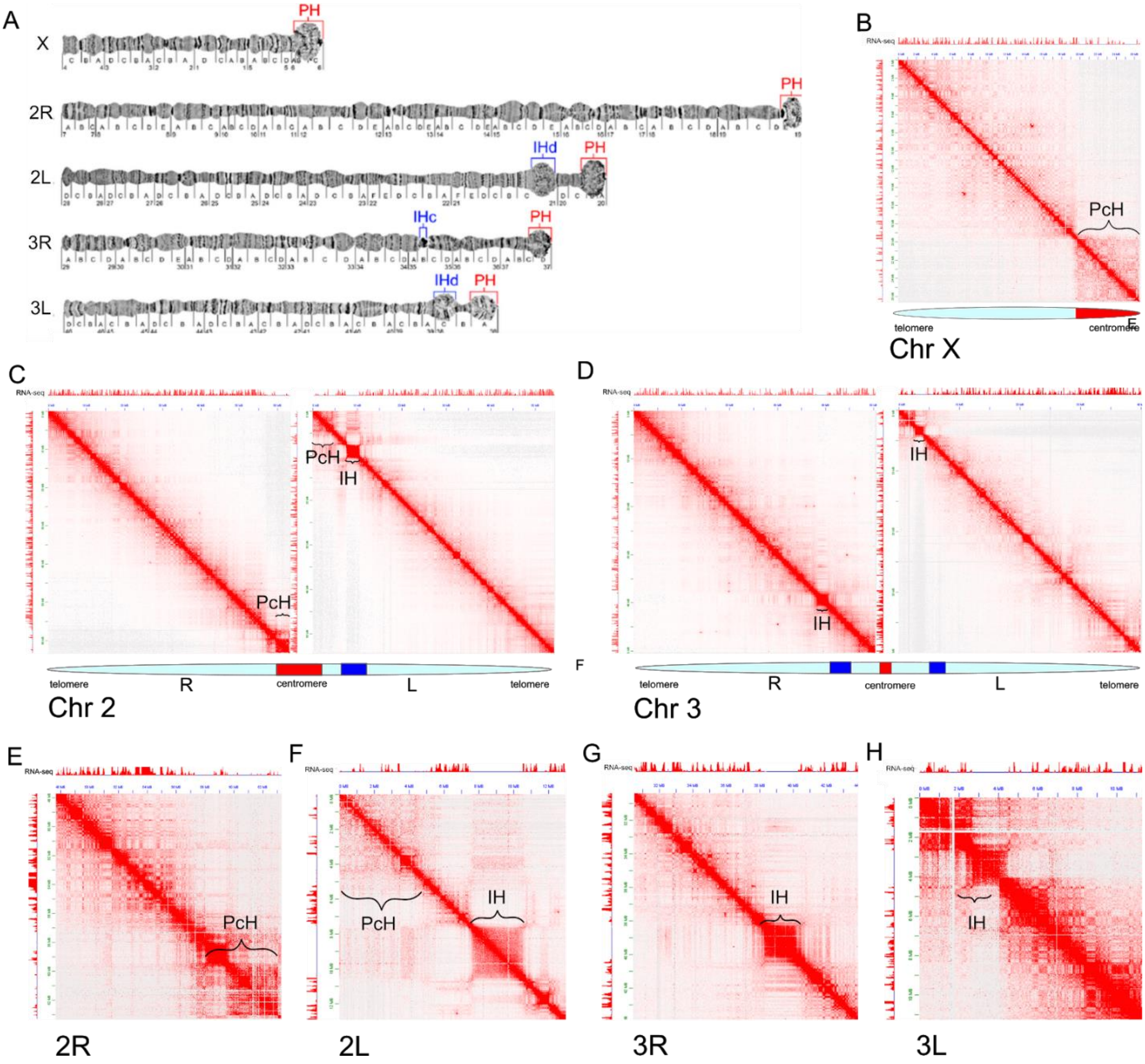


A. Physical map for *An.gambiae* taken from Sharakhova et al.[22]. B-D. Correspondence between pre-centromeric and intercalary heterochromatic blocks on Hi-C map for *An.merus* genome and standard cytogenetic map, shown for chromosomes X, 2 and 3. E-F. Zoomed-in images demonstrating reduction in RNA-seq signal for heterochromatic regions.

**Supplementary Figure 5.** Distribution of PC1 values on chromosomal arms for five *Anopheles* species.

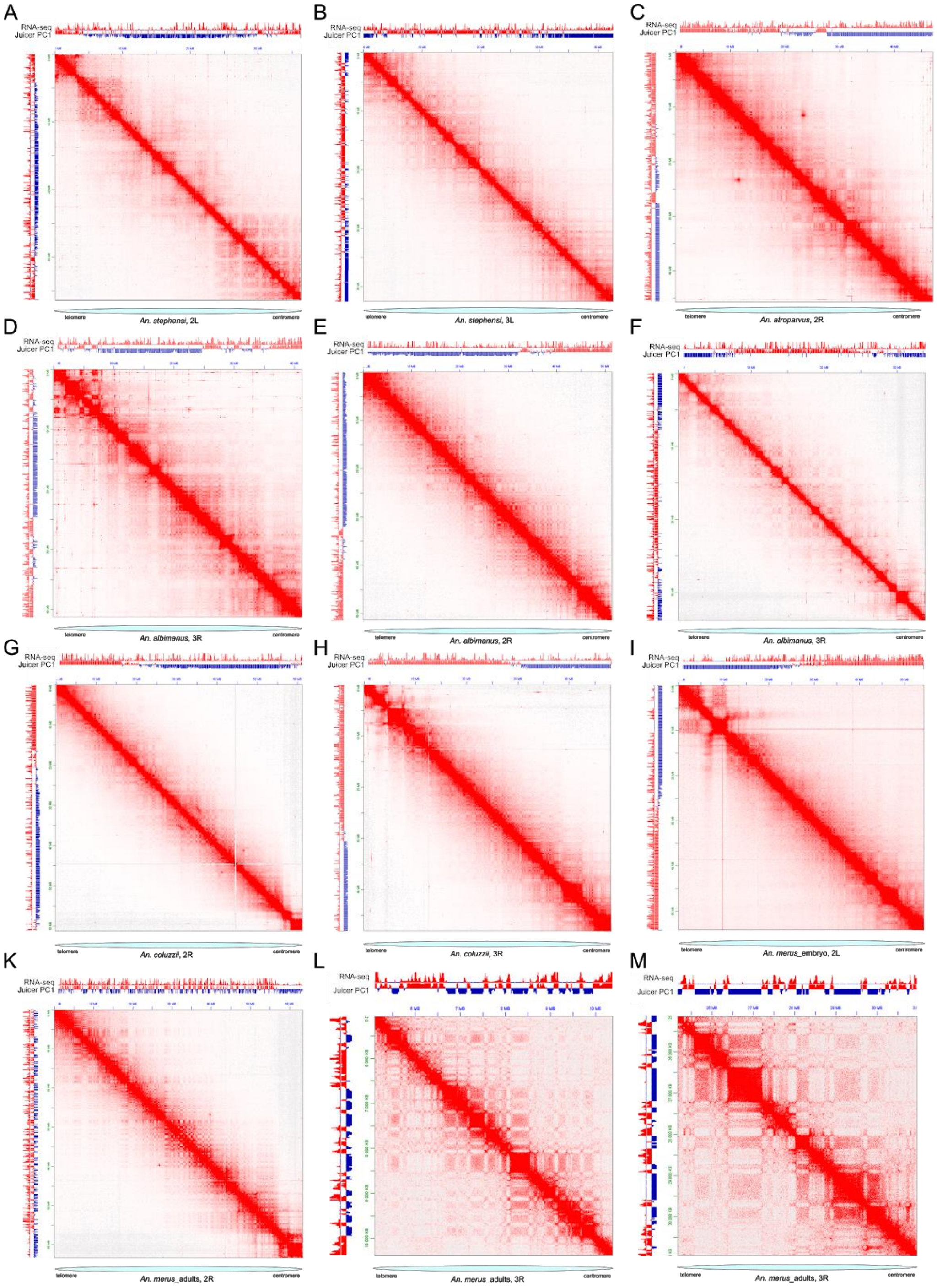


A-I. Hi-C heatmaps for *Anopheles* embryos. The first principal component (PC1) of the Hi-C matrix does not reflect observed plaid-pattern and does not correspond to RNA-seq data for majority of the chromosomes. K-M. Hi-C heatmaps for *An. merus* adults. Standard PC1 (Juicer) algorithm was able to define compartments that agree with plaid-pattern on the majority of *An. merus* chromosomes.

**Supplementary Figure 6.** Transformations of data allowing identification of compartments.

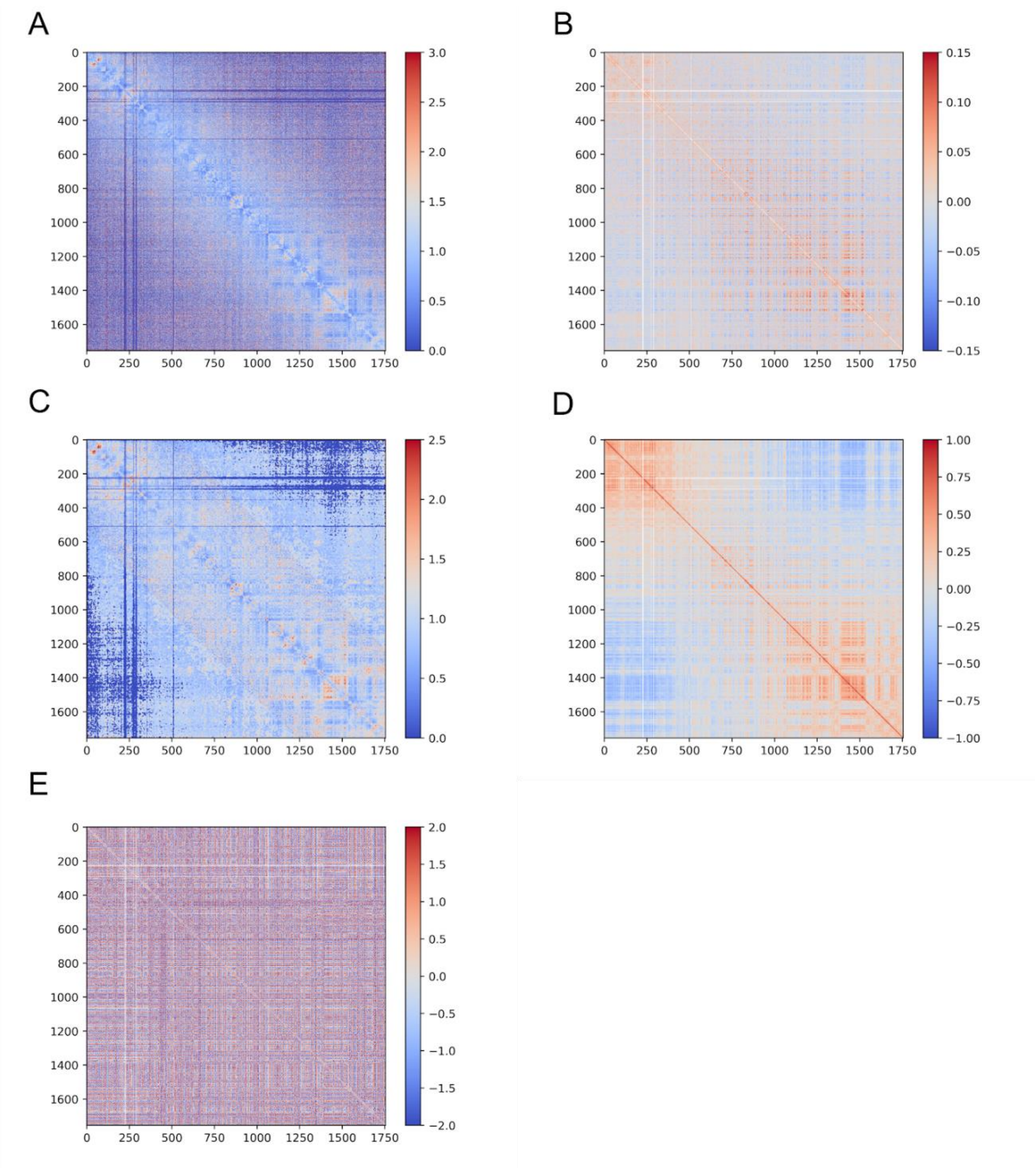


We start from KR-normalized observed/expected matrix (A) and cropped heterochromatic/repetitive regions at each side of chromosome arms. Then, we either directly computed Pearson’s correlation (B) and PC1, which results in “cropped” PC1 track, or performed local averaging for long-range interactions (C), computed Person’s correlation (D) and normalized correlation values within small 10-Mb blocks of obtained matrix, resulting in contrast enhanced data (E). We defined the first principle component of the matrix depicted in panel E as cePC1 values.

**Supplementary Figure 7.** TADs properties.

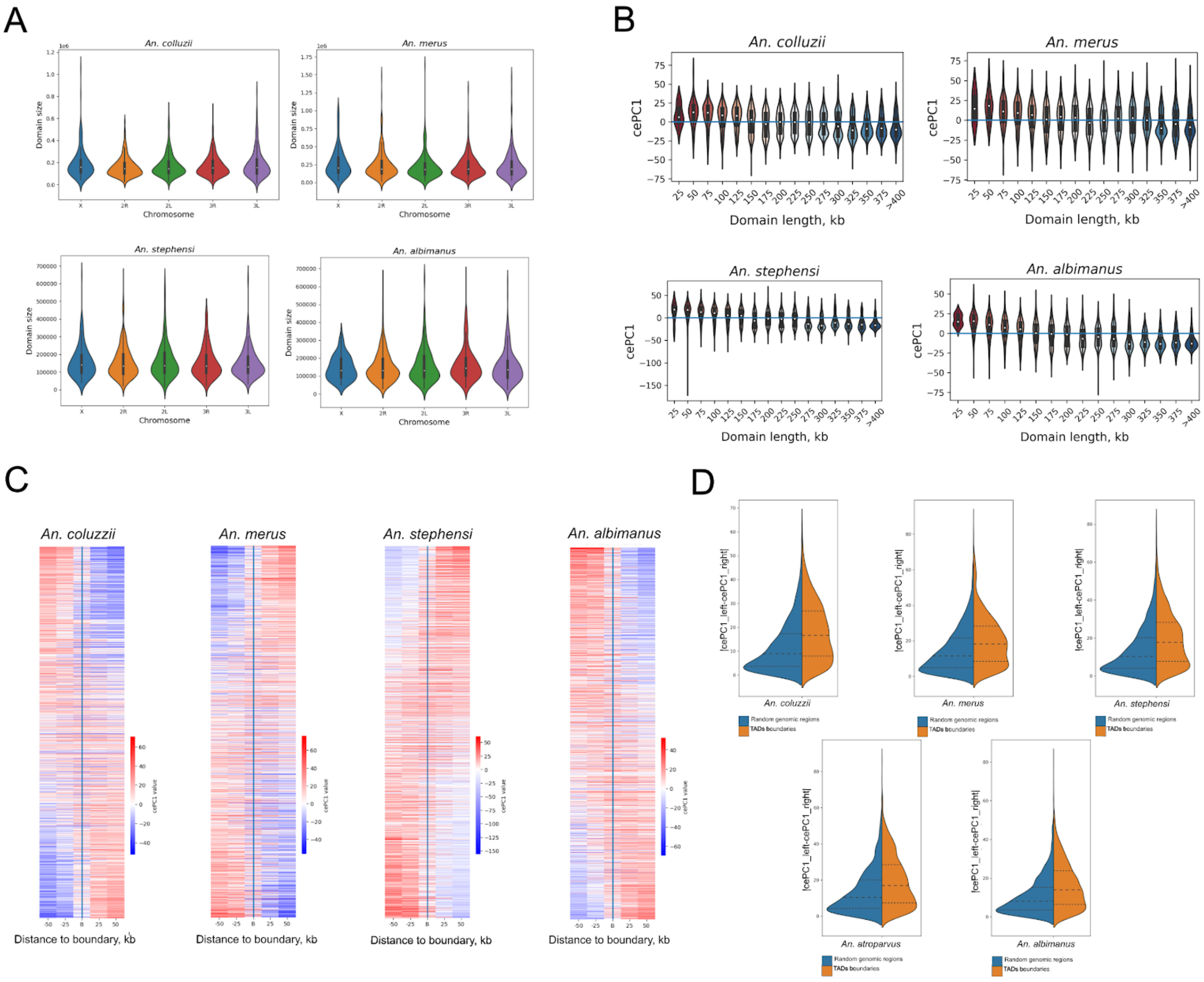


Characteristics of TADs and TAD boundaries provided for four *Anopheles* species (similar data for An. atroparvus shown in Fig. 6). A, B and C represent data as in Fig. 6, C, D and F, respectively. D. Quantification of changes of cePC1 values around TAD boundaries. Changes of cePC1 values around TAD boundaries (red distributions, right) compared to changes of cePC1 values in random genomic positions (blue distributions, left).

**Supplementary Figure 8.** Examples of loops in *An. atroparvus* genome, overlapping (A-E) or non overlapping (F-H) H3K27 histone mark.

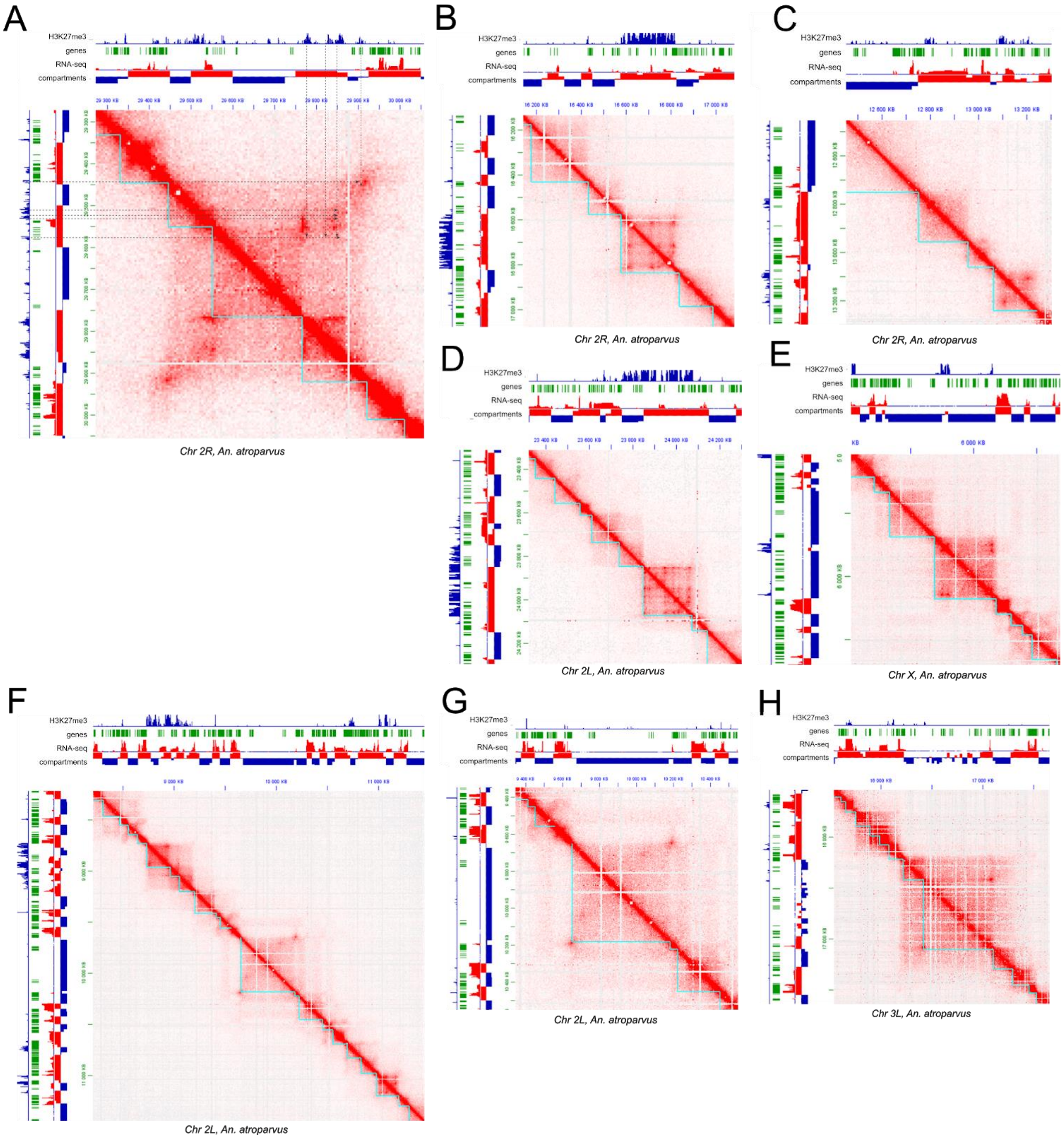

**Supplementary Fig. 9.** A-loop located on arm 2R (or arm 3R in *An. atroparvus*).

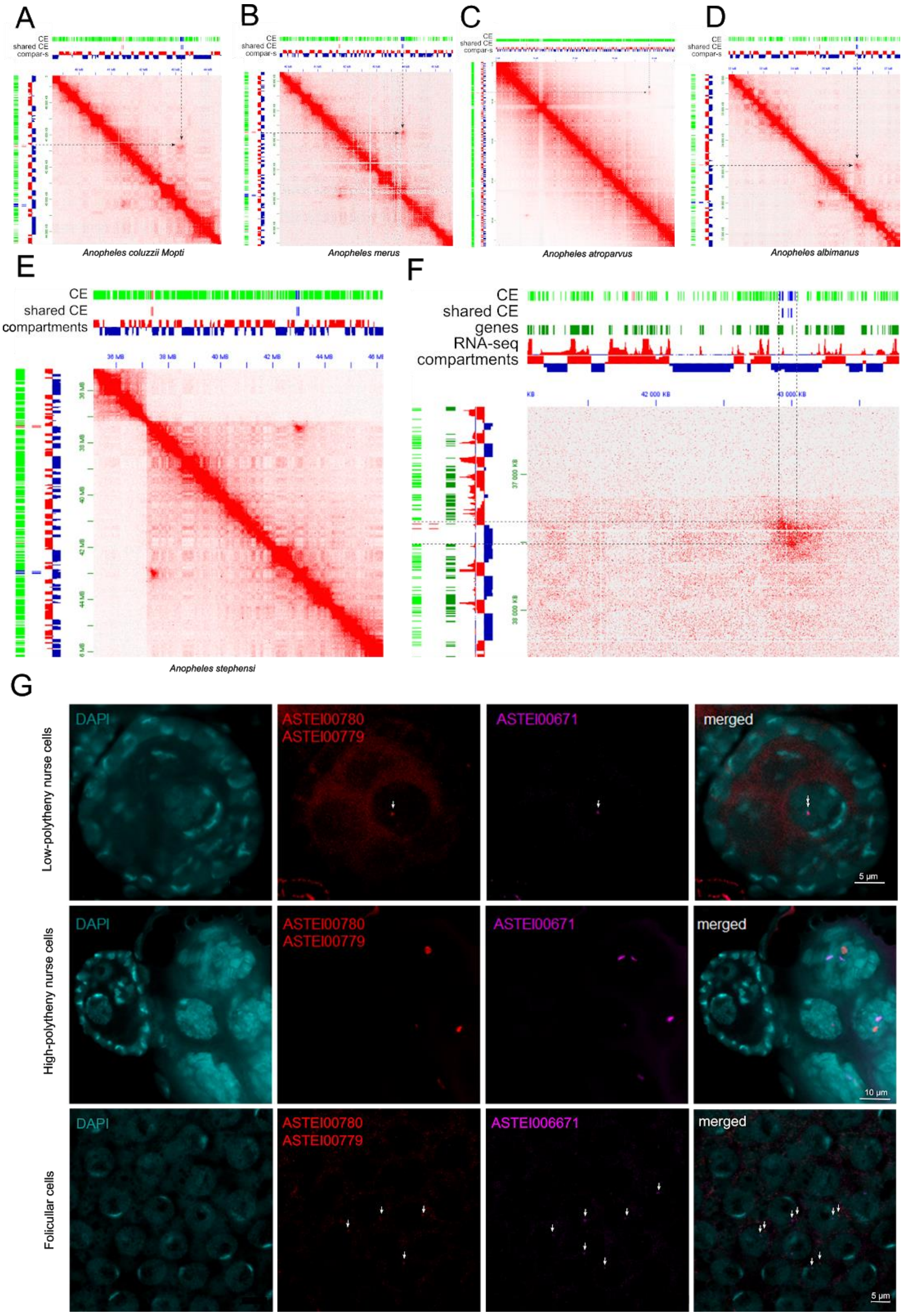


A-E Snapshots of Hi-C maps showing A-loop; F. Zoomed-in fragment of Hi-C map showing A-loop interactions in *An. stephensi*; G. 3D FISH analysis of co-localization of A-loop anchors in nurse cells (chromosomes with low-polytheny and high-polytheny), and follicular cells.

**Supplementary Fig. 10.** Interactions between anchors of long-range loops in *Anopheles* species.

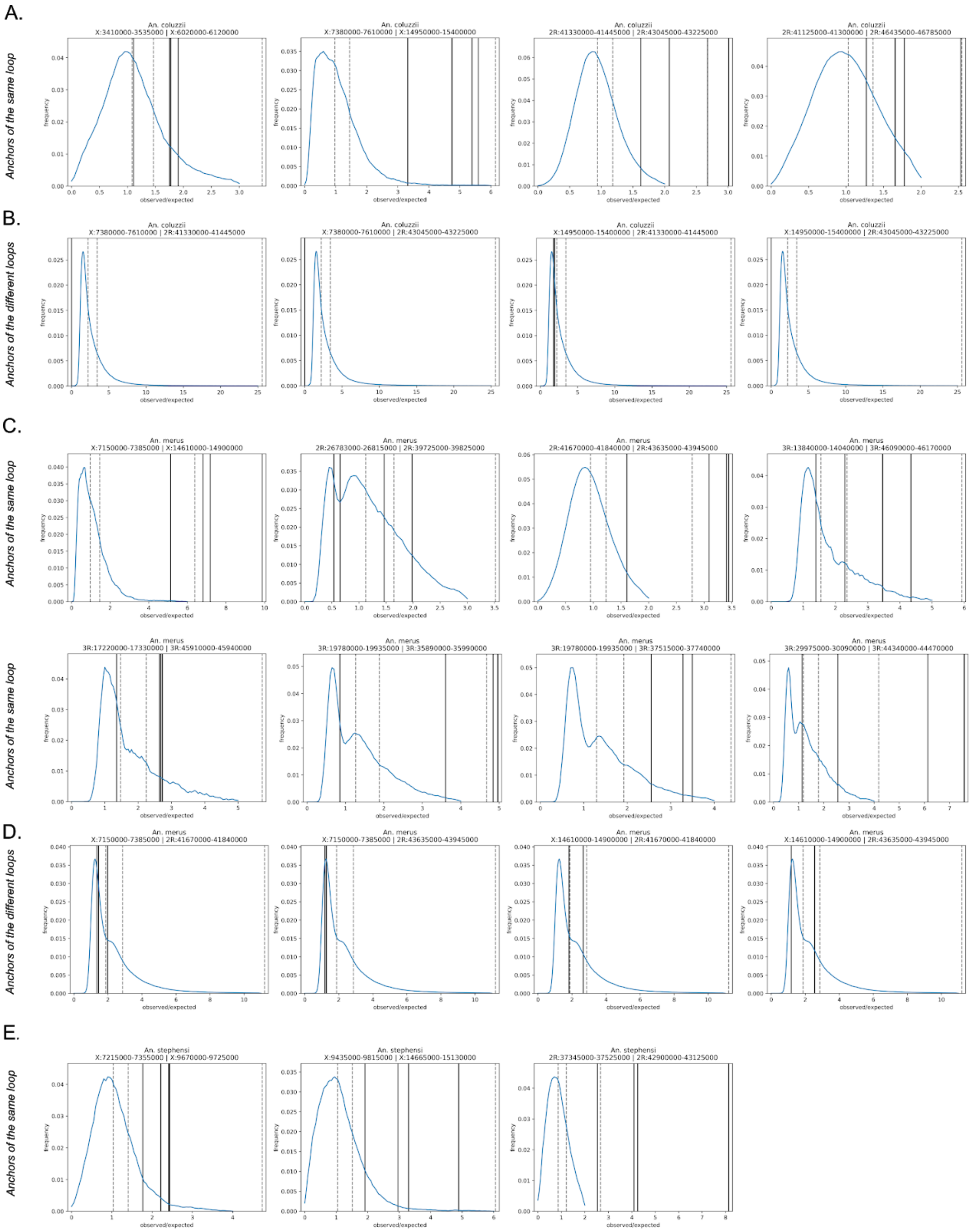

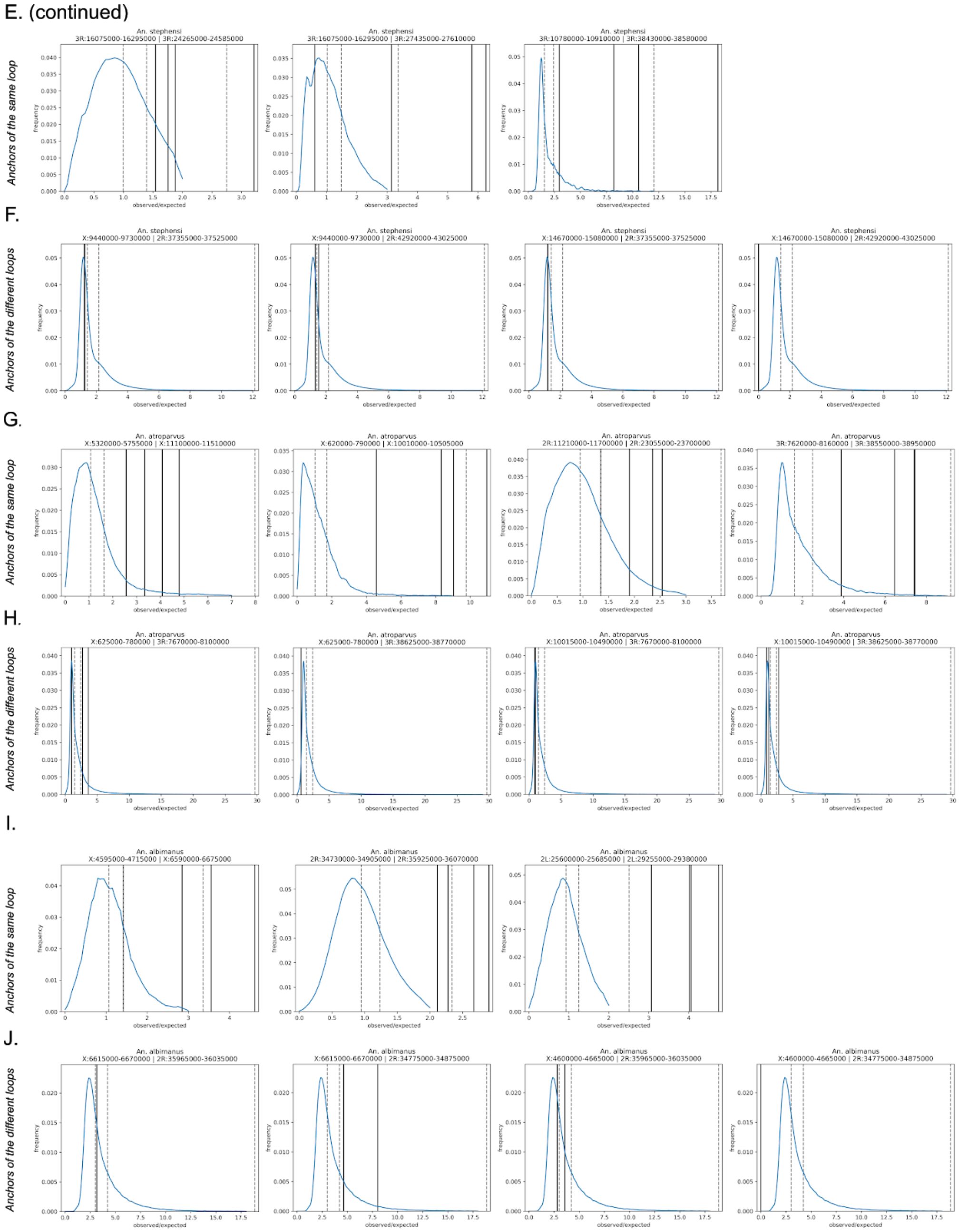


Each graph shows distribution of all observed/expected contact frequency ratios at distance matching distance of loop anchors in blue. We truncated the distribution at 99th percentile, and showed median and third quartile as dashed black vertical lines. Interactions between anchors are shown as black solid vertical lines. When anchors contain more than one bin (results presented for 25-kb binned data) we showed all possible pairwise interaction as separate solid lines. For each species, the first row of graphs show interaction between anchors of one loop, and the next row shows interaction of anchors of different loops.

**Supplementary Figure 11.** Long-range loops are not enriched for the H3K27me3 mark.

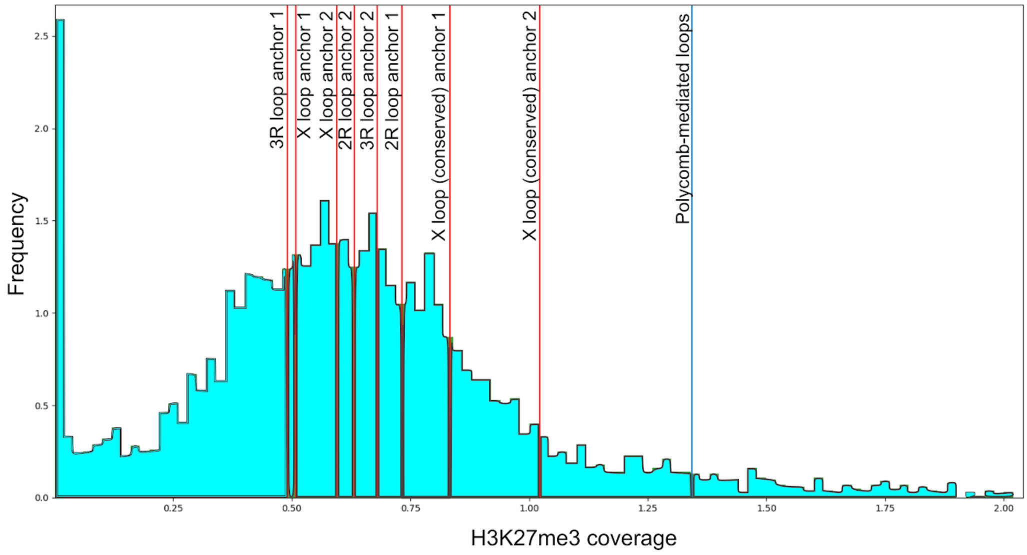


Blue histogram shows a distribution of H3K27me3 reads coverage for *An. atroparvus* genome (fold change over control). Fold change was computed as mean values for 5-kb bins. Red lines – mean of anchor values of A- and X-loops and other long-range loops described in Supplementary Table 3. Blue line - mean read coverage value of H3K27me3-associated (Polycomb) loops taken from Supplementary Table 7.

**Supplementary Figure 12.** The long-distance contact loops are not developmental-stage specific.

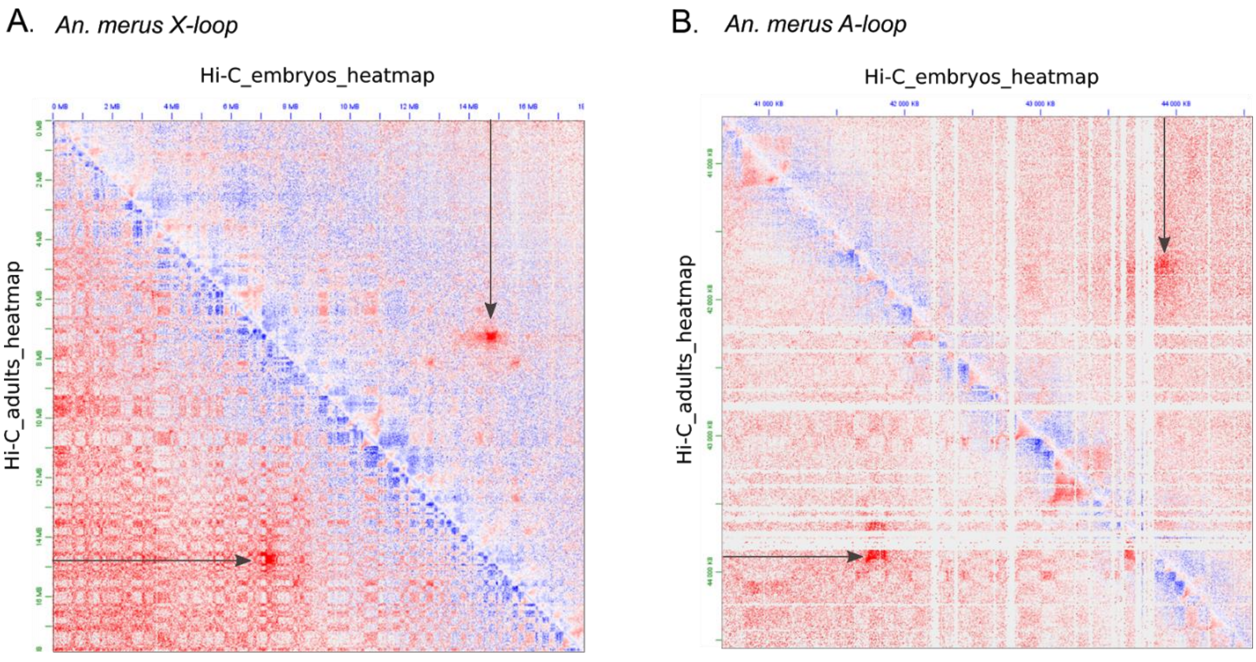


X- (A) and A-loops (B) are present in both adult and embryonic data available for *An. merus.* Shown observed vs control view on adult vs embryo *An. merus* Hi-C map at 25-kb resolution.

**Supplementary Figure 13.** Contact frequency decays non-uniformly with genomic distance.

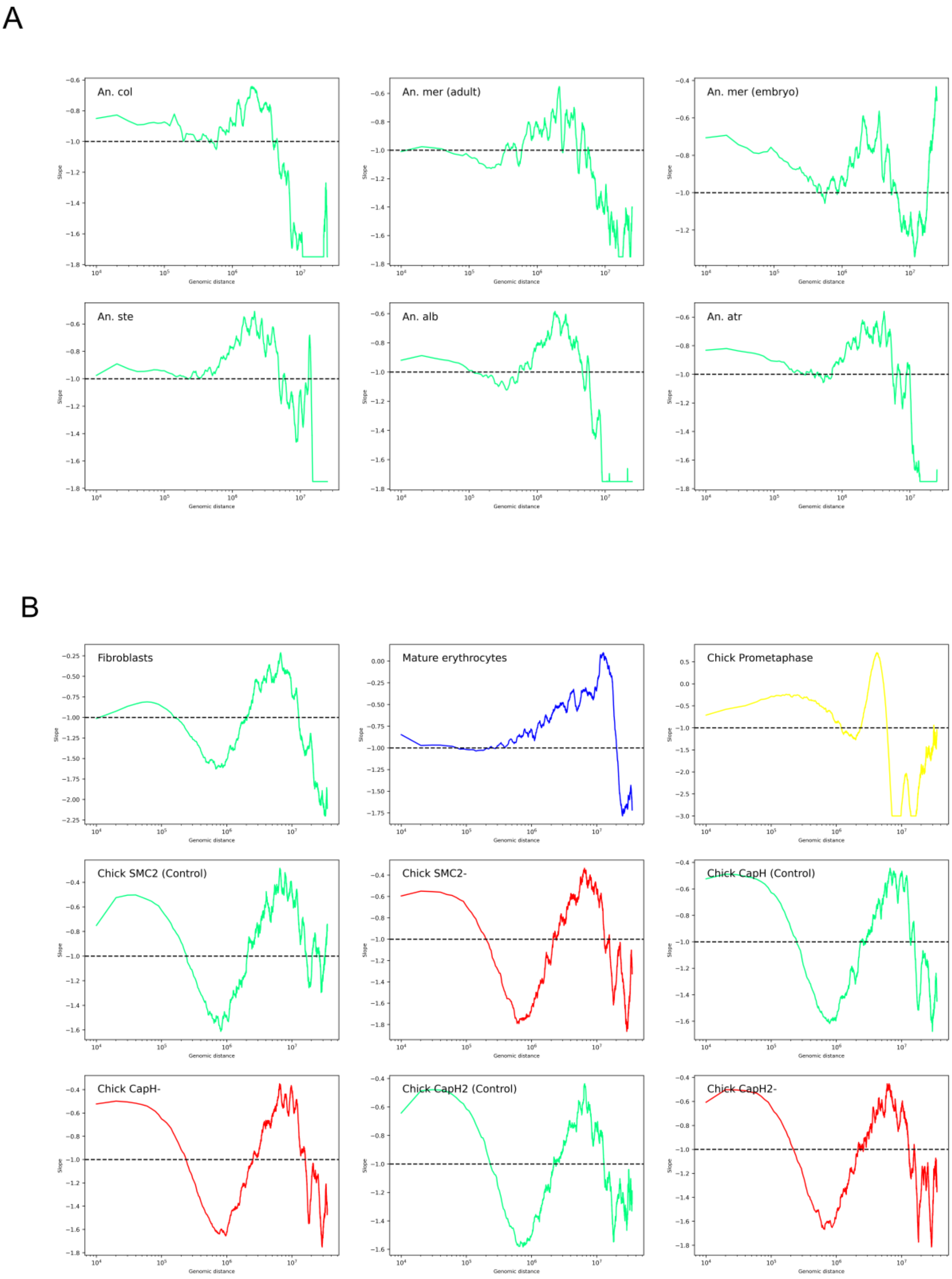

C

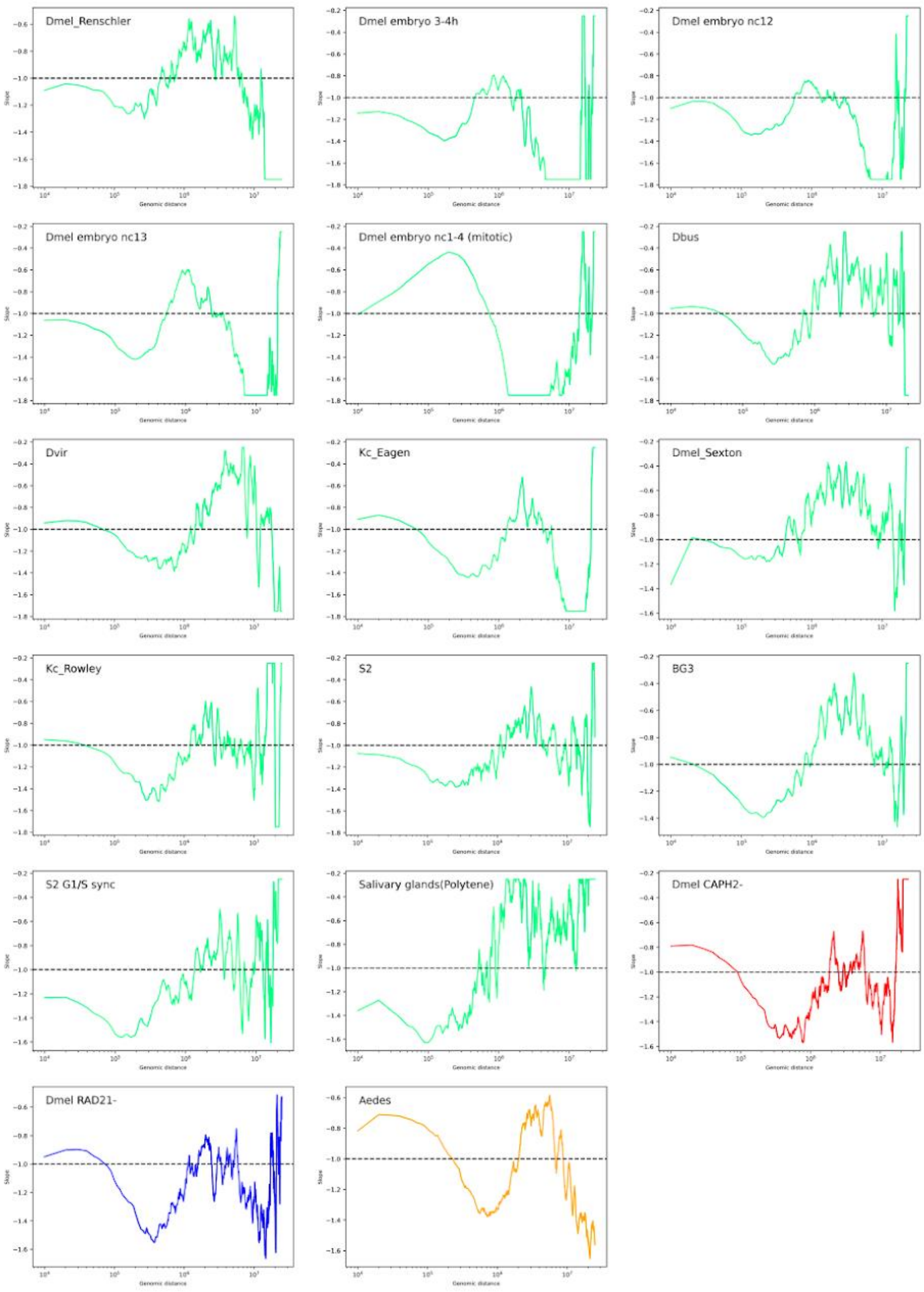

Dmel_Renschler
Dmel embryo 3-4h
Dmel embryo nc12
Dmel embryo nc13
Dmel embryo nc1-4 (mitotic)
Dbus
Dvir
Kc_Eagen
Dmel_Sexton
Kc_Rowley
S2
BG3
S2 G1/S sync
Salivary glands(Polytene)
Dmel CAPH2-
Dmel RAD21-
Aedes

D

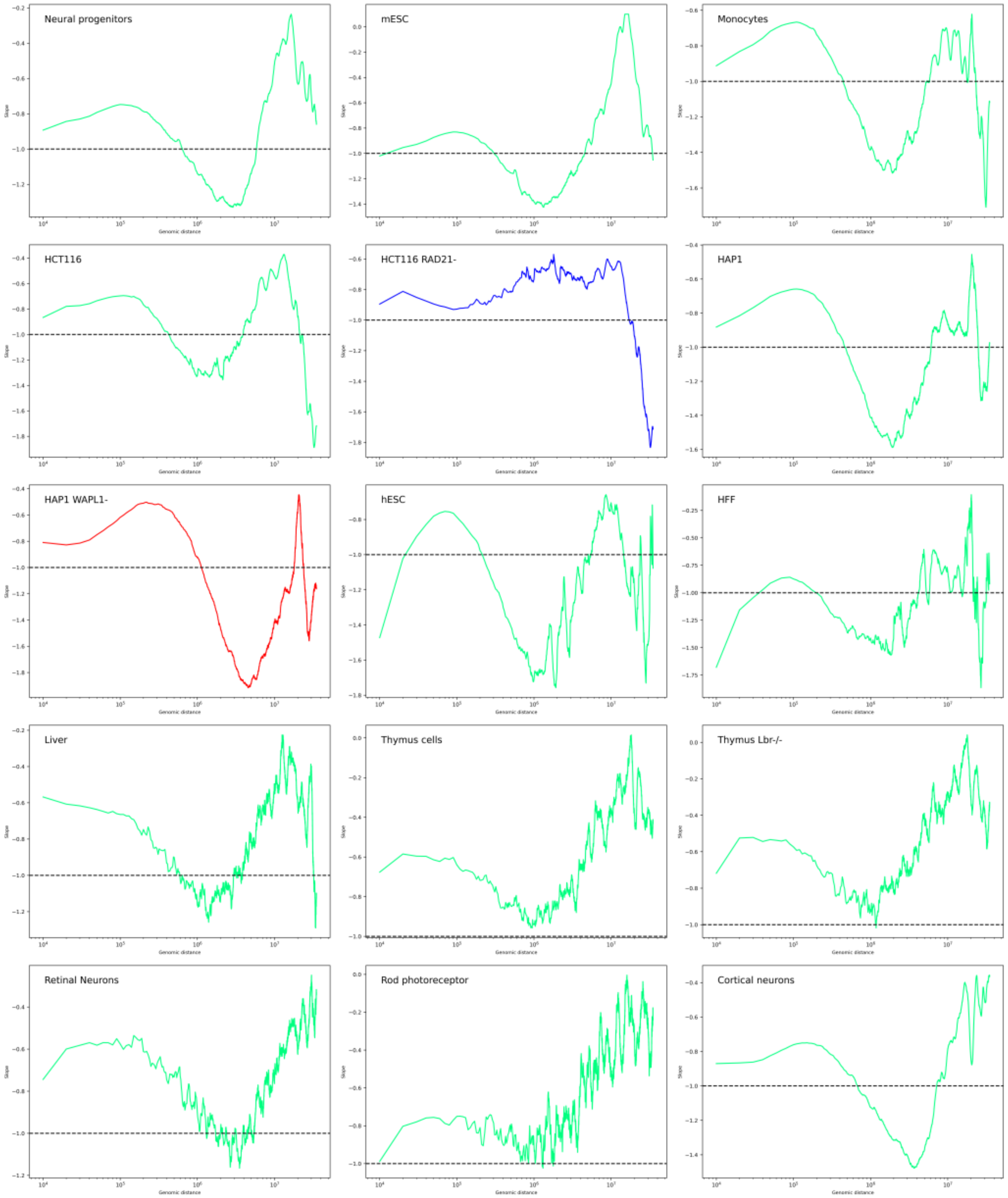


Data is shown as on Fig. 8, B, but for each species independently. *A* shows data for *Anopheles* species, *B* for other insects, *C* for chicken, and *D* for mammals. Sample type and/or species name are indicated on each plot. Plot colors correspond to colors used in Fig. 8.

**Supplementary Figure 14.** Egg dish design specified for collecting mosquito eggs for Hi-C and ChiP seq experiments.

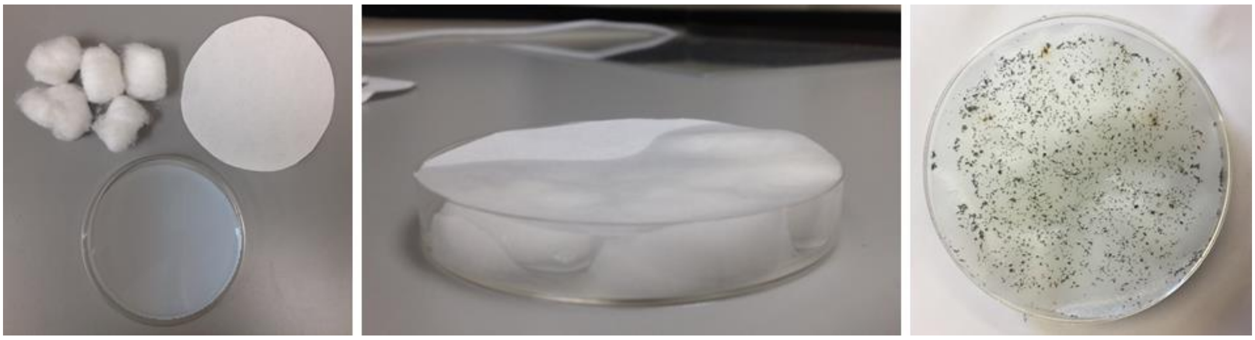

### Supplementary Protocols

#### Supplementary Protocol I

Hi-C experiment

*REAGENTS:*

autoclaved ddH2O
freshly made 50%-bleach/water solution
NaCl, 1M
NaCl, 5M
MgCl, 1M
KCl, 1M
HEPES, 1M (Sigma, H0887)
Glycine, 2M (Sigma, G7126)
Triton, 10% (Sigma, T8787)
Protease Inhibitor Cocktail (Sigma, P8340)
Freshly made PFA, 37% (Sigma, 158127)
DTT, 1M (Sigma, D9779)
SDS, 10-20% (Thermo Fisher Scientific, 15525017)
Neb 2.1 buffer (NEB, B7202) or CutSmart Buffer (NEB, B7204S)
MboI, 25 000U/ml (NEB, R0147)
T4-ligase buffer (NEB, B0202)
BSA, 20 mg/ml (NEB, B9000)
ATP, 100mM (Amresco, 0220)
T4-ligase, 5 000U/ml (NEB, M0202)
biotin-14-dATP, 0.4 mM (Thermo Fisher Scientific, 19524016)
dTTP, 10 mM
dGTP, 10 mM
dCTP, 10 mM
Klenow, 5 000U/ml (NEB, M0210)
RNase A, 10 µg/µL (Quagen, 19101)
proteinase K, 20 µg/µL (NEB, P8107)
phenol-chloroform-isoamyl mix solution (Sigma, P2069)
pure 100% ethanol
freshly made 70% ethanol
NaOAc, 3M, pH=5.2 (Thermo Fisher Scientific, R1181)
Tris-HCl, 10mM (Thermo Fisher Scientific, 1862815)
0.1 x TE buffer, pH=8
Qubit™ dsDNA HS Assay Kit (Thermo Fisher Scientific, 32854)
Agencourt® AMPure XP beads (Beckman Coulter, A63880)
Dynabeads, 10mg/ml (Thermo Fisher Scientific, 65601)
Tris-HCl, 0.5M (Thermo Fisher Scientific, 1862815)
EDTA, 0.5M, pH=8.0
Tween 20, 10% (Sigma, P9416)
NEBNext® Ultra™ II DNA Library Prep Kit for Illumina® (NEB, E7645S)
NEBNext® Multiplex Oligos for Illumina®, Index Primers Set 1 (NEB, E7335S)
autoclaved vacuum grease for phenol/chloroform DNA purification
agarose

*EQUIPMENT:*

10 cm plastic Petri dish
filter paper
cotton balls
100 µm nylon mesh

10, 60, 100 μm nylon filters (Millipore)
filter holders (Millipore)
15, 50 mL centrifuge plastic tubes
1.5, 2 mL Eppendorf LoBind tubes
Dounce's homogenizer (2ml)
Magnetic rack
Tube rotator
Thermomixer
PCR thermocycler
Centrifuge with 1.5-2ml and 15-50 mL buckets
Ultrasonicator (M220 COVARIS/E220 COVARIS)
Agilent BioAnanalyzer/TapeStation
Gel electrophoresis equipment
Qubit Fluorometer

#### *Hi-C protocol*

*Prefixing procedures: Day -2 (-3): Blood feeding*
We highly recommend using the colony with 100-150 mature females for producing sufficient amount of mosquito eggs. Blood feed the adult mosquito with fresh (0-5 days old) blood. Second and third blood feedings appear to be more productive in terms of amount of producing eggs.

*Day 0: Egg laying*
Put the egg dish (10-cm-Petri dish with wet cotton balls covered with filter paper (Supplementary Figure 14) accordingly to biological schedule of mosquito strain for overnight egg laying. In the case we used embryos not older than 15 hours because of the tissue differentiation and getting insensitive to bleach).

*Day 1: Embryo collecting and bleaching*
Next day the total amount of eggs must be counted before starting the Hi-C experiment to make sure that there is a sufficient number of eggs for further manipulations. Due to enormous loss of cells at the next protocol steps, not less than 2000-4000 embryos are highly recommended for generating qualitative Hi-C library.

After calculating, carefully collect eggs from the wet filter paper and incubate in presence of 50%-bleach-water solution for 10 minutes with shaking to unseal extraembryonic membranes (amnion and serosa). We used 6- or 12-well plastic plates and 100 μm nylon filters (Millipore) for more convenience. Accurately monitor the time of incubation; depending on the entire structure of covers which should be slightly "spotted" but not overdigested current duration of bleaching may vary but takes in average about 8-12 minutes. We do not suggest modifying bleach concentration because of the damageable effect of high bleach solutions at embryo tissue structure.

*Fixation, cell lysis and restriction*
Freshly prepare and pre-chill all buffer solutions before you start the experiment. Collect eggs after bleaching, remove the leftovers of bleach solution with filter paper, wash 2-3 times with ddH2O, remove the leftovers of any liquid, and add 10 mL fixing buffer with PFA (final concentration 3%). Homogenize the collected eggs with Dounce's or any alternative homogenizer for 2 minutes, filter with 100 μm nylon mesh (Millipore), collect cells and leave for crosslinking for 15 minutes at RT with slow rotation. Filter the cells with nylon mesh less pore-sized (~40-60 μm, Millipore) to decrease the income of cell

aggregates, incubate for additional 10 minutes. Quench PFA by adding appropriate amount of fixing buffer with glycine (20 mL per 10 mL buffer with PFA, total glycine concentration 1M) and incubate on ice for 5 minutes. Collect cells with centrifugation at $4^0$C, 3000 rpm, 15 minutes. Wash with cold 20 mL fixing buffer w/o PFA twice (first, resuspend the pellet into homogenous single-cell suspension in 1 mL of buffer, then add the rest of buffer), wash again with pre-chilled 1.25XNeb 2.1 buffer (buffer specialized for restriction endonuclease of your choice) reducing the reaction volume to 1-1.5 mL (avoid any abrupt temperature shifts), $4^0$C, 4000rpm, 15 min. After discarding the supernatant nuclei pellet should be quickly frozen in liquid nitrogen and stored at $-80^0$C up to several months or resuspended in 300 μL restriction buffer for continuing with Hi-C protocol (*check point – K0, take an ⅕ aliquot for library verification).

| **Reagent** | **Fixing buffer with PFA (10ml):** | | **Fixing buffer w/o PFA (25ml):** | | **Fixing buffer with glycine (20ml):** | |
|---|---|---|---|---|---|---|
| | Adding amount, μL | Final concentration | Adding amount, μL | Final concentration | Adding amount, μL | Final concentration |
| HEPES, 1M | 150 | 15mM | 375 | 15mM | 300 | 15mM |
| KCl, 1M | 600 | 60mM | 1500 | 60mM | 1200 | 60mM |
| NaCl, 1M | 150 | 15mM | 375 | 15mM | 300 | 15mM |
| MgCl, 1M | 40 | 4mM | 100 | 4mM | 80 | 4mM |
| Triton, 10% | 100 | 0.1% | 250 | 0.1% | 200 | 0.1% |
| DTT, 1M | 5 | 0.5mM | 12,5 | 0.5mM | 10 | 0.5mM |
| Protease inhibitor cocktail | 50 | | - | - | - | - |
| PFA, 37%* | 810 | 3% | - | - | - | - |
| $ddH_2O$ | 8,04 mL | - | 22,375 mL | - | 5,4 mL | - |
| Glycine, 2M | - | - | - | - | 12,5 mL | 1,2M |

* Freshly made PFA 37% is highly suggested. Paraformaldehyde does not dissolve but rather can be depolymerized in solution. Depolymerized solutions can be made in water bath with heating to 55-65°C. If necessary, further addition of 1-2 drops of a 1M NaOH solution may be required. According to CSHL protocol for 0,37g PFA powder use 1ml $ddH_2O$ and 14μL 1M NaOH (to enhance PFA solubility), incubate at 60-65°C water bath until solution becomes clear and white PFA flakes completely disappear. Cool the solution to room temperature and use the appropriate amount for preparing fixing buffer.

Take frozen cells aliquot and carefully defrost it on ice for ~10-15 minutes, resuspend in ~300 μL restriction buffer. To degrade DNA-unbound proteins add 10-20% SDS to total concentration 0.3% and incubate at 37°C for 1 hour with shaking, 1000 rpm. Quench SDS by adding 1/5*V 10% triton (~70-80 μL) and incubate at 37°C for 1 hour with shaking, 1000 rpm. After quenching add 150-200U restriction enzyme (MboI or …), accurately mix the reaction by pipetting and incubate overnight (12-15 hours) at 37°C with slowly overhead rotation (for preventing cells sticking), 300-500 rpm.

*Day 2: Biotin-14-dATP marking of DNA ends, ligation*

Next day, once restriction is completed, main solution becomes clearer and more transparent. To inactivate MboI-directed restriction incubate reaction mix at 62°C for 25 minutes, 1000 rpm. During inactivation progress prepare Klenow mix to full-fill the sticky ends with biotin-dATP.

Klenow mix (50 μL):

| **Reagent** | **Adding amount, μL** | **Final concentration** |
|---|---|---|
| biotin-14-dATP, 0.4 mM | 37.5 | 0.3 mM |
| dTTP, 10 mM | 1 | 0.2 mM |
| dGTP, 10mM | 1 | 0.2 mM |
| dCTP, 10 mM | 1 | 0.2 mM |
| Klenow, 5 000U/ml | 8 | 0.8 U/μL |
| ddH2O | 1.5 | - |

Incubate 45-90 minutes at 37°C with rotation, 500-1000 rpm.

While biotin-14-dATP labeling is in progress, prepare ligation mix. Keep T4-ligation buffer, BSA, T4-ligase, and ATP on ice before using.

Ligation mix (1200 μL):

| Reagent | Adding amount, µL | Final concentration |
|---|---|---|
| 10 x T4-ligase buffer* | 120 µL | 1 x |
| Triton, 10% | 100 µL | ~1% |
| BSA, 20 ng/µL | 6 µL | 0.1 ng/µL |
| ddH2O | (1200-x) µL, where x-volume of main reaction | |
| ATP, 100mM * | 12 | 1 mM |
| T4-ligase, 400 000 U/ml | 5 | 2000 U per reaction |
| * Defrost before exploitation only, not refrost and defrost repetitively (both solutions contain ATP-molecules which are extremely sensitive to high temperature). Use aliquots prepared in advance. | | |

Set up a 1 mL-pipette at 1000 µL point and accurately mix the ligation mix by pipetting before adding to the main reaction. Mix the ligation reaction thoroughly by pipetting and incubate at RT (20-25$^0$C) with slow rotation, 300-500 rpm, overnight (15-18 hours). In 2 hours ligation has been started add fresh portion of T4-ligase and ATP to maximize ligation efficiency. Do not allow to appear some sticking cell conglomerates!

*Day 3: Crosslink reversal*

When ligation has been completed, add 7 µL 10 µg/µL RNase A to degrade any present amount of RNA, incubate for 1 hour at 37°C with mixing, 1000rpm. To remove protein-DNA crosslinks add 50 µL proteinase K (20 µg/µL), incubate 1 hour at 56°C with shaking, 1000 rpm. Then, add 0.1V 5M NaCl (V – volume of reaction), incubate at 68°C at least for 2 hours or overnight (do not allow high temperatures for a long period (more than 12-15 hours) because of the risk to cause damages in chromatin structure leading to unqualified Hi-C library).

*Day 4 (or Day 3 - continued): DNA purification, Hi-C library check-in*

Cool the reaction to room temperature and extract DNA by ethanol precipitation. To eliminate the presence of any embryo covers or agglutinated cell conglomerates we suggest using classical phenol-chloroform-isoamyl* purification (or phenol/chloroform) with adding a drop of vacuum grease sterilized by autoclaving. Vacuum grease works as a sponge and adsorb all the rest of embryo covers and allows separating two reaction phases very clear and accurate w/o even any mixing between phases. Preheating phenol-chloroform-isoamyl mix to RT appears to be a crucial step and using chilled/cold solution can lead to a total loss of DNA.

After overnight incubation at 68°C cool main reaction to RT and split into two aliquots for more convenience. Add 1V phenol-chloroform-isoamyl solution, mix thoroughly, centrifuge at max speed for 15 minutes. Carefully transfer the clear upper phase to the new 2 mL LoBind tube (lower phase and middle phase contain all other impurities such as proteins residues, eggs covers, excluding DNA).

Then, continue to perform ethanol DNA extraction according to suggested proportions: add 1.6V pure ethanol and 0.1V 3M NaOAc, incubate for 1-2 minutes at RT with rotation, transfer tubes at -80°C and keep for 15 minutes (or until the liquid becomes frozen), centrifuge at 4°C at max speed for 15 minutes. Visible white/light yellow pellet should precipitate at the tube bottom. Transfer tubes on ice and leave for a couple minutes before discarding the supernatant. Carefully wash DNA pellet twice with pre-chilled 800 µL 70-80% ethanol, centrifuge at RT, max speed. Accurately mix the pellet each time avoiding cells sticking to the plastic tips and tubes possibly. The actual size of pellet should become less after second washing because of diluting DTT and ATP particles at RT. After washing steps perform air-drying for 5 minutes, then dissolve the pellet with 130µL 10mM Tris-HCl, incubate at 37°C for 15 minutes for better efficiency, and split the solution into 10µL (for Qubit, 2µL+PCR 1µL+ 5µL gel electrophoresis) and XµL aliquots (for COVARIS, depending on instrument technical requirements). Keep samples at -20°C before next steps.

Perform High Sensitive Qubit assay to estimate current DNA concentration using recommended protocol and check the library quality by PCR-analysis with specified 3C primers. 10-200ng/µL range (1.3-25µg in total) is optimal and sufficient quantity to continue with Hi-C protocol.

*Day 5 (or Day 4): DNA shearing + size selection*
After verifying DNA concentration of HiC library continue with ultrasonic DNA shearing. The optimal COVARIS requirements should be chosen based on BioAnalyzer/TapeStation or regular gel electrophoresis results given that the majority of DNA fragments have to be performed as a smear between 300-500 bp on electrophoregram.

Accurately transfer sheared DNA to the fresh tube, wash COVARIS tube with ddH2O, increase the reaction volume to exactly 200 µL, and perform 300-500 bp DNA fragments purification and size selection with Agencourt® AMPure XP beads following to recommended protocol.

Pre-warm AMPure XP beads to RT, mix thoroughly before using, add exactly 0.55X volumes (110 µL) to the reaction, mix by pipetting, and incubate at RT for 5 minutes. Keep the liquid containing DNA fragments shorter than 500 bp by separating beads at magnetic rack for 2-5 minutes. Transfer the clear solution to the new tube. To purify 300-500 bp DNA fragments from RNA and short fragments add new portion of AMPure beads - 0.1X (30 µL) – to the main reaction, mix thoroughly, and incubate for 5 minutes at RT. Separate beads and liquid at magnetic rack, keep the beads (!). Wash the beads with freshly made 70%-ethanol two times without agitation, keeping the beads on magnet all the time. Evaporate the small amount of ethanol by air-dry for 3-7 minutes, but be careful and avoid overdrying – beads should have dark brown color without any visible drops of ethanol, light color of pellet and small cracks evidence for overdrying which can lead to decreasing in DNA yield. Dissolve the pellet in 312 µL 10 mM Tris-HCl, TE buffer, or regular ddH2O. Estimate DNA concentration by dsDNA High Sensitivity Qubit Assay and run 10 µL of purified solution on 2% agarose gel to verify the proper size of selected DNA fragments. Keep the tube at -20°C or continue with the next step.

*Day 6 (or Day 5): biotin pull down + library preparation for Next Generation Illumina sequencing*

We suggest to aliquot the beads mix for single use samples to prevent extra heating each time. Pre-heat Dynabeads to RT. Thoroughly wash 150 μL of 10mg/μL dynabeads with 400 μL 1 x Tween Washing Buffer (5mM Tris-HCl, 0.5mM EDTA, 1M NaCl, 0.05% Tween 20), separate on magnetic rack. Resuspend the beads in 300 μL 2 x Binding Buffer (10mM Tris-HCl, 1mM EDTA, 2M NaCl) by pipetting and add to the main reaction. Mix thoroughly and incubate at RT for 15 minutes with slow mixing to enhance binding between biotinylated DNA and streptavidin beads. After incubation separate dynabeads on magnet for 5 minutes. More slowly and "sharp-edged" beads movements can be expected because of moving beads with streptavidin-biotin-DNA complexes in a different manner. Discard the supernatant, wash the beads twice with Tween Washing Buffer with 2-minutes incubation at 55°C each time, then one time wash with 150 μL 10mM Tris-HCl buffer without heating. Do not mix the beads during the washing steps! Separate the beads on a magnetic rack (discard supernatant, keep the beads)! After the last washing add 50 μL 10 mM Tris-HCl buffer, resuspend the mixture thoroughly, transfer to the PCR tube, and continue with NEBNext® Ultra™ II DNA Library Prep Kit for Illumina® and NEBNext® Multiplex Oligos for Illumina® Index Primers Set 1 to amplify the library directly on streptavidin beads.

Follow the recommended protocol instructions and index compatibility recommendations provided by NEB for NEBNext® Ultra™ II DNA Library Prep Kit for Illumina® with some exceptions because of the case of amplification directly on beads.
We strongly suggest not make any stops and additional defrosting/frosting steps to maximize the reaction efficiency.

Defrost all the reagents on ice, mix thoroughly, and add the following components to the new nuclease-free PCR tube:

| **Reagent** | **Adding amount** |
|---|---|
| NEBNext Ultra II End Prep Enzyme Mix (green top) | 3 μL |
| NEBNext Ultra II End Prep Reaction Buffer (green top) | 7 μL |
| Hi-C library diluted in 10 mM Tris-HCl | ~50 μL (will be over because of the beads volume) |

Mix entire reaction by pipetting up and down 10-20 times

Mix entire reaction by pipetting up and down 10-20 times without extensive bubbling, perform quick spin, but make sure then that dynabeads are spread through the tube evenly not laying at the bottom. Place in the thermocycler with ≥75°C heat lid setting, incubate 15 minutes at 20$^{0}$C, mix the beads directly at thermocycler, and incubate at 20°C for 15 minutes more. Thoroughly mix the beads avoiding bubbling, incubate at 65°C for

15 minutes, mix directly at thermocycler, and incubate for 15 minutes more. Hold at 4°C before continuance with the next steps.

Defrost on ice the following reagents, add to the main reaction (~60-65 µL), and mix thoroughly 10-20 times by pipetting:

| Reagent | Adding amount |
|---|---|
| NEBNext Ultra II Ligation Master Mix (red top) | 30 µL |
| NEBNext Ultra II Ligation Enhancer (red top) | 1 µL |
| NEBNext Ultra II Adaptor for Illumina | 2.5 µL |

Total reaction volume can differ because of the volume covered by beads but should be in range ~95-100 µL in total. Perform a quick spin to prevent any on-tube-side-left drops but then make sure that beads are spread across the entire reaction volume evenly. Incubate at 20°C (heated lid off) for 20 minutes with intermediate pipetting the beads. Add 3 µL USER Enzyme (red top), mix well, and incubate at 37°C for 15 minutes with the lid setting heated to ≥47$^0$C. After incubation transfer the reaction mixture to LoBinding 1.5 mL tube and perform cleanup from extensive amount of adaptor. Discard the liquid containing adaptors by separation the beads on magnetic rack, wash twice with 600 µL Tween Washing Buffer with 2-minutes heating at 55$^0$C, separate beads for at least 5 minutes. Wash the beads with 100 µL 10 mM Tris-HCl, separate on magnet, discard the liquid phase, and add 10 µL 10 mM Tris-HCl to perform PCR.

We would suggest splitting the total reaction volume in two aliquots and performing PCR for one half of the Hi-C library to verify the optimal quantity of PCR cycles based on the final amount of DNA. Usually, 9-10 cycles are sufficient but in case of some DNA loss during the washing or pull-down steps it may be necessary to increase the number of cycles to 11-12.

Defrost the following reagents and add to the sterile PCR tube to total volume of 25 µL:

| Reagent | Adding amount |
|---|---|
| Adaptor ligated DNA fragments on the Dynabeads | 7.5 µL |
| NEBNext Ultra II Q5 Master Mix (blue top) | 12.5 µL |
| Universal PCR Primer (blue top) | 2.5 µL |

| | |
|---|---|
| Index primer (blue top) | 2.5 µL |

Perform PCR according to the following conditions:

| Cycle step | Temp | Time | Cycles |
|---|---|---|---|
| Initial denaturation | 98°C | 30 seconds | 1 |
| Denaturation | 98°C | 10 seconds | 10 |
| Annealing/Extension | 65°C | 75 seconds | |
| Final extension | 65°C | 5 minutes | 1 |
| Hold | 4°C | ∞ | |

Mix the reaction directly into thermocycler after 3rd and 6th cycles to allow the amplification even from “the hidden” beads. After PCR is completed increase the reaction volume to 125 µL by adding 10 mM Tris-HCl or ddH2O, separate the beads on magnetic rack for 5 minutes, and keep the liquid discarding the beads. Transfer supernatant to the new LoBind tube and perform AMPure beads cleanup following to recommended instructions. To select DNA fragments upper than 400 bp add exactly 0.7 volume of pre-heated to RT AMPure beads, pipette thoroughly, incubate for 5 minutes at RT, separate on magnet, and keep the beads! Wash twice with 70-80% freshly diluted ethanol excluding any agitation. Remove any last drops of ethanol, perform air-drying for 3-5 minutes at RT until AMPure beads pellet becomes slightly light and wrinkled. Do not allow overdrying which can affect on solubility of DNA. Dissolve Hi-C library in 30 µL 0.1 x TE buffer and measure the current concentration by High Sensitivity dsDNA Quibit. Total amount of DNA should be higher than at least 15 mM but should be checked more precisely in accordance to MiSeq/HiSeq specific requirements. If the total amount of DNA after PCR is sufficient for MiSeq+HiSeq experiments split the solution between two tubes and store at -20°C until sequencing. If the producing amount of DNA is not enough – perform PCR increasing the number of cycles to 11-12. Verify the concentration by High Sensitivity Qubit dsDNA assay. Use the best PCR products – less cycles and more DNA - for next generation sequencing.

#### Supplementary Protocol II

*REAGENTS:*
anti-trimethyl-histone H3 (Lys27) (Millipore, #07-449) antibody
agarose beads (Pierce™ Protein A/G Agarose, ThermoFisher #20421)
autoclaved ddH2O
freshly made 50%-bleach/water solution
NaCl, 1M
MgCl, 1M
KCl, 1M
HEPES, 1M (Sigma, H0887)
Glycine, 2M (Sigma, G7126)
Triton, 10% (Sigma, T8787)
Protease Inhibitor Cocktail (Sigma, P8340)
Freshly made PFA, 37% (Sigma, 158127)
DTT, 1M (Sigma, D9779)
sodium deoxycholate (Sigma Aldrich, 30970)
N-Lauroylsarcosine sodium solution (Sigma Aldrich, 61747)
SDS, 10-20% (Thermo Fisher Scientific, 15525017)
EDTA, 0.5M, pH=8.0
EGTA
Tris-HCl, 0.5M, (Thermo Fisher Scientific, 1862815)
LiCl, 1M
NP-40
$NaHCO_3$
BSA, 20 mg/ml (NEB, B9000)
RNase A, 10 μg/μL (Quagen, 19101)
proteinase K, 20 μg/μL (NEB, P8107)
phenol-chloroform-isoamyl mix solution (Sigma, P2069)
pure 100% ethanol
freshly made 70% ethanol
NaOAc, 3M, pH=5.2 (Thermo Fisher Scientific, R1181)
0.1 x TE buffer, pH=8
Qubit™ dsDNA HS Assay Kit (Thermo Fisher Scientific, 32854)
Agencourt® AMPure XP beads (Beckman Coulter, A63880)
Tris-HCl, 0.5M (Thermo Fisher Scientific, 1862815)

NEBNext® Ultra™ II DNA Library Prep Kit for Illumina® (NEB, E7645S)
NEBNext® Multiplex Oligos for Illumina®, Index Primers Set 1 (NEB, E7335S)
autoclaved vacuum grease for phenol/chloroform DNA purification
agarose

*EQUIPMENT:*

100 μm nylon mesh
10, 60, 100 μm nylon filters (Millipore)
filter holders (Millipore)
15 mLcentrifuge plastic tubes
1.5, 2 mL Eppendorf LoBind tubes
Dounce's homogenizer (2ml)
Tube rotator
Thermomixer

PCR thermocycler
Centrifuge with 1.5-2ml
Bioruptor Dianogene
Agilent BioAnanalyzer/TapeStation
Gel electrophoresis equipment
Qubit Fluorometer

#### *ChIP-seq experiment*

~1000-2000 mosquito eggs at 15-18 hours after oviposition were used to prepare ChIP-seq library. Embryos were bleached in 50% bleach for 5-10 minutes, homogenized in 1.8% formaldehyde/buffer A (60 mM KCl, 15 mM NaCl, 15 mM HEPES, 4 mM $MgCl_2$, 0.05% triton, 0.5 mM DTT, protease inhibitors) and filtered through the 60 and 10 pore-sized filter consequently to remove giant cell clusters. Formaldehyde was quenched by adding glycine to final 225 mM. 2-3 washing steps in buffer A were performed to remove formaldehyde residues. Then, samples were snap frozen in liquid nitrogen until in use.

Cell and nuclear lysis: de-freeze nuclear pellet on ice for 20-30 minutes, add 1 mL of lysis buffer (140 mM NaCl, 15 mM HEPES, 1 mM EDTA, 0.5 mM EGTA, 1% triton, 0.5 mM DTT, 0.1% sodium deoxycholate, protease inhibitors), and incubate on rotation wheel for 10 minutes at RT. Centrifuge for 5 minutes at 4000 rpm at $4^0$C. Add 0.5 mL lysis buffer, 5 µL 10% SDS, 5-8 µL 30% N-Lauroylsarcosine sodium solution, and incubate for 10-15 minutes.

Perform sonication using the following shearing conditions on Bioruptor Dianogene: 8-10 cycles of 10/10sec ON/OFF in lysis buffer with SDS and NP40.

After sonication, add 0.5 mL lysis buffer or IP dilution buffer (167 mM NaCl, 16.7 mM Tris-HCl, 1.2 mM EDTA, 0.01% SDS, 1.1% triton, protease inhibitors), pellet the chromatin by centrifugation for 20 minutes at maximum speed at 4C. Aliquot 1/10 volume to check the quality and quantity of sheared chromatin. Measure the concentration and use ~10 µg of sheared chromatin for each ChIP-seq experiment. Approximately, 1000 eggs produce ~10-20 µg of chromatin. We have used *Drosophila melanogaster Canton-S* $3^{rd}$-stage larvae chromatin as a spike-in*. The spike-in quantity was approximately 5-10% of start material (~0.5-1 µg)

*Spike-in: 8 *Drosophila melanogaster Canton-S* $3^{rd}$-stage larvae were crushed with scalpel and homogenized in buffer A with 1.8% formaldehyde for 10 min. Then, we performed sonication following the same protocol as for mosquito eggs.

All next steps should be performed on ice or at 4°C.

Dilute sheared chromatin solution at least 5 times in IP dilution buffer to proceed with IP (such as for 200 µL of sonicated chromatin allowing the reaction volume to be 1 mL).

Prepare agarose beads: 50 µL for pre-clearing step and 50 µL for ChIP experiment. For 1V of beads use 10V of a buffer solution during each washing step. Wash 3 times in PBS, then 1 time in IP dilution buffer. After washing, dilute the beads in 100 µL IP dilution buffer to obtain a working aliquot of agarose beads mix per one experiment.

Chromatin pre-clearing step: mix ~10 µg of chromatin with 50 µL previously washed agarose beads. Incubate overnight at 4°C with slow rotation. Next day transfer the liquid to a fresh tube throwing the beads and keep 10% of chromatin solution as ChIP-seq input. Store the input at -20°C before proceeding with ChIP DNA precipitation.

The second part of prepared beads (100 µL of working aliquot) mix with 0.1% BSA solution in PBS and incubate for 1-2 hours at 4°C with rotation to block nonspecific interactions. Then, wash 3 times 1xPBS and 1 time in IP dilution buffer. Combine pre-cleared beads with target antibody (Millipore, #07-449, 5-10 µg) diluted in 1 mL IP dilution buffer and incubate overnight at 4°C with slow rotation. Next day wash the beads-antibody complexes in 1xPBS 3 times and in IP dilution buffer 1 time. Dissolve the beads in a total volume of 100 µL per ChIP experiment.

Combine pre-cleared chromatin with agarose beads-antibody complex and incubate overnight at 4°C with slow rotation.

Next day thoroughly wash the agarose beads in a series of buffer solutions: 3 times in low salt buffer (LB: 150 mM NaCl, 20 mM Tris-HCl, 2 mM EDTA, 1% triton, 0.1% SDS), 2 times in high salt buffer (HB: 500 mM NaCl, 20 mM Tris-HCl, 2 mM EDTA, 1% triton, 0.1% SDS), 2 times in LiCl buffer (LiB: 0.25 M LiCl, 10 mM Tris-HCl, 1 mM EDTA, 1% NP-40), 1 times in TE buffer. Incubate samples for 10 minutes at 4°C with rotation during each washing step, then spin for 3 minutes at 1000 rpm, 4°C. Elute DNA in 250 µL elution buffer (EB: 1% SDS, 0.1 mM $NaHCO_3$) by incubation at 65°C for 10 minutes. Repeat elution to increase the yield.

De-freeze the input samples and proceed to crosslink reversion. Incubate input and experimental chromatin samples at 65°C overnight in presence of 0.25M NaCl, 10 mM EDTA and 40 mM TrisHCl.

Next day add 50 µL proteinase K to the samples, incubate for 2h at 55°C. Extract the DNA with phenol-chloroform (or specified cleaning columns) followed by ethanol precipitation. Dissolve the DNA pellet in 55 µL H2O or purify DNA using AMPure beads according to manufacturer recommendations.

Proceed with ChIP-seq library preparation for the Illumina sequencing with NEBNext® Ultra™ II DNA Library Prep Kit for Illumina® (NEB, E7645S).
